# The Genome-Wide Effect of Drift and Selection over a Single Generation

**DOI:** 10.64898/2026.08.04.742829

**Authors:** Gabriele Sgarlata, Graham Coop

## Abstract

The relative importance of genetic drift *versus* selection to evolutionary change has long been debated. This debate has mainly focused over long-time-scales (e.g. hundreds of thousands of generations), leaving the question of short-term evolutionary change relatively unaddressed. Our knowledge about the effects of selection on genetic change over short time scales is often based on identifying major allele frequency changes at few loci with large selective advantage. Yet selection often acts on polygenic traits where the short-term response is shaped by small shifts in allele frequency at many loci that will be difficult to distinguish from genetic drift. Here, we quantify the genome-wide effects of polygenic selection over a single generation, using the idea that alleles in stronger genetic correlation (LD) with selected alleles are expected to show greater variance in allele frequency change than expected under genetic drift. We derive expressions relating variation in LD among loci to the variance in allele frequency change due to linked selection and genetic drift and leverage this theory to quantify the contribution of linked selection to a single generation of allele frequency change. To demonstrate our approach, we decompose the genome-wide allele frequency change in the UK Biobank using fitness proxy phenotypes. We show that selection makes a small, but significant, contribution, with genetic drift making up the large majority of the change in allele frequencies. Our framework could be applied to other organisms for which data on number of offspring or allele frequencies over consecutive generations are available, enabling investigations of the short-term, genome-wide effects of polygenic selection across a wide range of species.

## Introduction

The debate over the contribution of genetic drift and selection to evolutionary change traces back to the inception of the modern evolutionary synthesis (Fisher, 1930; Wright, 1931; Provine, 1989; Dietrich, 2006). Genetic drift generates changes in allele frequencies because of chance correlations between individuals’ genotypes and their reproductive success, as well as the randomness of Mendelian transmission. While selection generates allele frequency change because the genotype of an individual covaries with their fitness (the expected reproductive success given their genotype, Price, 1970; Robertson, 1966, 1968), this selective covariance can arise because the allele at a locus directly affects fitness but also because of its correlation to other alleles influencing fitness, e.g. on the same chromosome (correlated selection or linked selection, Maynard Smith AND Haigh, 1974; Charlesworth *et al*., 1995; Nordborg *et al*., 1996; Barton, 2000). We have good evidence of the long-term impact of correlated selection on genome-wide levels of polymorphism through patterns of diversity across many species (Begun and Aquadro, 1992; Sella *et al*., 2009; Cutter and Payseur, 2013; Corbett-Detig *et al*., 2015; Coop, 2016; Charlesworth and Jensen, 2021). In humans, linked selection has had a moderate impact on long-term patterns of polymorphism (Mcvicker *et al*., 2009; Cai *et al*., 2009; Hernandez *et al*., 2011; Murphy *et al*., 2022).

Over shorter time-scales we have many good examples of selection on large effect alleles and phenotypes (Dobzhansky, 1943; Fisher and Ford, 1947; Hendry and Kinnison, 1999; Kreiner *et al*., 2022). In humans, in particular, a number of cases of selection driven change in ancient DNA at individual loci and on polygenic scores have been documented (Mathieson and Terhorst, 2022; Irving-Pease *et al*., 2024; Akbari *et al*., 2026). We also have examples of selection driving genetic change within a single human generation (Allison, 1964; Mostafavi *et al*., 2017; Sanjak *et al*., 2018; Mathieson *et al*., 2023). Estimating the short-term, genome-wide effects of selection is challenging, however, because fitness is often expected to be highly polygenic and so selection generates small changes in frequency at many loci leaving a very diffuse signal.

In humans, current day patterns of reproductive success are influenced by many cultural, economic, and social factors, and also reflect more biological factors such as the timing of puberty, reproductive aging, and fertility (Balbo *et al*., 2013; Sear *et al*., 2016). Variation in human reproductive success shows low but significant heritability (Kirk *et al*., 2001; Byars *et al*., 2010; Zietsch *et al*., 2014; Mills and Tropf, 2024), as is the case in many species (Bonnet *et al*., 2022). Genome-wide association studies (GWAS) have shown that most fitness-related traits, such as longevity and number of children, are highly polygenic and subject to various forms of confounding (Barban *et al*., 2016; Pilling *et al*., 2016; Joshi *et al*., 2017; Howe *et al*., 2022; Mathieson *et al*., 2023; Tan *et al*., 2024). Thus, the impact of selection will often be very weak on individual alleles but spread over many loci. Conversely, human populations are currently very large and so the effects of drift per generation are also expected to be very small. As such, it is an open question how selection and drift trade off in modern humans.

Here we propose a method for estimating the relative effects of selection and drift on allele frequency change in a single generation. We illustrate this approach in humans using genome-wide association studies of fitness proxies in the UK biobank. We find that linked selection makes a small but measurable impact on genome-wide allele frequency change.

## Results and Discussion

We aim to develop simple approximations for the contributions of selection and drift in terms of quantities that we can estimate empirically. To do so, we begin by imagining a set of *N*_0_ diploid individuals and following these individuals from birth until they themselves have completed their reproductive lives. The *i*^*th*^ individual contributes *k*_*i*_ children (*k*_*i*_ ≥ 0) to the next generation and has genotype *x*_*i*_ ∈ *{*0, 1, 2*}*, which represents the number of copies of a particular allele at the focal SNP *x*. The allele frequency change (Δ*p*_*x*_) at the focal site to the next generation, formed by their offspring, is (Eq. S1-S4):

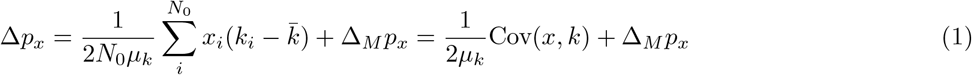

where *µ*_*k*_ = 2*N*_1_*/N*_0_ is a population size scaling factor between generations, which corresponds to the mean absolute fitness of the parental population. Intuitively, changes in allele frequency in the next generation arise from the covariance between the parental genotypes and offspring number, and the randomness of Mendelian segregation from heterozygote parents (Δ_*M*_*p*_*x*_).

Next, we decompose the number of offspring *k*_*i*_ into its expectation given the individual’s genotype and the stochastic deviation from this expectation; namely, *k*_*i*_ = *µ*_*k*_*f*_*i*_ + *d*_*i*_ (following a quantitative genetics approach to decomposing change, see e.g. Santiago and Caballero, 1998). The term *µ*_*k*_*f*_*i*_ gives the expected number of offspring given an individual’s genotype and that of their partner(s). While, the deviation *d*_*i*_ arises from stochastic events and environmental factors, and has a mean of zero across individuals 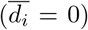. Then the allele frequency change can be written as

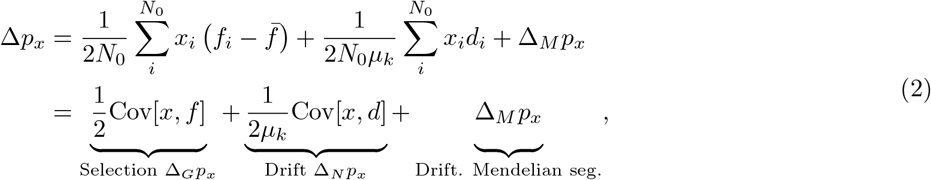

and the expected squared allele frequency change as

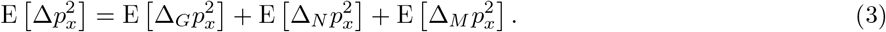

Thus, the expected change in allele frequency can be viewed as arising from three components, which we assume to be uncorrelated (see SI Appendix S3 for a relaxation of this assumption): i) the covariance between the genotype at a focal SNP and the fitness of individuals (Δ_*G*_*p*_*x*_, Robertson, 1966; Price, 1970), ii) the covariance between the genotype at a focal SNP and environmental deviation (Δ_*N*_*p*_*x*_) and iii) the randomness of Mendelian segregation (Δ_*M*_*p*_*x*_) (see e.g. Buffalo and Coop, 2019). The latter two terms correspond to the contribution of genetic drift, noting that here genetic drift includes both sampling noise from parents to offspring and correlations between genotypes and environmental causes of variation in reproductive success.

We assume a polygenic additive model for relative fitness with *L* biallelic loci, such that 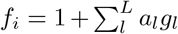, where *a*_*l*_ is the effect size and *g*_*l*_ ∈ *{*0, 1, 2*}* is the genotype at the selected locus *l*. Then 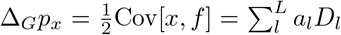 where *D*_*l*_ is the covariance (linkage disequilibrium, LD) between the genotype at the focal site *x* and the selected locus *l* (Eq. S11). For simplicity, we begin by assuming no systematic signed covariance between selected loci (E(*a*_*l*_*D*_*l*_*a*_*l*_*′ D*_*l*_*′*) = 0, for *l* ≠ *l*′) due to previous generations of selection (Bulmer effect, Bulmer, 1974; Walsh and Lynch, 2018) or assortative mating; however, our full model includes these effects (SI Appendix S2.3). Under this model and simplifying assumption, we can write the expected squared change in allele frequency due to polygenic linked selection as

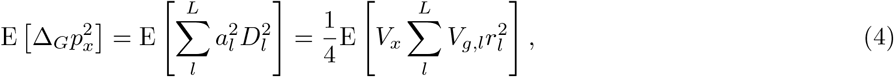

where *V*_*x*_ is the focal site expected heterozygosity, 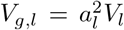 is the contribution of the *l* locus to the additive genic variance for fitness 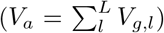, and 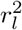 is the squared genotype correlation coefficient between our focal locus *x* and the selected locus *l* (SI Appendix S1.1, Buffalo and Coop, 2019). Thus, the effect of polygenic linked selection on the squared allele frequency change at a site is a weighted sum over its LD to all selected sites, where the weights are the contributions of the selected loci to the additive genic variance (Fig. 1A). Then, we can write the expected squared allele frequency change at a site as

**Figure 1:**
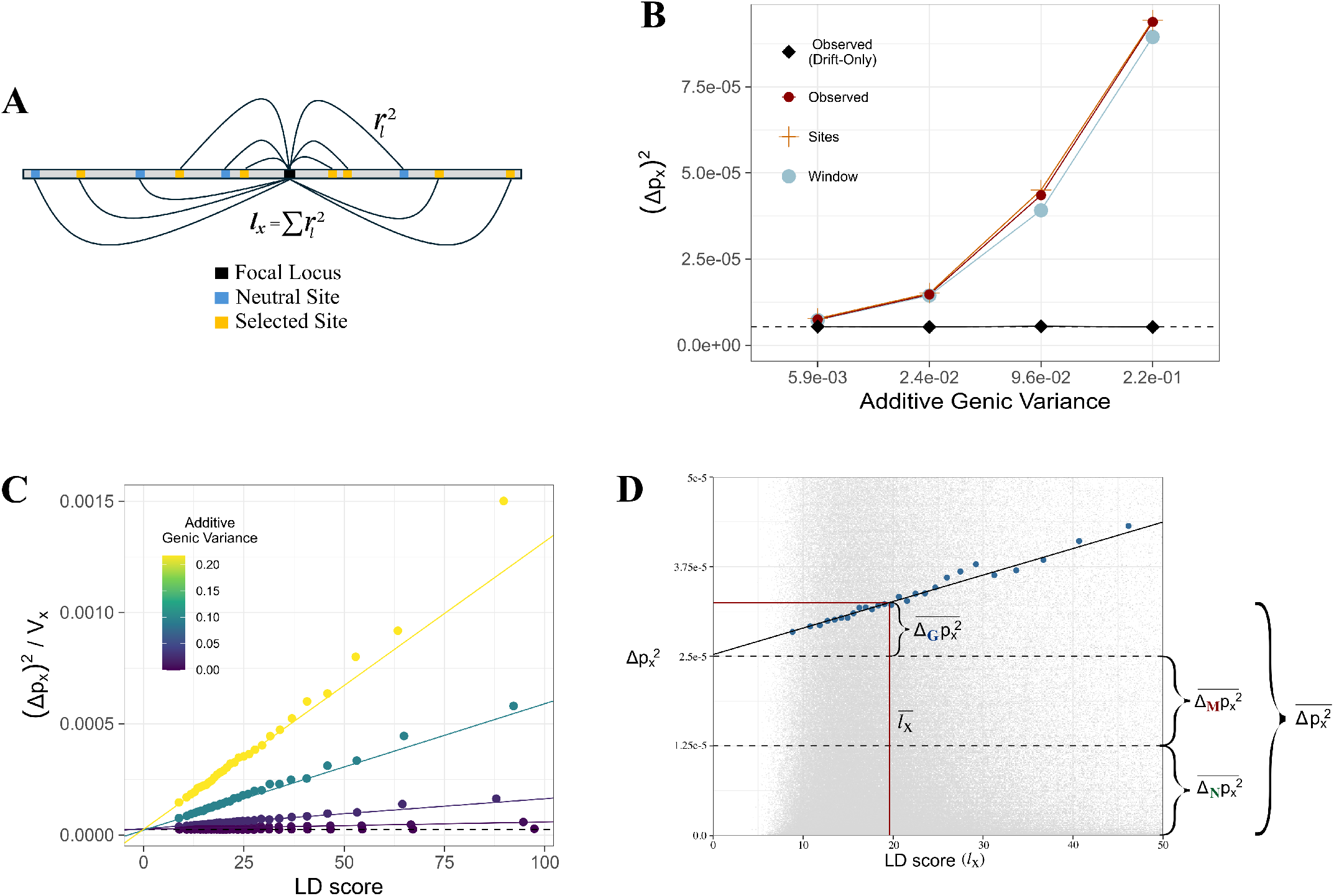
Predicting allele frequency change over a single generation varying the additive genic variance. **A)** Schematic representation of LD score calculation for a focal site *x*. **B)** The average squared allele frequency change over a single generation with predictions using the LD score of only the selected sites weighted by the true effect sizes (“Sites” Eq. 6) or using the LD score over neutral and selected sites multiplied by the average additive genetic variance (“Window” Eq. S28). **C)** Directional selection simulations show that increasing the additive genic variance for reproductive success increased the slope of the LD score regression for the squared allele frequency change. The colored points show the average calculated in 2.5% quantiles of the LD score predictor. **D)** Graphical representation of the decomposition of the squared allele frequency change based on LD score regression. Each grey dot is a single SNP’s estimated squared allele frequency change, normalized by *V*_*x*_. All simulations are based on the recombination map of chromosome 22 scaled on a simulated chromosome of 5 Mbp, where an additive trait (∼ 5, 000 loci) is under selection for a single generation. The points show the mean in 5% bins of LD score. Lighter colours indicate simulations with higher genetic variation for the fitness associated trait. To change the additive genic variance we change the variance of the effect sizes of the alleles contributing to trait variation.

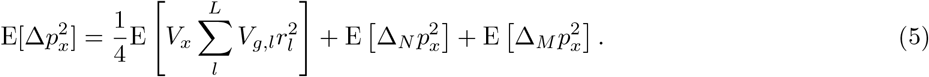

Simulations show that Eq. 5 and S28 accurately predict the average squared allele frequency change in a panmictic population that undergoes a single generation of selection on a highly polygenic trait (“Sites” in Fig. 1B).

We can then approximate Eq. 5 in terms that we can estimate from data. We do not know which loci are selected and what their additive genic variance is, but Eq. 5 can be modified to average the additive genic variance across all loci within some sufficiently large window (*L*_*W*_) around a focal locus:

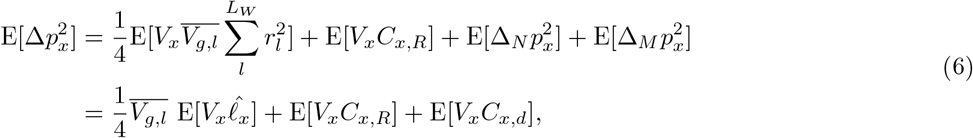

where 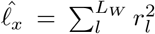 is the LD score at the focal locus *x* summed over the *L*_*W*_ loci in a surrounding window (Bulik-Sullivan *et al*., 2015). The term *V*_*x*_*C*_*x,R*_ represents an error term from the contribution of polygenic linked selection that is not well captured by averaging the additive genic variance across all loci. The term *V*_*x*_*C*_*x,d*_ captures the fact that the neutral components of the squared allele frequency change also scale with the expected genotypic variance at a locus (SI Appendix S1.2, S1.3). The approximation in Eq. 6 predicts that we should expect a larger squared allele frequency change for SNPs with higher LD score, as on average they are correlated with more selected loci. This prediction holds well in simulations (Fig. 1C), which show a steeper linear slope under simulations with a higher additive variance for fitness.

The approximation based on Eq. 6 performs well in estimating the squared allele frequency change due to selection (Fig 1B), with a slight underestimation due to the selection error term *C*_*x,R*_. This approximation also assumes that there is no correlation between the additive genic variance of a selected locus (*V*_*g,l*_) and its squared LD to the focal locus 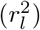, which will be violated when selection acts consistently over long time periods. However, the approximation can be improved by stratifying the squared genotype correlation at a focal site 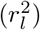 by the minor allele frequency (MAF) of the loci within the window *L*_*W*_ (Finucane *et al*., 2015), allowing each MAF bin to have its own contribution to the total additive genic variance (Eq. 12; Fig. S1).

Importantly, these considerations suggest a simple approximation that is readily relatable to data. Namely, given estimates of allele frequency change across a generation, we can fit the linear model

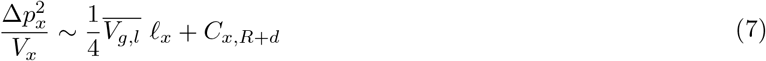

using a large number of loci (*x*) spread throughout the genome to estimate 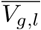. This is mathematically equivalent to LD score regression, which has been widely adopted to estimate SNP-heritabilities and genetic correlations from GWAS summary statistics (Bulik-Sullivan *et al*., 2015, 015b). The slope of LD score regression is used to estimate the genetic contribution of a phenotype, while the intercept absorbs the GWAS sampling noise and environmental stratification (although this separation can fail, for example due to strong or inhomogeneous stratification; De Vlaming *et al*., 2017; Berg *et al*., 2019, see Appendix S2.2, Eq. S40, S41).

The slope and intercept of this linear regression allow us to decompose the average squared allele frequency change (red horizontal line in Figure 1D). The contribution of selection 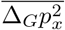 is estimated by the height of the average point above the intercept 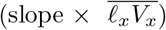. The contribution of drift is estimated by equating the intercept of the regression with the contributions due to non-genetic variance in offspring number and Mendelian segregation 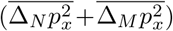. This approach is likely to underestimate the contribution of selection to changes in allele frequency as the intercept can be inflated above its neutral expectation by selective factors not captured by the LD score on common variants (*C*_*x,R*_). To better account for this, we can include other genomic predictors in the regression, for example local recombination rate or a proxy for the density of selected sites.

Our simulations above are based on a model of directional selection on a trait; however, because our model simply relies on LD to a polygenic set of selected alleles, it captures different modes of selection. For example, under simulations of stabilizing selection on a highly polygenic trait, higher additive genic variance leads to steeper slopes in the LD score regression and so greater changes in allele frequency due to selection (Fig. 2A). Selection on traits that are undergoing assortative mating represents a strong violation of our basic model (Eq. 4, SI Appendix S2.3), as trait-increasing alleles are brought together into genome-wide LD through assortative mating. This violation means that the slope of the LD score regression can no longer be interpreted as the additive genic variance for the trait (Border *et al*., 022a; Veller *et al*., 2024, SI Appendix S2.3). However, even under assortative mating the contribution of selection to allele frequency change can still be directly quantified by the slope (SI Appendix S2.3), with positive assortative mating acting to amplify selection driven allele frequency change (see Fig. 2).

**Figure 2:**
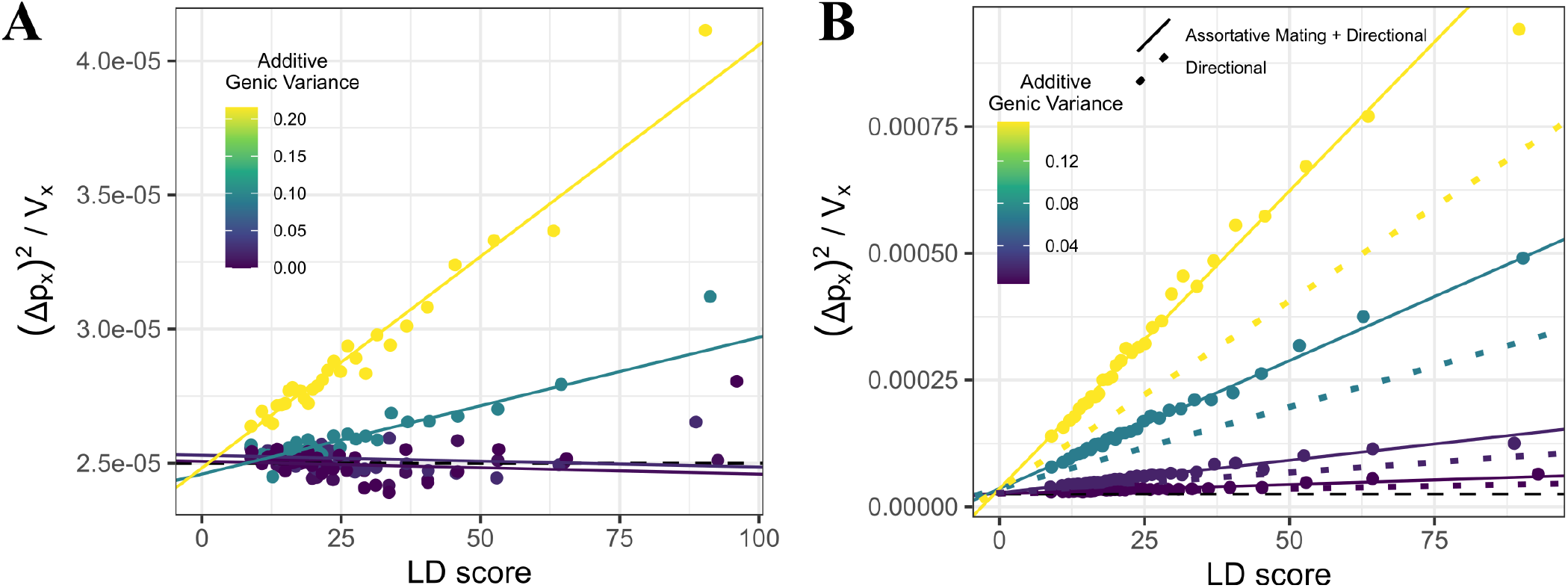
LD score regression on squared allele frequency change under other models of selection. Horizontal dashed lines show the expectation under genetic drift only. The simulation details of the trait and genome match Figure 1. **A)** Simulations under stabilizing selection. **B)** Simulations under assortative mating and directional selection. In the assortative mating simulations the population undergoes positive assortative mating on the phenotype *f*, with a correlation coefficient of 0.4 between mates for 10 generations with no selection. Following this period of assortative mating there is a single generation of selection and the dots and solid line show the resulting relationship between LD score and the squared allele frequency change. The dotted lines shows the expectation under directional selection with no assortative mating, with matching per site additive genic variance.

### Allele frequency change in the UK biobank

To apply this approach to empirical data, we turned to the UK Biobank. While the UK Biobank does not include multigenerational data for most participants, we can predict the squared allele frequency change needed for Eq. 7 by performing GWAS on a proxy for fitness (Fig. S2), namely the self-reported phenotypes “number of live births” for females and “number of children fathered” for males. We will interpret these as the lifetime reproductive successes (*k*) of individuals, but we note that the people in the UK Biobank are sampled as adults, thus our results omit the contribution of early life viability effects. We can express the squared allele frequency change, scaled by the observed genotypic variance *V*_*x*_, by rewriting Eq. 1 as

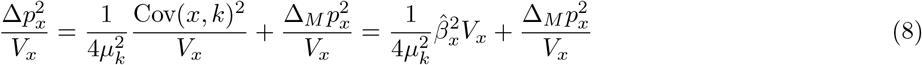

where 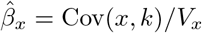 is the estimated effect size at locus *x* from a GWAS on the number of offspring *k*, which measures the contribution of variance in lifetime reproductive success to the squared allele frequency change at a locus. We can then perform LD score regression on the GWAS effect sizes to estimate the contribution of polygenic linked selection to the squared allele frequency change. Without the genotypes of the children, we do not get to observe the outcome of Mendelian segregation. However, 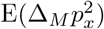 can be accurately calculated for each SNP as this follows from binomial sampling given its observed level of heterozygosity (Eq. S21).

We restricted our GWAS analysis to the subset of UK Biobank individuals designated as ‘White British’ (WB) to ensure relatively little population stratification, thus reducing the amount of environmental confounding in the GWAS (151, 675 males and 175, 228 females, see Methods). GWASs were conducted separately in males and females, so our initial allele frequency change calculations will be for males and females separately. We did not include any principal components to correct for population structure in the GWAS (e.g., principal components). However, when we do so, the results change only slightly (Fig. S3-S7). The squared allele frequency change averaged across loci is very small (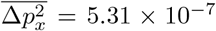 for female “number of live births”), corresponding to very slight shifts in allele frequency (on average ∼ 7 / 10, 000), with ∼ 50% of this change coming from Mendelian segregation out of heterozygote parents alone 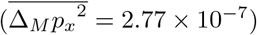.

We performed LD score regression on the predicted squared allele frequency change in the WB cohort (Fig. 3) for common alleles (MAF *>* 5%). The impact of selection at a locus will also include associations to rare selected alleles. Thus, in this regression, we also included the ‘B-value’ at each focal site, a measure of the impact of linked selection (background selection) on the genetic diversity of a site that accounts for both the local recombination rate and the density of functional sites (Murphy *et al*., 2022), and the log_*e*_ of recombination rate. The *R*^2^ of this regression is very small (female “number of live births”: 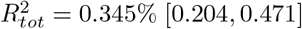, indicating that the correlates of linked selection explain a small but significant proportion of the variance in estimated effect sizes across SNPs (Tab. 1). Local recombination rate explains none of the variance, once B-value is included, likely because B-value is a function of recombination rate, and so we omit it from further analyses. Both LD score and B-value contribute to the variance, but we do not interpret this as indicative of the mode of linked selection. Rather, we consider that the B-value absorbs the contribution of linked selection that is not captured by the LD to common alleles, that is the contribution of rarer alleles in a SNP’s genomic environment (*C*_*x,R*_ in Eq. 6).

**Figure 3:**
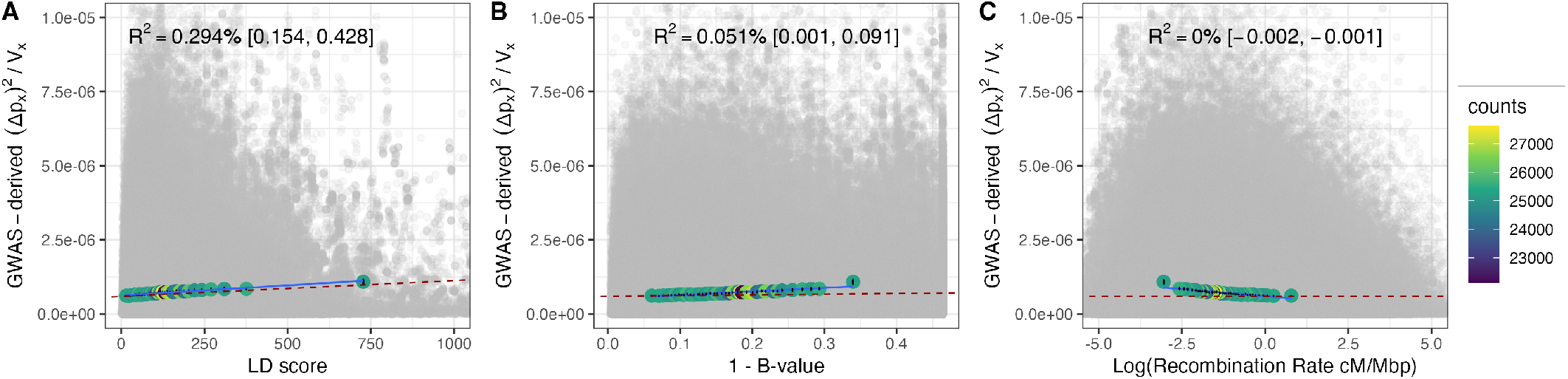
Decomposing the squared allele frequency change inferred from population GWAS effect sizes for common SNPs. Here the GWAS is in females on the phenotype ‘number of live births’. Each grey dot is a single SNP’s estimated squared allele frequency change, normalized by *V*_*x*_ (Eq. 8). In each panel the same set of SNPs are plotted by the **A)** LD score, **B)** 1 − B-value, **C)** log_*e*_ recombination rate. The colored points show the average calculated in 2.5% quantiles of the LD score predictor. The dashed red line shows the slope obtained from the multivariate regression through the underlying data, the *R*^2^ is reported as a percentage at the top of each panel. The blue line shows the linear regression on the binned values as a guide to the eye.

**Table 1:** Proportion of total variance and total change in squared allele frequency attributable to linked selection. *R*^2^ denotes the coefficient of determination for the multivariate regression with LD score, B-value, and log_*e*_ recombination rate. 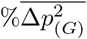 and 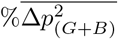 represent the percentage of squared allele frequency change explained by LD score alone or LD score and B-value, respectively. Unscaled and scaled intercepts from LD score regression (*C*_*x,R*+*d*_ and *χ*_0_) are also provided. Results are shown for female, male, and combined-sex WB cohorts, as well as for the MAF-partitioned analyses (Strat. columns), all based on GWAS analyses without PCs covariates. All values are reported as point estimates with 95% confidence intervals derived from a leave-one-chromosome-out jackknife.

|  | Female | Male | Strat. Female | Strat. Male | Combined Sex |
| --- | --- | --- | --- | --- | --- |
| $R^2_{\text{tot}}$ | 0.345% [0.204, 0.471] | 0.096% [0.013, 0.167] | 0.482% [0.21, 0.657] | 0.146% [0.0, 0.238] | 0.459% [0.231, 0.666] |
| $R^2_{\text{LD score}}$ | 0.294% [0.152, 0.426] | 0.077% [0.004,0.14] | 0.422% [0.153, 0.605] | 0.125% [−0.011, 0.208] | 0.364% [0.14, 0.569] |
| $R^2_{\text{B-value}}$ | 0.051% [0.005, 0.095] | 0.019% [−0.001, 0.037] | 0.060% [0.009, 0.102] | 0.021% [−0.001, 0.041] | 0.095% [0.045, 0.144] |
| $R^2_{\text{rbp}}$ | 0.0% [−0.002, 0.0] | 0.0% [−0.001, 0.001] | 0.0% [−0.001, 0.0] | 0.001% [−0.001, 0.002] | 0.0% [−0.002, 0.001] |
| $\overline{\% \Delta p^2_{(G)}}$ | 4.72% [2.79, 6.53] | 2.52% [0.92, 4.03] | 4.90% [3.62, 5.98] | 2.97% [1.31, 4.10] | 5.20% [2.70, 7.57] |
| $\overline{\% \Delta p^2_{(G+B)}}$ | 7.34% [5.54, 9.05] | 4.29% [2.32, 6.17] | 7.69% [6.02, 9.043] | 4.8% [2.65, 6.44] | 8.93% [6.30, 11.46] |
| $C_{x,R+d, \text{uni}}$ | $6.13 [6.01, 6.27] \times 10^{-7}$ | $8.60 [8.44, 8.76] \times 10^{-7}$ | $6.16 [6.07, 6.23] \times 10^{-7}$ | $8.61 [8.48, 8.77] \times 10^{-7}$ | $3.50 [3.50, 3.50] \times 10^{-7}$ |
| $C_{x,R+d, \text{multi}}$ | $5.94 [5.83, 6.07] \times 10^{-7}$ | $8.44 [8.25, 8.63] \times 10^{-7}$ | $5.94 [5.84, 6.03] \times 10^{-7}$ | $8.44 [8.26, 8.63] \times 10^{-7}$ | $3.55 [3.44, 3.67] \times 10^{-7}$ |
| $\chi^2_{0, \text{uni}}$ | 1.05 [1.02, 1.07] | 1.03 [1.01, 1.05] | 1.05 [1.04, 1.06] | 1.03 [1.01, 1.05] | 1.06 [1.03, 1.10] |
| $\chi^2_{0, \text{multi}}$ | 1.01 [0.99, 1.04] | 1.01 [0.99, 1.03] | 1.01 [1.00, 1.03] | 1.01 [0.99, 1.03] | 1.02 [0.99, 1.05] |

We estimated the relative, average contribution of selection 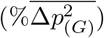 to Δ*p*^2^ by quantifying the contribution of local LD as 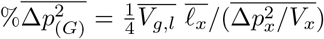 where 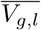 is the slope estimated from the partial regression of 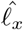 on 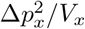. We find that the contribution of linked selection due to local LD is small 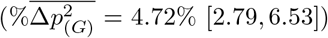 but the contribution of selection increases when we include the B-value 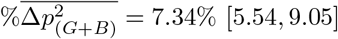, Tab. 1).

Selection over many generations will drive larger effect alleles to lower frequency, which is seen empirically for many human traits (e.g. O’Connor *et al*., 2019; Simons *et al*., 2025). Therefore, we ran the partitioned LD score regression stratifying the squared genotype correlation of each focal SNP by the minor allele frequency (MAF) of the other loci (Finucane *et al*., 2015), which slightly increased the estimated contribution of selection to allele frequency change (Tab. 1, Tab. S4, S5, Tab. S10, S11, Fig. S8-S11, Fig. S18-S21). Thus, linked selection due to differences in reproductive success makes a small but detectable contribution to allele frequency change in our sample.

In turn, genetic drift accounts for much of the average change in allele frequency, ∼ 93% of the squared change in the female WB cohort. The contributions of environmental variation in reproductive success 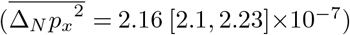 and Mendelian segregation to genetic drift are of similar magnitude 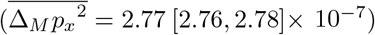. Our calculation of the environmental variation in reproductive success is closely related to the intercept of a typical LD score regression on *χ*^2^ GWAS summary statistics (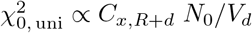, SI Appendix S4). This intercept will be 1 when deviations of the *χ*^2^ statistics from the LD score model are attributable just to individual-level sampling noise, but it will be greater than 1 if there are additional unaccounted for sources of variance in GWAS effect sizes uncorrelated to the LD score, including environmental confounding (Bulik-Sullivan *et al*., 2015) or departures from the LD score regression model (Loh *et al*., 2018). Our estimate of the intercept from the univariate LD score regression is significantly higher than the intercept value of a model where individual-level sampling noise is the only source of variance in effect sizes (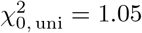, Tab. 1, Fig. S7-S17). However, we find the intercept in our multivariate regression to nearly match the unconfounded prediction (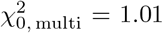 for a hypothetical SNP experiencing the weakest linked selection LD score = 0, B-value = 1, highest recombination bin value) suggesting that there is no strong contribution of environmental confounding to the squared allele frequency change. This is consistent with our finding that our analyses are insensitive to including the genetic PCs. Indeed, we find that our calculated level of allele frequency change due to drift in females 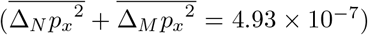 is close to the theoretical prediction for a panmictic population with a matched effective population size given by the sample size and variance in offspring number (4.18 *×* 10^−7^, see Eq. 17, Wright, 1938).

Our estimates of the proportion of allele frequency change due to linked selection through males are somewhat lower than in females (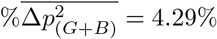; Fig. S12-S17, Tab. 1). This lower estimate in males reflects three factors: (i) the variance in reproductive success is higher in males than females 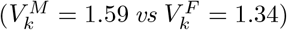, such that there is more allele frequency change through males, (ii) the slope of the LD-score regression is somewhat lower in males than in females, such that less of this allele frequency change is attributable to selection as opposed to drift, (iii) the cohort size of male is smaller than that of females, introducing more sampling noise in the effect sizes and hence more drift. To account for (iii), we adjusted the contribution of drift in females to match the lower cohort size in males (see Methods 1.6). In doing this, the relative contribution of selection to allele frequency change in females is closer to the estimate from the male cohort but still not as low 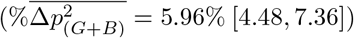. The female estimate decreases slightly further if we rescale the residual variance in reproductive success (numerator in Eq. 21) by the ratio in the variance in reproductive success between the two cohorts 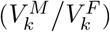, giving 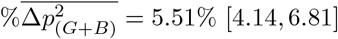. Overall, based on these calculations, this suggests that ∼ 60% of the difference in 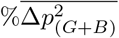 between males and females cohorts is due to increased drift and ∼ 40% to the lower effect of linked selection in males.

We can also look at the relative effects of selection and drift in the combined sample of females and males. Few of the male and female individuals in the UK Biobank are parents of the same families and so we approximate the combined across sexes contribution of variance in reproductive success using the sex-average effect 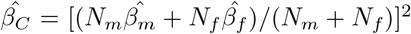. This represents the allele frequency change from the contributions of male and female transmissions in the WB cohort (see Methods for alternative approaches to combine male and female effect sizes; Fig. S22-S27; Tab. S13, S17). In doing this, we find that the relative contribution of linked selection increases somewhat in the combined sex sample 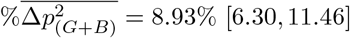, Tab. 1) reflecting the increased, pooled sample size that decreases the contribution of genetic drift. This is expected since the relative contribution of drift *versus* selection depends on the size of the cohort considered (Eq. S26), all else being equal. Below we show empirically how the contribution of drift *versus* selection changes with the cohort size and how it can be extrapolated to larger cohorts with the same demographic characteristics. But first we examine how much of the effect of selection on allele frequency change in the WB cohort is attributable to direct *versus* non-direct genetic effects.

### The genetic contribution to allele frequency change arises through direct effects

Our population-based GWAS effect sizes 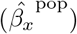 capture the direct effects (*β*^*D*^), i.e. the effects of nearby causal alleles that are in LD with the focal allele in the genotyped individuals, but also other sources of environmental confounding and non-direct genetic effects (Young *et al*., 2019; Veller and Coop, 2024, see SI Appendix S2). These non-direct genetic effects include indirect parental effects at a locus (Wolf *et al*., 1998; Kong *et al*., 2018) as well as genetic confounding due to systematic long-distance LD between loci induced by assortative mating and prior generations of selection (Bulmer, 1971, 1974). While estimates of heritability and genetic correlations based on the slope of the LD score regression should be free from most forms of environmental confounding, they still absorb non-direct genetic effects (SI Appendix S3.1; Border *et al*., 022a,b; Veller and Coop, 2024). In many applications the inclusion of these non-direct genetic effects is viewed as a source of bias. However, the contribution of selection to allele frequency change at a locus will include indirect genetic effects and correlated selection on other unlinked loci (Barton and Servedio, 2015), and therefore they should be included in the estimated contribution of selection.

To understand the role of direct genetic effects to selection-driven allele frequency change, we conducted a sib-GWAS on the WB cohorts (2, 383 male sibs and 4, 245 female sibs; see Methods). The sib-GWAS SNP effect sizes will be an estimate of the direct effect 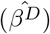 under the assumptions that the siblings are representative of the WB samples as a whole, that there are no indirect sib effects, and that genotypes are randomly distributed across interacting backgrounds (Veller *et al*., 2024). The estimated sib-effects have a lot statistical noise due to the small sibling-study sample size and, as a consequence, the LD score regression using the squared sib-GWAS effect sizes returns a nonsignificant slope (Fig. S28). The population-based GWAS effect size at a SNP can be decomposed into the contribution of the direct genetic effect, the non-direct genetic effect, and environmental noise and confounding 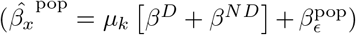, thus the squared population-based effect and hence the squared allele frequency change will reflect the sum of the pairwise product of these various terms (Eq. S64 in SI Appendix S3.1). It follows that the total contribution of the direct effects to the squared allele frequency change arises from the squared direct effect and the covariance between direct and non-direct genetic effects (*β*^*D*^(*β*^*D*^ +*β*^*ND*^); Eq. S64). We estimate this total contribution by performing LD score regression on the product of the estimated population-based and sibling-based GWAS effect sizes at each SNP (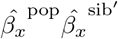 ; Fig. 4C; see SI Appendix S3.3). We find that the slope of this LD score regression is similar to the one obtained from the population-based GWAS LD score regression (Fig. 4A *versus* 4C). Thus, much of the signal of linked selection is due to genetic effects and not to environmental confounding alone, as environmental confounding would not be correlated with the direct effects across SNPs. The slope of this LD score regression, corresponding to the additive genic covariance of the direct and population effect 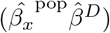, is closely related to the genetic variance used to define the allele frequency change in the fundamental and secondary theorems of natural selection, forming a link to classical results (Fisher, 1941; Crow and Nagylaki, 1976, see SI Appendix S6). Previous work has shown that much of the genetic variance for number of offspring, as captured by population-based GWAS effect sizes, arises in part from various forms of non-direct genetic effects (Howe *et al*., 2022; Tan *et al*., 2024). Thus, while our results suggest that the signal of linked selection is primarily due to genetic effects, rather than environmental confounding, much of this signal may be due to a tangle of diffuse correlations between selected loci distributed across the genome.

**Figure 4:**
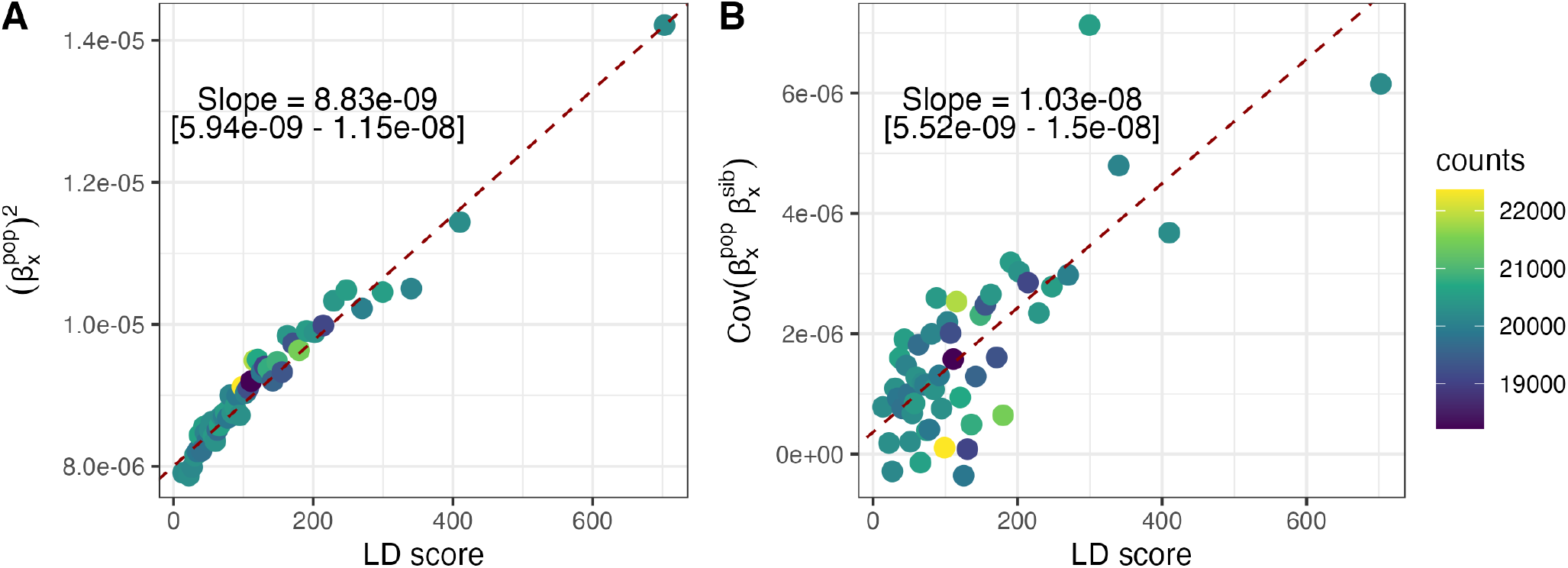
LD score regression using population and sibling-based GWAS for the female phenotype ‘number of live births’. Points show average values in bins of 2% quantiles of LD score. The linear regression through the underlying data is shown, and summary statistics are given as text. **A)** Squared population-based GWAS effect sizes 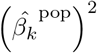 as in Fig. 3. **B)** The product of the effect sizes from population-based and sibling-based GWAS 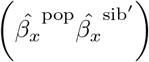. The SNP effect sizes from sib- and population-GWAS have slightly different adjustments for inbreeding (*F*_*x*_) to place estimated effects on the same scale (see Eqs. S68, S77 and S84).

### The scaling of allele frequency change with cohort size

A key determinant of the rate of genetic drift is population size and so we investigated how the contribution of selection scales with cohort size. Up to this point, our estimates have focused on the contributions in the WB cohort; to examine the effect of smaller cohort size, we randomly downsampled the WB cohort to a range of sample sizes (we do this with the female WB cohort to avoid the statistical issues of combining the sexes). Downsampling should not affect the expected additive genic variance. At small sample sizes, however, the estimated slope of the LD score regression is noisier, and we observed a weak downward bias in the estimated additive genic variances (Fig. 5A). However, the empirical estimate of the slope of the LD score regression is correctly recovered, on average, for the larger sub-samples (e.g. using 60% of the WB cohort). As we move to smaller cohort sizes, the genome-wide average change in frequency is larger, as allele frequencies are more variable due to drift (blue points in Fig. 5C). Thus, we find that selection contributes less to allele frequency change in smaller cohorts (blue points in Fig. 5B).

**Figure 5:**
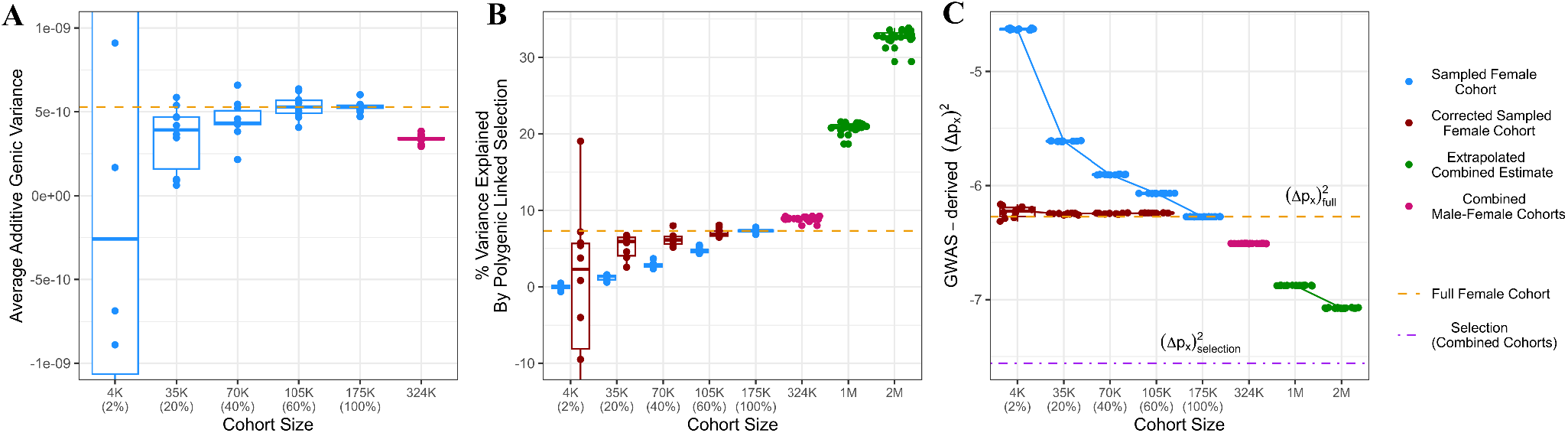
The effect of female WB cohort size on: **A)** The estimated average additive genic variance per site (slope of the LD score regression). **B)** The percentage of change in squared allele frequency change explained by selection. **C)** log_10_ of the estimated average total squared allele frequency change per site. Each blue dot shows the measure of interest computed on random subset of individuals for a given sample size. The red dots show the subsample measures corrected back to the full WB cohort size. The boxes show the quantiles and average across 10 replicated subsamplings. The orange line shows the measure calculated in our full WB cohort. The green points show the extrapolation of the squared allele frequency change to hypothetical larger WB cohorts.

We can also predict the relative contribution of selection in larger cohort sizes. We begin by demonstrating how we can scale back up to the full WB cohort from our downsampled cohorts (Methods 1.6). In each subsample, we adjusted the average total squared change in allele frequency (blue points in Fig. 5C) to reflect the greater averaging of individual-level noise in a larger sample size. In doing this adjustment, we are able to use the smaller sample sizes to recover the total squared allele frequency change in our full WB cohort (red points in Fig. 5C) and, in larger cohorts (40% − 60% of the full WB cohort), to accurately recover the fraction of allele frequency change due to selection in the full WB cohort (red points in Fig. 5B). Based on this adjustment, we extrapolated how much lower the total squared allele frequency change would be at larger sample sizes (green points in Fig. 5C), assuming a sample from a hypothetical population whose genetic constitution and demographic properties match those of the WB cohort. As the contribution of selection would remain on average the same in this hypothetical larger cohort (purple line in Fig. 5C), selection is expected to play a larger role in the squared allele frequency change. Importantly, this larger relative role does not reflect a larger absolute contribution of selection, but merely the diminishing role of drift in allele frequency change in larger cohorts. For example, the average change in allele frequency 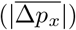 is projected to drop from 7*/*10, 000 in the female WB cohort (N = 175, 228) to 3.65*/*10, 000 in a WB-like cohort of a million individuals.

In extrapolating to larger cohorts, we are making the strong assumption that the WB cohort is representative of larger samples. However, this assumption is unlikely to hold given that the WB cohort is a particular subsample of the UK population and, in addition, was recruited by opt-in strategy, and therefore is subject to various sample-selection biases (Fry *et al*., 2017). If we were to sample large groups of people at random from across the UK, we would sample across various socio-environmental gradients that would be correlated with genotypes. If these environmental gradients contribute to variation in offspring number, the relative contribution of genetic drift could actually increase in larger cohorts. In addition, the additive genic variance may change as a broader distribution of environmental heterogeneity as sampling expands. Finally, the size and composition of the cohort we are interested in depends on the question we wish to address. Do we wish to understand the contribution of drift and selection in the past generation in the UK? Or in some much broader cohort?

### Concluding remarks

Over longer time periods, the contribution of selection on allele frequency change can be magnified, as changes due to selection are compounded over generations, while changes due to drift usually are not (Santiago and Caballero, 1998; Buffalo and Coop, 2019, 2020). However, despite a number of convincing examples of selection at a small number of loci from ancient DNA (e.g. Mathieson and Terhorst, 2022; Irving-Pease *et al*., 2024; Akbari *et al*., 2026), recent work has found that selection seems to play a small role in genome-wide allele frequency change over the past thousands of years in Europe (Simon and Coop, 2024; Akbari *et al*., 2026). While recombination breaks down the associations of an allele to selected loci over time, reducing the compounding effect of linked selection, our LD scores are calculated between SNPs at a relatively small genomic scale (a centi-Morgan), and so the associations that they capture could be compounded over a hundred of generations. Yet we find no evidence of a genetic correlation between our estimates of allele frequency change and the temporal gradients in allele frequency estimated from ancient DNA by Akbari *et al*. (2026) (Fig. S29). This is consistent with the view that environments and selective pressures have shifted many times over thousands of years, preventing selection from compounding as efficiently. The compounding effect of selection on allele frequency change will also be limited if the alleles that contributing to selective change turnover on short time-scales (e.g. as they would under strong background selection).

Our results focused on the consequences of variance in reproductive success due to drift and selection following a single cohort of individuals, and therefore ignore the consequences of migration to allele frequency change. Previous work on ancient DNA has found that gene flow may have been the dominant factor in allele frequency change in the UK (Simon and Coop, 2024), and elsewhere in Europe, over the past thousands of years in the history of people we today associate with European genetic ancestry. This change reflects demographic expansions and historical migrations of various groups to and within Europe (Haak *et al*., 2015; Allentoft *et al*., 2015). Therefore, the small changes we infer from selection and drift may be dominated by changes that can occur through migration over short time-scales in time series data (Chen *et al*., 2019; Simon and Coop, 2024). One open challenge is how to use estimates of the contribution of migration, genetic drift, and selection from short time slices and time series to understand these processes role in shaping current day patterns of genetic diversity, as diversity in the present dat reflects how these processes played out in the genetic ancestors of a sample over much longer time-scales.

Our analyses could be extended to explore the contributions of drift and selection across components of lifetime reproductive success in age structured populations such as humans (indeed the large cohort designs of human genetics may lend themselves to such studies Lewontin, 1968; Byars *et al*., 2010; Sanjak *et al*., 2018; Mostafavi *et al*., 2017). Applying these approaches to other species could also allow a detailed exploration of these factors across species. That said, our LD score regression approach requires very large sample sizes to be accurate, which places it out of reach in most non-human systems. Our use of LD score regression follows from the derivation of allele frequency change based on the sum of LDs (Eqs. S12-S18). However, at its core, our approach relies on estimating the additive genic variance for fitness, and a closely related set of approaches estimates this quantity based on the genetic relatedness matrix (Bulik-Sullivan, 2015; De Vlaming *et al*., 2017; Geeta Arun *et al*., 2026). These approaches can be more statistically efficient than LD score regression (Visscher *et al*., 2014; De Vlaming *et al*., 2017; Geeta Arun *et al*., 2026) and so may offer a way of estimating the contribution of drift and selection to allele frequency change using much smaller samples. The heritability of fitness is low but measurable in many natural populations (Kulbaba *et al*., 2019; Shaw, 2019; Bonnet *et al*., 2022), suggesting that decomposing allele frequency change to understand the short-term genome-wide response to selection and drift should be possible in a wide range of species.

## Acknowledgments

We thank Jeremy Berg, Doc Edge, Chuck Langley, Molly Przeworski, Guy Sella, Carl Veller, and members of the Coop lab for helpful discussions and comments on earlier drafts. This research has been conducted using the UK Biobank Resource under Application Number 75040. Funding was provided by the National Institutes of Health (R35 GM136290, awarded to GC). The authors acknowledge the High Performance Computing Core Facility at the University of California, Davis, for providing computational resources.

## 1 Material and Methods

### 1.1 Simulation Set-Up

We carried out simulations of a Wright–Fisher population in SLiM v4.0.1 (Haller and Messer, 2023). We considered four scenarios: *Neutral, Directional linked-selection, Stabilizing linked-selection*, and *Directional linked-selection with Assortative Mating*. Across all scenarios, we simulated a constant-size population of 10, 000 diploid individuals, each carrying a single 5 Mbp chromosome with a mutation rate of 1 *×* 10^−8^ per bp per generation. We tested several uniform recombination rates (1 *×* 10^−9^, 1 *×* 10^−8^, 1 *×* 10^−7^, and 9 *×* 10^−7^ per bp per generation) as well as the recombination map of human chromosome 22, rescaled to the 5 Mbp length of the simulated chromosomes. We note that our goal here is not to simulate under realistic parameters for humans, but merely to have recombination variation in our simulations.

Polygenic linked-selection was simulated assuming an additive polygenic trait, such that the breeding value of the *i* individual is 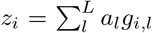, where *g*_*i,l*_ ∈ {0, 1, 2} is the allele count at selected locus *l* and *a*_*l*_ is the additive effect size. To ensure sufficient initial genetic diversity for selection to act on, for each scenario we first simulated a constant-size population of 10, 000 diploid individuals under neutrality in msprime (Baumdicker *et al*., 2022), recording the output as a tree sequence. The tree sequence was then edited to assign a non-zero effect size to 25% of variant sites, with effect sizes drawn at random from a normal distribution with mean 0 and standard deviation set to one of 0.413 *×* 10^−2^, 0.826 *×* 10^−2^, 1.65210^−2^, or 2.478 *×* 10^−2^. The resulting tree sequence was then loaded into SLiM and the simulation was run for one generation, with new mutations being uniformly added with equal probability as either neutral or with effect size drawn from the aforementioned distributions. *Directional linked-selection* was simulated using an exponential fitness function on the additive polygenic trait. *Stabilizing linked-selection* was simulated using a Gaussian fitness function centered on an optimum of 0 with a variance *V*_*s*_ = 1.

#### Directional linked-selection with Assortative Mating

We wished to simulate a period of assortative mating for ten generations prior to selection to allow the build up of systematic long distance LD. Thus, we slightly modified the setup described above. We used the same edited tree sequence as for the other linked-selection simulations, in which neutral and selected variant sites are distinguished by labeling their mutations “m2” and “m1”, respectively, in addition to the assigned effect sizes. For the assortative-mating (AM) simulations, we re-edited this tree sequence to set all effect sizes to zero while retaining the “m1”/”m2” classification. The re-edited tree sequence was then loaded into SLiM and the simulation run for 10 generations under AM without selection. Assortative mating was modeled by first taking each individual’s count of “m1” alleles as their phenotype, then using the mateChoice() callback so that individuals choose mates by phenotypic similarity, according to a Gaussian mating-preference function with mean equal to the focal individual’s phenotype and standard deviation 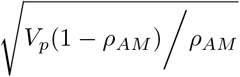 where *V*_*p*_ is the population phenotypic variance and *ρ*_*AM*_ is the target AM correlation. This expression for the standard deviation is derived from the joint probability density of a mated pair and its precision matrix. After the 10 generations of AM, the tree sequence at the final time point was recorded and subsequently edited to add effect sizes to the “m1” sites and use them as the basis of the selected phenotype. This final tree sequence was then loaded into a SLiM simulation combining directional selection, as described above, with the mateChoice() callback for AM and was run for one generation. Results were recorded as tree sequences and analysed in Python using tskit (Wong *et al*., 2024; Kelleher *et al*., 2016). All results are based on 100 replicates of each scenario. Code is available at https://github.com/gsgarlata.

### 1.2 UK Biobank Data Analysis

The UK Biobank is a large-scale prospective cohort study including ∼ 500, 000 individuals aged between 37 − 73 years, who were recruited between 2006 and 2010 from across the United Kingdom. We used the UK Biobank imputed genotype data version 3 from 487, 713 individuals (field ID 22418). Genotyping, quality control and genotype imputation were conducted by UK Biobank, Wellcome Trust Center for Human Genetics (WTCHG), University of Oxford, UK (Bycroft *et al*., 2018).

We followed standard sample quality controls (Bycroft *et al*., 2018) by removing individuals with sex-chromosome aneuploidy (field ID 22019), discordant genetic sex (field ID 22001) and self-reported sex (field ID 31), heterozygosity and missing data outliers (field ID 22027) and individuals with *>* 10% of missing data. To reduce the effects of population structure on our analyses, we used a set of individuals with relatively homogeneous ancestry. In particular, we retained individuals who self-identified as ‘White British’ (field ID 21000), fell within 7 standard deviations of the centroid along the first 6 genetic PCs from a principal components analysis of the genotypes (field ID 22006), and are unrelated (field ID 22020). We retained only high-quality SNPs with missingness *<* 0.1, Hardy-Weinberg equilibrium (HWE) test P-value *>* 1*e* − 5 *×* 10^−*N·*0.001^, where *N* is the sample size, (Greer *et al*., 2024) with mid-p adjustment (“midp”) (Graffelman and Moreno, 2013) and filtering out variants with excess heterozygosity (“keep-fewhet”), non-multiallelic, imputation quality (INFO) *>* 0.8, minor allele frequency (MAF) *>* 0.001 and minor allele count (MAC) *>* 100. Filtering was performed in plink v2.0 (Chang *et al*., 2015).

This final “White British” (WB) UK Biobank sample was then separated in a female and male cohort (field ID 31), for which we retrieved phenotypic data related to reproductive output, “number of livebirths” for females (field ID 2734) and “number of children fathered” for males (field ID 2405). The genotype dataset was then filtered to individuals with this phenotype. Population GWAS analyses were performed on the WB sample of unrelated individuals, resulting in a female and male WB cohort of 175, 228 and 151, 675 unrelated individuals with ∼ 1 million SNPs.

We also performed sibling GWAS analyses, for each sex we obtained a full-siblings WB cohort using UK Biobank provided estimates of kinship coefficient (*ϕ*) and the proportion of SNPs for which the individuals share no allele (IBS0) (available at the file “Bulk/Genotype Results/Genotype calls/ukb rel.dat” in the UK Biobank database). To call sibling pairs we used the following conditions: 0.1767767 *< ϕ <* 0.3535534 and IBS0 *>* 0.0012 (Bycroft *et al*., 2018). We then filtered these sibling pairs such that both individuals were in the WB grouping, their reported sex matched their inferred sex, were not identified by the UK Biobank as heterozygosity and missing data “outliers”, and had reproductive output information. This resulted into a female and male WB cohort of 4, 245 and 2, 383 full-siblings pairs with ∼ 1 million SNPs.

We removed all of the sibling-based GWAS individuals from the set of individuals used for the population-based GWAS, to ensure the two sets of GWAS were independent.

### 1.3 GWAS

Population GWAS analyses were carried out with REGENIE v3.3 (Mbatchou *et al*., 2021), a machine-learning method that uses a two-steps process to estimate effect sizes across the genome. We also performed GWAS analyses including the first 20 principal components (field ID 22009) as covariates. The reproductive output trait used in these analyses was not standardized. Family GWAS was performed using genetic differences between full siblings with snipar (Young *et al*., 2022). Note that we further validated our analyses using a different set of population GWAS effect sizes on the same phenotypes (Howrigan *et al*., 2023), which gives qualitatively similar results (see Fig. S30, S31).

### 1.4 LD Score Regression

Multivariate weighted LD Score regression was used to estimate the average contribution of selection, environment and Mendelian segregation on squared allele frequency change divided by observed genotypic variance 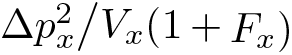. The squared allele frequency change across SNPs was estimated from the GWAS effect-sizes for the number of children, and thus includes only the selection and environmental contributions (i.e., no information on Mendelian segregation) (Section S3).

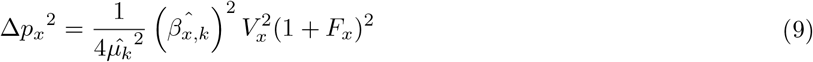

where 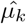 is the estimated mean fitness, obtained by calculating the average number of children in the focal cohort, 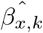 is the GWAS effect-size, *V*_*x*_ is the expected genotypic variance at site *x* and *F*_*x*_ is its inbreeding coefficient. We then perform the multivariate regression:

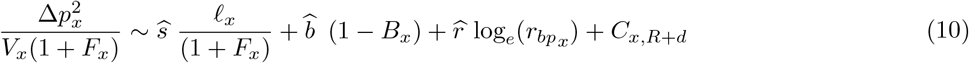

using LD score (*ℓ*_*x*_), 1 - B-value (1 − *B*_*x*_) and log_*e*_ of recombination rate 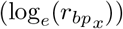 as our predictors of linked selection. *C*_*x,R*+*d*_ is the intercept of the LD score regression, absorbing the contributions of environment, and polygenic linked-selection not well captured by our linked selection predictors. 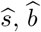 and 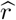 are the regression coefficients of the predictors. Note that we use 1 − *B*_*x*_, instead of *B*_*x*_, as one of the predictors, so that it shares the same positive relationship with squared allele frequency change as LD score and recombination rate. This will be useful when we estimate the squared allele frequency change of the hypothetical SNP experiencing the weakest linked selection (LD score = 0, B-value = 1, highest recombination bin value).

For the analysis of the UK Biobank cohorts, we used LD Score values from Gazal *et al*. (2017), computed from 9, 997, 231 biallelic SNPs with minor allele count ≥ 5, which were calculated in a set of 489 unrelated European individuals (EUR) from the 1000 Genomes Project phase 3. LD scores values for the MAF partitioned and unpartitioned analyses are available at https://zenodo.org/records/10515792. For the MAF partitioned analysis we used the LD scores from this file 1000G_Phase3_baselineLD_v2.3_ldscores.tgz. We retrieved the B-values from Murphy *et al*. (2022), which were estimated from the Yoruba population using the 1000 Genomes Project phase 3 VCF files spanning all 26 populations from across the world. The recombination map was retrieved from Bhérer *et al*. (2017) (Refined_EUR_genetic_map_b37). In particular, we used the refined European maps, derived from the 97, 723 meioses from individuals of European origins. In all these analyses, we removed SNPs with MAF *<* 0.05.

Following (Bulik-Sullivan *et al*., 2015), to correct for the correlation between SNPs, the multivariate LD score regression is weighted by the inverse of the LD score computed using only the *S* SNPs included in the regression:

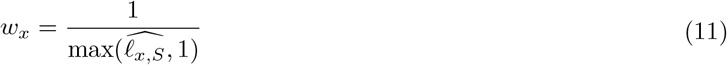

#### Partitioned LD score regression

We also performed partitioned LD score regression (Finucane *et al*., 5 11) on categories defined by MAF bins, an approach that allows each MAF category to contribute differently to the total additive genic variance. Following this approach, we estimated the contribution of each MAF category to polygenic linked-selection. We included ten MAF bins: [0.050, 0.071], (0.071, 0.098], (0.098, 0.131], (0.131, 0.171], (0.171, 0.215], (0.215, 0.265], (0.265, 0.319], (0.319, 0.377], (0.377, 0.439], (0.439, 0.5]. For each variant in the dataset, ten LD score measures were obtained, one for each MAF category, computed between the focal variant and all other variants in that category. The multivariate LD score regression then takes the form:

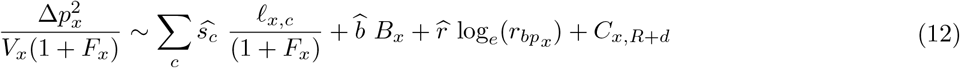

where *c* indicates the MAF category and *ℓ*_*x,c*_ is the LD score of the focal variant with the variants within the MAF category.

#### Estimating the average allele frequency change due to selection

The estimated slopes from the regression are then used to estimate the squared allele frequency change due to selection (Section S2):

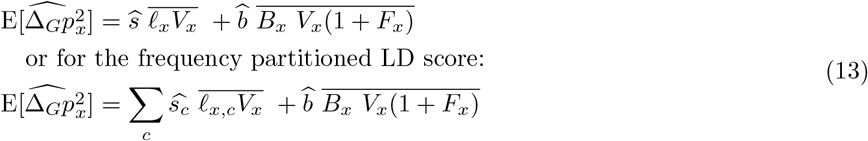

The squared allele frequency change due to the environmental variation in reproductive success is estimated as

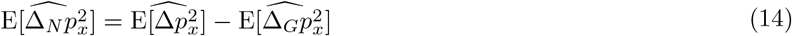

and the average squared allele frequency change due to the Mendelian segregation (Section S1.3) as:

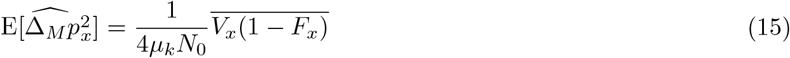

To estimate the proportion of squared allele frequency change due to polygenic linked selection, we then used:

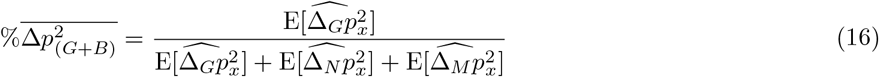

We also compared our estimates to the squared allele frequency change predicted by the theory for a panmictic population of the same size and with the same variance in offspring number Wright (1938):

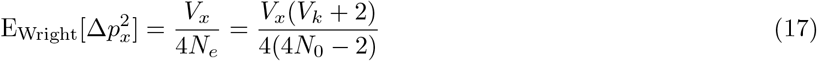

Confidence intervals were obtained by block jackknife resampling, using chromosomes as blocks. In particular, for each of the twenty-two autosomes, we removed all variants on that chromosome and re-estimated polygenic linked selection from the remaining data (Eq. 10,13-16). The resulting twenty-two leave-one-chromosome-out estimates were then used to construct a 95% confidence interval.

### 1.5 Combining GWAS Sex-specific Effect Sizes

The GWAS effect-sizes for the number of offspring have been obtained from two independent samples of males and females of size *N*_*m*_ and *N*_*f*_, respectively. We tested three approaches, two of which are typically used in GWAS meta-analyses, to aggregate effect-sizes from the two cohorts through weighted average: sample-size weighted mean, fixed-effect and random-effect. The “fixed-effect” and “sample-size weighted mean” models assume that both cohorts share the same true underlying population effect size (i.e., homogeneity) and that differences between cohorts are solely due to sampling error. The “random-effect” model considers that the true underlying population effect size may vary between cohorts due to, for instance, ancestry and environment or sex (i.e., heterogeneity). The “sample-size weighted mean” approach consists in weighting both effect-size and standard errors by the sample size of each cohort. By doing so, we assume that standard errors between cohorts differ exclusively in sample size but not in allele frequencies. The combined effect-size 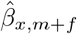, where *m* is for males and *f* for females, is given by:

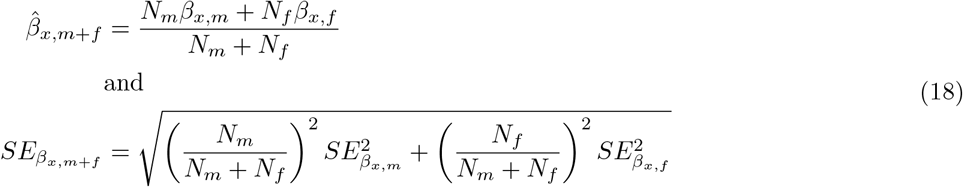

where 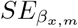 is the standard-error of the effect-size of cohort *m* (*β*_*x,m*_). Note that the expression for 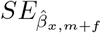 follows from the standard formula for the variance of a linear combination of independent estimates. The “fixed-effect” and “random-effect” models differ in the way the weights (*w*_*m*_ and *w*_*f*_) are computed (Borenstein *et al*., 8 24), but their general expression is given by:

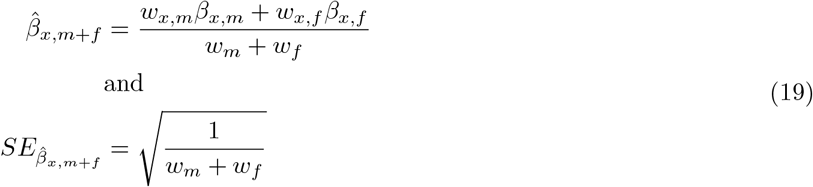

### 1.6 Extrapolating to Larger Cohort Size

In Eq. 9, we show that the genome-wide average squared allele frequency change can be estimated from the squared GWAS effect size for number of children average across loci. Here, we ask how this quantity scales with sample size, allowing us to infer what fraction of the squared allele frequency change reflects polygenic linked selection *versus* environmental variation in a hypothetical larger cohort. Consider a cohort of *N*_sam_ individuals, from which GWAS effect-sizes 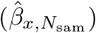 are estimated. Assuming that the true effect size at locus *x* (*β*_*x*_) is independently drawn from a normal distribution 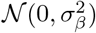, the OLS estimator 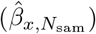 is itself normally distributed conditional on the true effect size, that is:

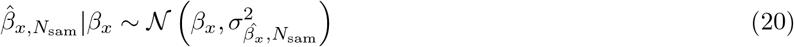

where 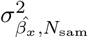 is the squared standard error of 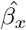 and is defined as

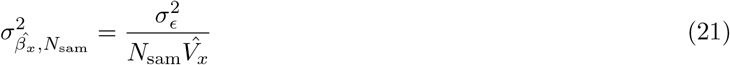

where 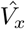 is the genotypic variance from *N*_sam_ and 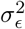 is the residual variance of the phenotype. Therefore, 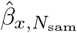 is normally distributed as 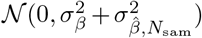. If we consider the expectation over loci, we can state that 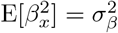 and 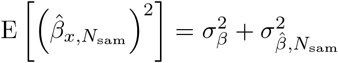. The aim of this section is to show that we can re-scale 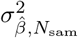 to a cohort of *N*_full_ individuals and obtain an estimate of 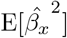 in such cohort. We can do that by replacing 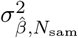 with 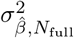, such that:

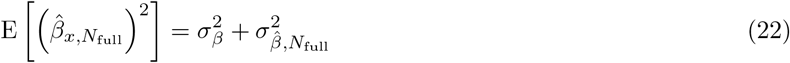

The true population variance in effect size 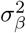 can be estimated from 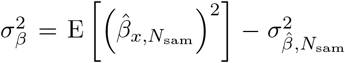, where the first term is obtained by averaging the squared of the GWAS effect-size for the number of children across *K* loci 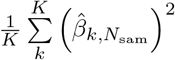 and the second by averaging the squared standard error of the estimated effect size 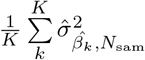. To account for the difference in the number of individuals between *N*_sample_ and *N*_full_, we can correct the expected squared standard error of the estimated effect size across all sites as:

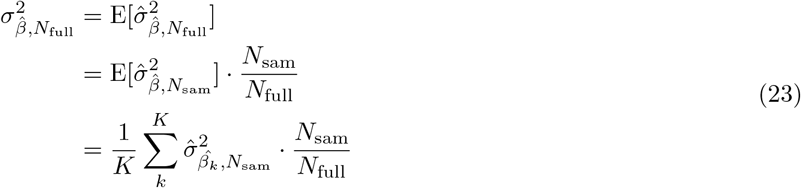

where 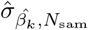 is the standard error of the effect size estimated from *N*_sam_ at site *k*. Using those estimates, we can infer the 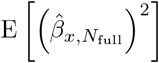 and therefore the genome-wide average squared allele frequency change in the *N*_full_ cohort.

### 1.7 Ancient DNA data analysis

We tested whether the change in allele frequency due to polygenic linked selection estimated from the UK Biobank cohort is correlated to polygenic linked selection signatures in earlier generations. We used the ancient allele frequency change effect sizes of Akbari *et al*. (2026), who combined DNA from 8, 433 unrelated ancient individuals spanning the past 14, 000 years with 6, 510 present-day individuals to infer temporal allele-frequency gradients, and hence genome-wide selection coefficients. In Section S5, we summarize why estimated selection coefficients are equivalent to the effect sizes obtained from a population GWAS on the number of children. We therefore treated the selection coefficients estimated by Akbari *et al*. (2026) as GWAS effect sizes and applied univariate LD score regression on the product between WB population GWAS effect sizes and estimated selection coefficients to test whether estimates of allele-frequency change in the UK Biobank cohort are genetically correlated with those obtained from ancient DNA temporal series.

## 2 Mathematical Appendix

## S0.1 Roadmap to Theory

We begin by writing a decomposition of the allele frequency change in a single generation (Section S1) and detail the genetic, environmental, and Mendelian segregation components (Section S1.1, S1.2, and S1.3 respectively). We show how the genetic component can be approximated in terms of the LD score (Section S2) and how this is affected by population structure (Section S2.2) and long distance LD (due to assortative mating and the Bulmer effect, Section S2.3).

We then show how we can write the predicted allele frequency change using GWAS effect-sizes for the number of children (Section S3). We discuss how population GWAS absorbs various non-direct effects and how these can be separated through (Family GWAS S3.3 and the product of the Population and Family GWAS effects S3.2).

## S1 Decomposition of Allele Frequency Change: Selection, Environment, and Mendelian Segregation

Consider a sampled cohort of *N*_0_ diploid individuals, where at a biallelic SNP *x*_*i*_ ∈ {0, 1, 2} indicates the diploid genotype of individual *i* at a given biallelic site. Then, the allele frequency in our initial cohort (0) can be expressed as:

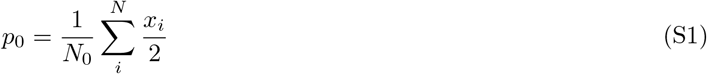

The *i*^*th*^ individual contributes *k*_*i*_ offspring to the next generation, which consists of *N*_1_ individuals 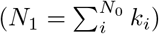, such that the corresponding allele frequency at time *t* = 1 is:

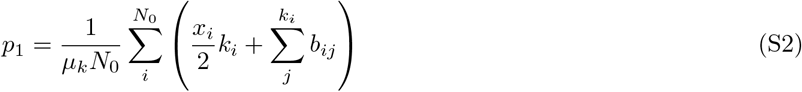

where (*x*_*i*_*/*2)*k*_*i*_ is the expected contribution of each individual to the next generation, given their number of children, and *µ*_*k*_ = 2*N*_1_*/N*_0_ is a rescaling factor of the cohort size in the following generation which is equivalent to the mean absolute fitness of the cohort. Mendelian segregation is captured by *b*_*ij*_, the deviation away from the expected genetic contribution of the *i*^*th*^ parent to child *j* due to the randomness of transmission out of heterozygotes. If the *i*^*th*^ individual is homozygous *b*_*ij*_ is zero, while for heterozygotes *b*_*ij*_ = 1*/*2 if the individual *i* transmits the focal allele to child *j*, and *b*_*ij*_ = −1*/*2 if it transmits the other allele. Assuming that Mendelian segregation is truly random at our locus, we can model this as 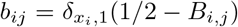 with *B*_*i,j*_ ∼ *Bernoulli*(1*/*2) and 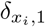 being an indicator function which is 1 if the *i*^*th*^ individual is heterozygous and 0 otherwise.

The allele frequency change over one generation can thus be expressed as:

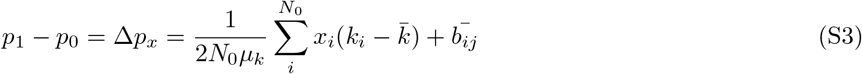

where 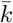 is the average number of offspring per individual and 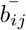 is the average random Mendelian contribution. Note that the first term in Eq. S3 can be interpreted as the allele frequency change due to variance in offspring number over individuals, which can be rewritten as the covariance between the focal locus *x* and the number of offspring *k*:

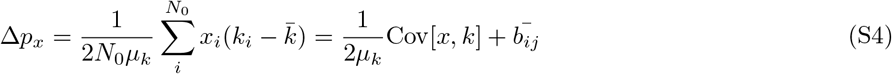

Note that this expression recalls the Robertson-Price equation, which describes how the population mean of a trait changes from one generation to the next.

Following a quantitative genetics approach (e.g. Santiago and Caballero, 1995, 1998; Buffalo and Coop, 2019) the number of offspring *k*_*i*_ can be divided into two terms: i) the expected contribution of individual *i* due to its genotype *f*_*i*_ (relative fitness); and ii) a deviation from *f*_*i*_ due to random sampling or environmental factors *d*_*i*_:

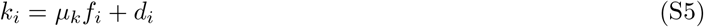

where E[*f*] = 1 and E[*d*] = 0. The allele frequency change can now be expressed as

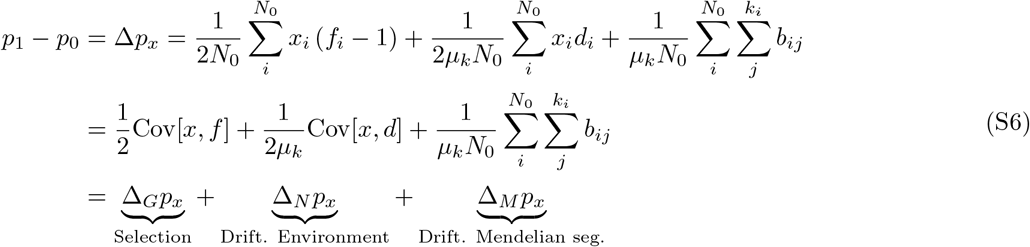

Since we have chosen to track a random allele at a given focal locus, we are not interested in the sign of the allele frequency change, but only in its magnitude. Therefore, we can focus on the squared change in allele frequency 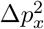, which under our decomposition and in expectation is

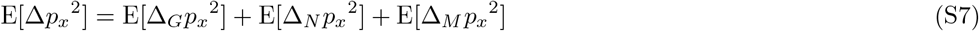

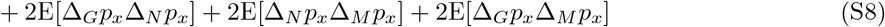

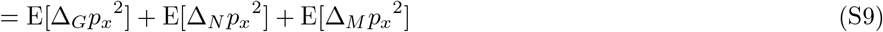

The cross terms involving Δ_*M*_*p*_*x*_ are zero in expectation as Mendelian segregation is random. We set E[Δ_*G*_*p*_*x*_Δ_*N*_*p*_*x*_] = 0, as we assume that genotype-environment correlations (between *f*_*i*_ and *d*_*i*_) are absorbed into *f*_*i*_ (we return to this point below when we consider sources of environmental confounding). Note that the expectation of Eq. S7 can also be interpreted as the variance in allele frequency change, since E[Δ*p*_*x*_] = 0, conditional on the tracking of a random allele.

### S1.1 Contribution of Genetic Selection Under Additive Model

We assume that the relative number of offspring of a single diploid individual given its genotype (individual relative fitness, *f*_*i*_) can be modeled as an additive trait. At each of the biallelic selected sites, 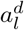 is the effect size of a randomly chosen focal allele, where *d* stands for direct genetic effect, and *g*_*l*_ is the diploid genotype at the selected site *l* (we return to indirect genetic effects below). Then the relative fitness can be written as

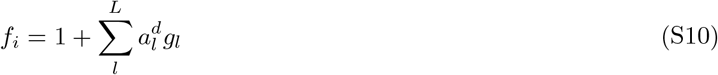

We can then write the genetic contribution to the allele frequency change at our focal site as 1

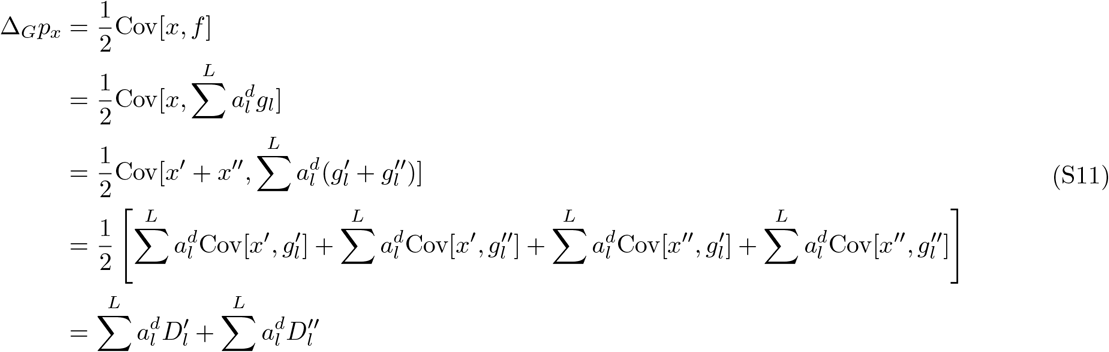

where *x* = *x*′ + *x*″ and 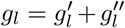 indicates that the genotypic value at a given site is composed of the contribution of each of the two gametes, such that *x*′ and 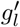 have been inherited from one parent and *x*″ and 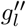 from the other. Since linkage disequilibrium *D* is defined as the genotypic covariance between two loci, and assuming no difference in LD inherited from mothers and fathers, 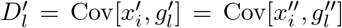 represents the linkage disequilibrium between alleles lying in the same gamete (called gametic or cis-LD), while 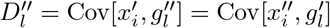 represents the linkage disequilibrium between alleles in different gametes (called non-gametic or trans-LD). Then, the genetic contribution to the squared allele frequency change is:

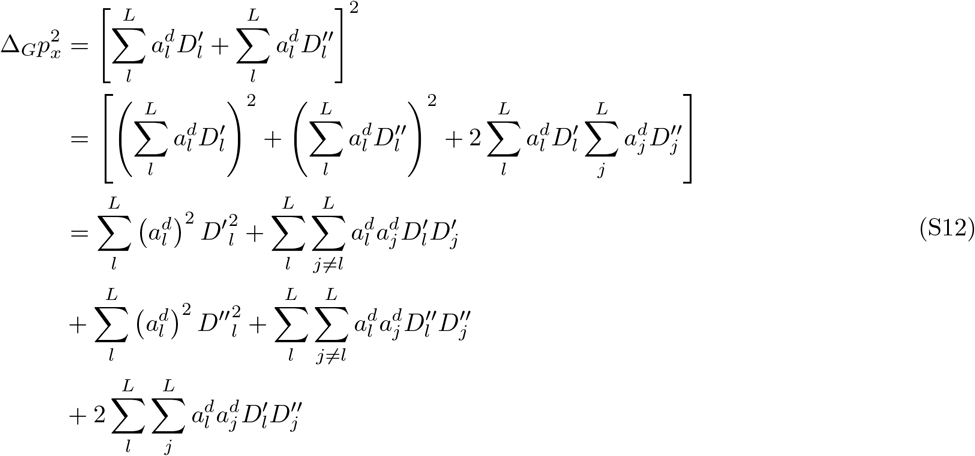

The first term can be expanded as:

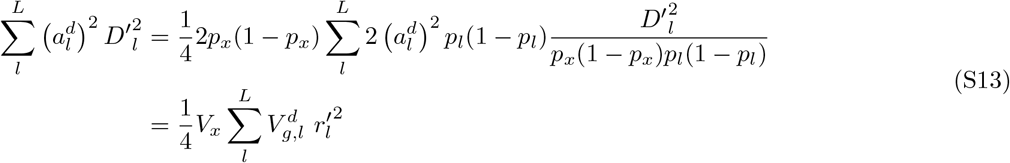

where *p*_*x*_ and *p*_*l*_ are the allele frequency at the focal site *x* and the selected site *l*, respectively, *V*_*x*_ = 2*p*_*x*_(1 − *p*_*x*_) is the expected genotypic variance in a panmictic population, 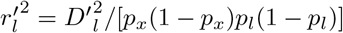 is the gametic squared Pearson correlation between the focal site *x* and the selected site *l* and 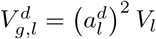 is the additive genic variance of site *l* for fitness. The second term leads to

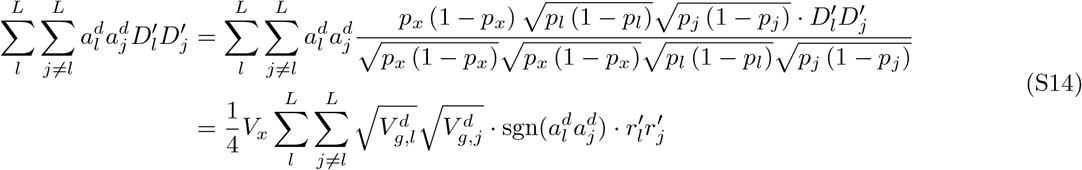

Similarly, the third term becomes:

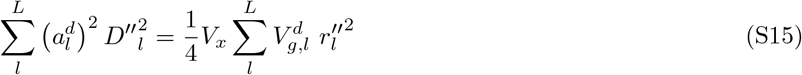

and the fourth term:

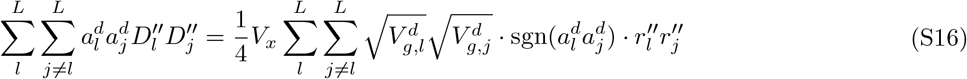

Finally, the fifth term can be written as:

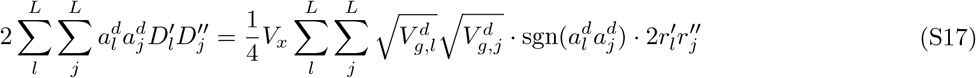

Therefore, 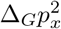 can be expressed as

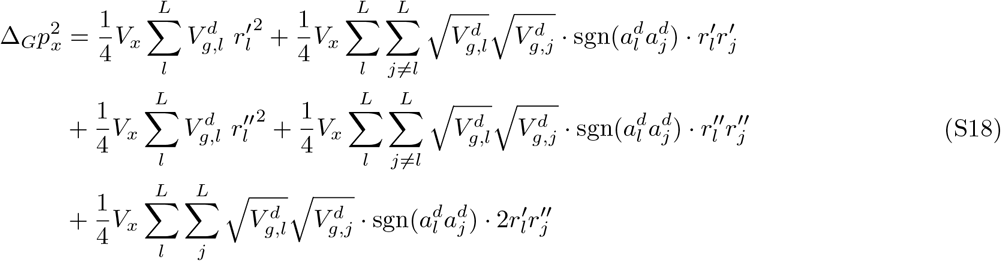

The first term on the right-hand side of Eq. S18 is the squared cis-LD between the focal sites and selected sites, and it is this term, after some approximations, that will give us the LD score regression. All subsequent terms on the right-hand side of Eq. S18 that carry the product of the effect size signs 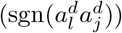 are zero in expectation, if there is no systematic signed LD due to the Bulmer effect or assortative mating. We return to the interpretation of these terms in Sections S2 and S2.3.

### S1.2 Contribution of Environmental Sources of Variance in Offspring Number

The general expression for the squared allele frequency change due to non-genetic contributions to offspring number in a population is given by:

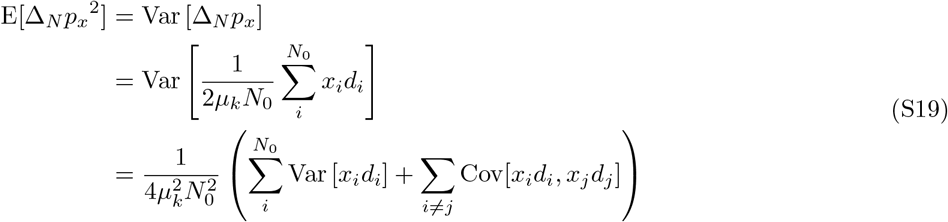

If the genotype at the focal locus *x* is statistically independent of *d* and there is also independence between *d*_*i*_ and *d*_*j*_, then Eq.S19 simplifies to:

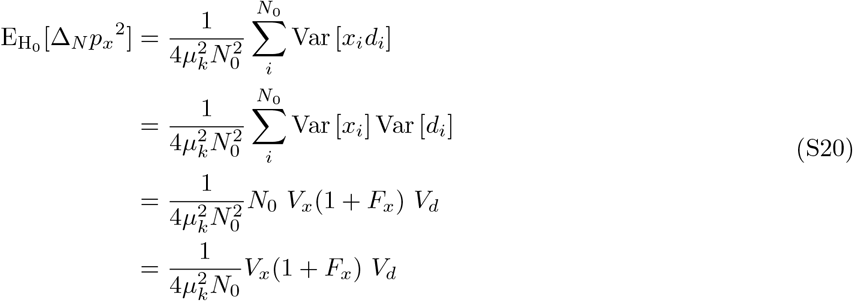

The independence of the environment contributions to allele frequency change is the common assumption in Wright-Fisher models, underlying the idea that drift scales inversely with the population size. However, in natural, geographically-spread populations there can be gradients in environmental factors influencing reproductive success that can align with gradients in allele frequencies generating drift, these drift terms are not merely sampling noise and so will not approach zero as sample sizes become very large.

### S1.3 Contribution of Mendelian Segregation

The term 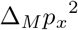 is the squared allele frequency change due to Mendelian segregation from heterozygote parents to their offspring, and, in expectation, corresponds to the variance in allele frequency change due to Mendelian segregation. In principal, if we observe genotypes in the offspring generation, we can observe the alleles transmitted out of each heterozygote and so directly calculate the contribution of Mendelian segregation averaged over markers (see e.g. Chen *et al*., 2019). However, in practice, we often will not have access to the offspring’s genotypes, but we can replace the average with its expectation. From our definition of Mendelian segregation in Eq. S6, we can derive

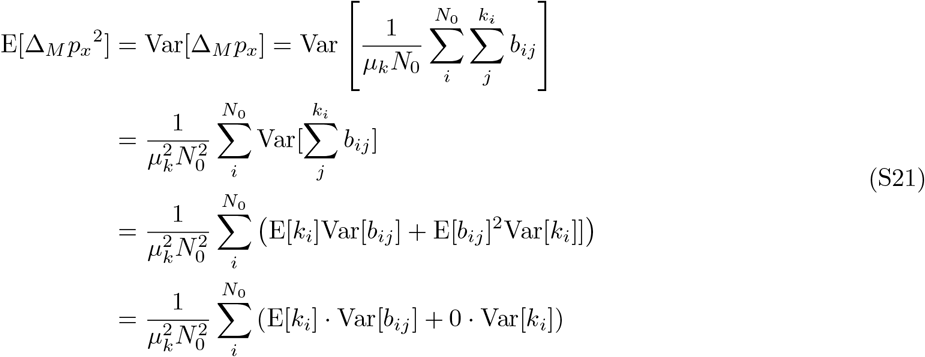

where the third and fourth lines follow from the total law of variance for a conditional variable such that

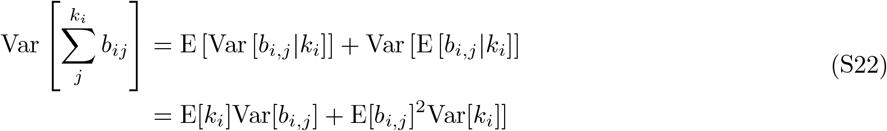

and considering 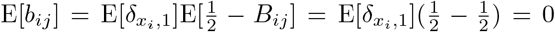. Note that E[*k*_*i*_] = E[*µ*_*k*_*f*_*i*_] = *µ*_*k*_, given that E[*f*_*i*_] = 1, and

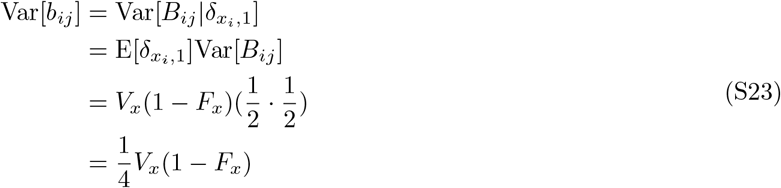

where 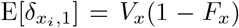 corresponds to the expected proportion of heterozygotes in the parental cohort, *F*_*x*_ being the inbreeding coefficient at locus *x*. Then we can write Eq. S21, the expected contribution of Mendelian segregation at a locus as:

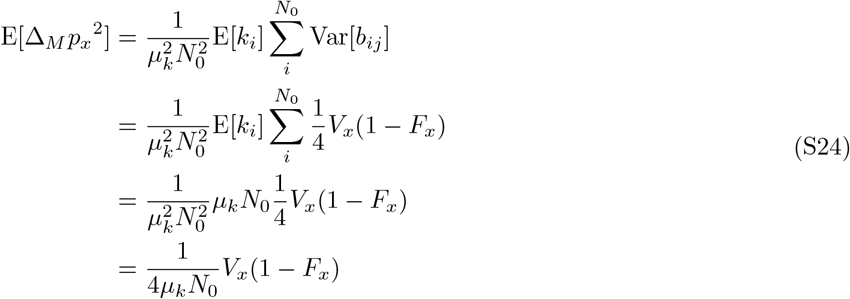

This expression is useful because we can still calculate the expected contribution of Mendelian segregation when we do not have access to the genotypes in the offspring cohort.

## S2 LD Score Regression

### S2.1 Full expression

Using Eq. S18, S19 and S21, we can write the full expression for the expected squared allele frequency change as:

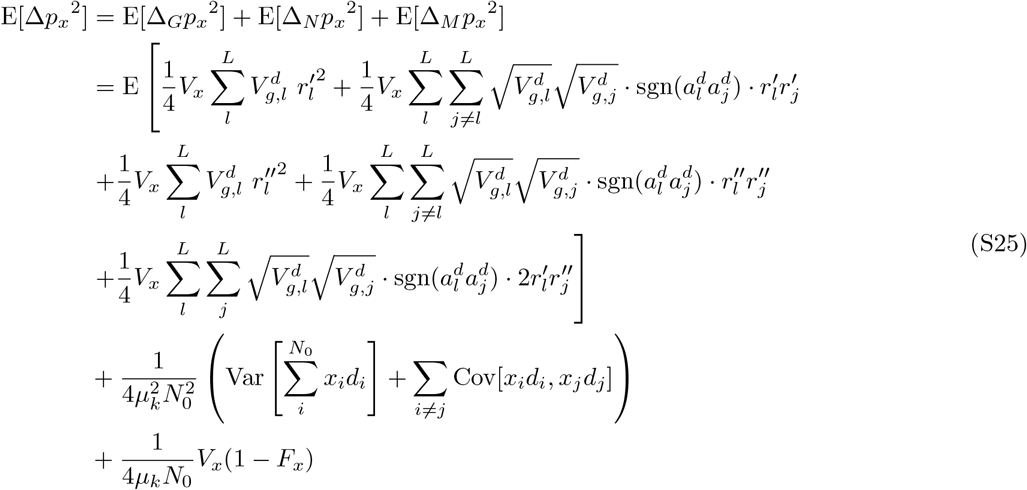

It is useful to normalize Δ*p*_*x*_^2^ at a locus *x* by its genotypic variance *V*_*x*_(1 + *F*_*x*_), to avoid correlations due to scaling of *r*_*l*_ and Δ*p*_*x*_^2^, and to simplify the expression for the contribution of genetic selection (first term) assuming that *r*_*l*_ is not correlated with the additive genic variance at a selected site 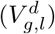. Under this assumption, we can decompose 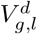 into its mean and a deviation term, 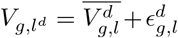, and if the deviation 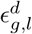 is small, it follows that 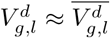. If we assume no covariance between genotype *x* and the random deviation in offspring number *d*, this becomes:

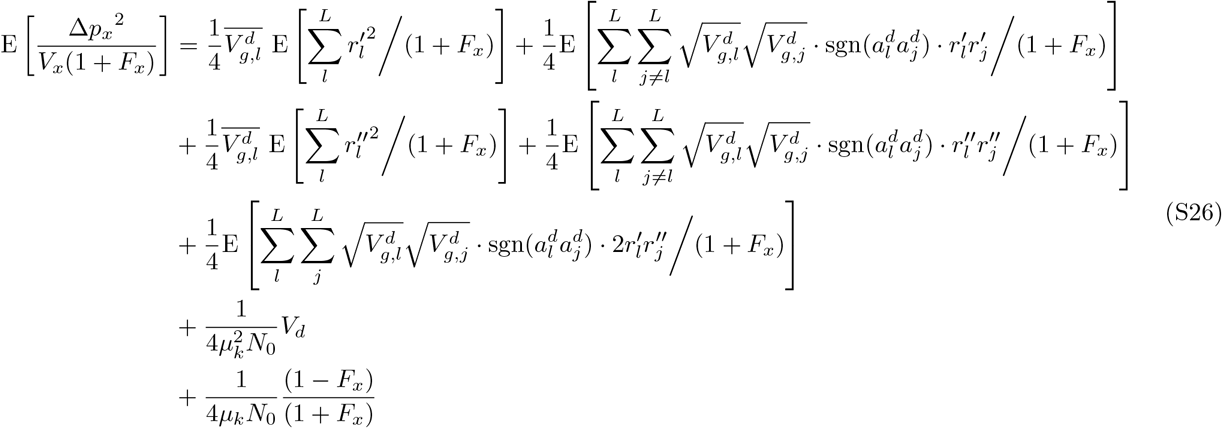

This expression can be further simplified if we assume that i) the trans-LD terms 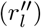 and the cross-product of cis-LD terms 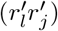 are negligible; ii) the inbreeding coefficient is negligible (*F*_*x*_ ≈ 0); iii) the population size has remained constant over generations (*µ*_*k*_ = 2); and iv) the offspring number of each individual follows a Poisson distribution (*V*_*d*_ = *µ*_*k*_ = 2). Then

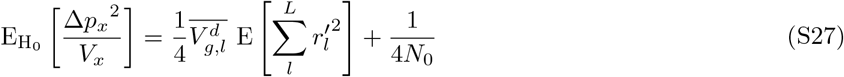

This is a very similar equation to that derived by Bulik-Sullivan *et al*. (2015) for the LD score regression on the GWAS chi-squared statistics with no environmental or genetic confounding (up to their scaling by the sample size). Their assumption of no environmental confounding in the GWAS is equivalent to our assumption that there is no covariance between genotype *x* and the random deviation in offspring number *d*.

According to Bulik-Sullivan *et al*. (2015), the LD score measured in a sample deviates from the true LD score in a population as follows 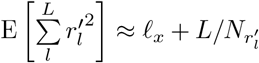, where *ℓ*_*x*_ indicates the true LD score, *L* the number of loci and 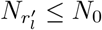 the sample size used to calculate the sample LD score. Therefore Eq. S28 becomes:

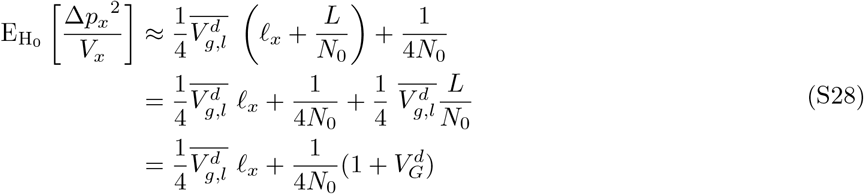

where 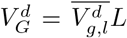 is the average total additive genic variance. Note that the second term of this expression, in expectation, is no longer dependent only on the sample size *N*_0_, but is inflated by the total additive genetic variance 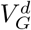. This means that the full expression in Eq. S26 will also be inflated by a similar factor.

### S2.2 Population Structure and Environmental confounding

In this section, we work through a simple model to illustrate the relationship between the LD score regression slope, genetic differentiation and environmental stratification. The estimates of genetic contributions from the slope of the LD score regression is generally considered to be free of environmental stratification, however, this can break down in some cases. To explore this, we follow Bulik-Sullivan *et al*. (2015) and model population structure by considering a sample of individuals 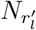, where 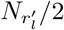 are sampled from population 1 (*P*_1_) and 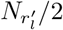 from population 2 (*P*_2_).

#### Genetic Differentiation

Genetic differentiation between populations 1 and 2 is exclusively driven by genetic drift (i.e., genetic stratification). Thus, given a focal locus *x*, E[*x*|*P*_1_] = E[*x*] + *f*_*x*_ and E[*x*|*P*_2_] = E[*x*] − *f*_*x*_, where *f*_*x*_ is the drift term at locus *x*, which is sampled from a normal distribution *f*_*x*_ ∼ *N* (0, *F*_*ST*_ **V**), where **V** is a correlation matrix across loci and *F*_*ST*_ is the Wright’s *F*_*ST*_, and E[*x*] is the expected genotypic value in the *N*_*r*_*′* sample. Both **V** and E[*x*] refer to the ancestral genetic composition before the splitting of the two populations. Therefore, under a polygenic model, the sample LD score is given by Bulik-Sullivan *et al*. (2015):

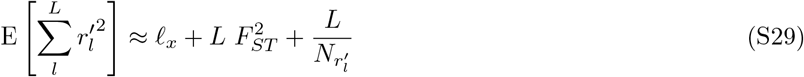

This expression shows that the expected LD score in a sample drawn from a mixture of populations reflects the true LD score, inflated by the degree of genetic differentiation between them 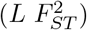. Since *F*_*ST*_ is assumed to be constant across loci, there is no correlation between *ℓ*_*x*_ and *F*_*ST*_ .

#### Environmental Stratification

The “number of offspring” phenotype (*k*_*i*_) is also influenced by population structure via environmental stratification, such that:

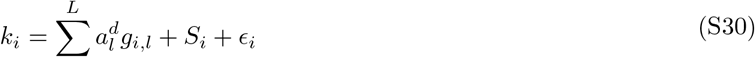

where *S*_*i*_ is an environmental stratification term defined by 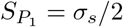 for *i* ∈ *P*_1_ and 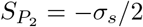 for *i* ∈ *P*_2_. By defining 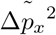 as the squared allele frequency change normalized by the observed genotypic variance in the sample *V*_*x*_(1 + *F*_*x*_ + *F*_*ST*_), then:

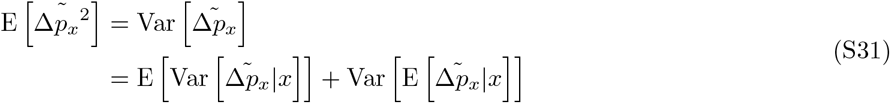

which follows from the law of total variance and decompose the within-sample and between-samples effects. Note that the conditioning on *x* means conditioning on the genotypes of the mixed sample and that the expectation is calculated over various realizations of the evolutionary process. For simplicity, and in agreement with the cohort sample considered in the present study, we assume that *F*_*x*_ and *F*_*ST*_ are small enough such that (1 + *F*_*x*_ + *F*_*ST*_) ≈ 1. Taking the assumption that *x* and *d* are independent, the first term is given by the first line of Eq. S26, since by conditioning on *x* the only source of randomness is given by the genotypes at the selected loci:

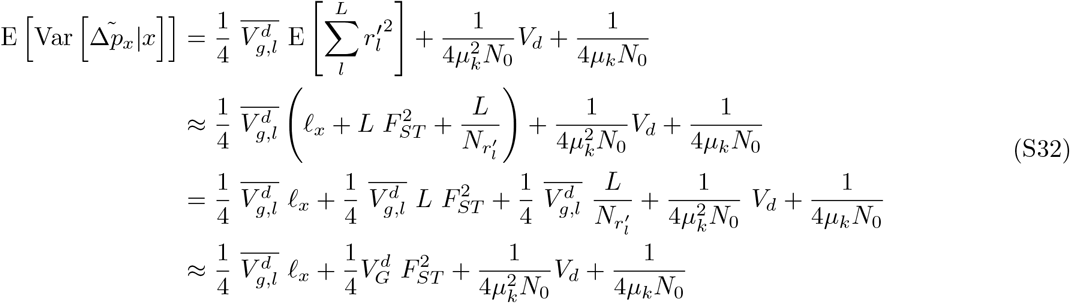

where 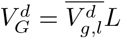 and 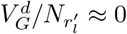, given that additive genic variance is typically very small and cohort sample size very large. To derive the second term, we first need to define:

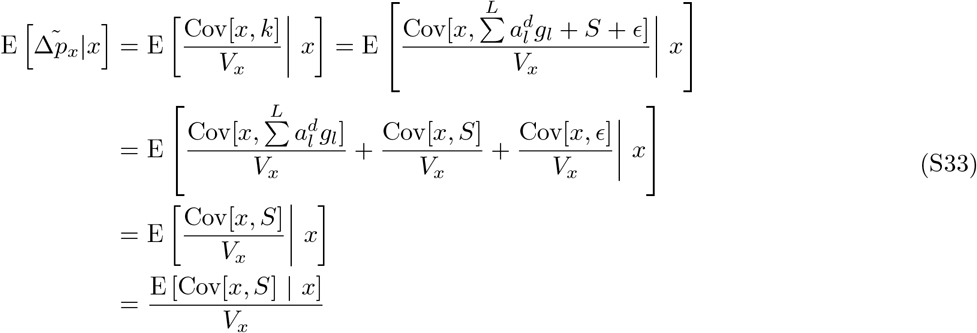

where the third line considers 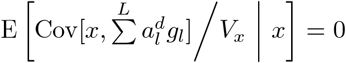, given that 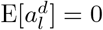, and 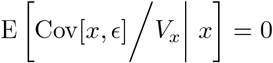 since the genotype *x* does not covary with noise *ϵ*. Moreover, in the last line we show that *V*_*x*_ can be moved outside the expectation, as it is constant conditional on the sampled genotypes being known. Given that *S* is a deterministic quantity, conditioning on *x* and using the law of total covariance lead to:

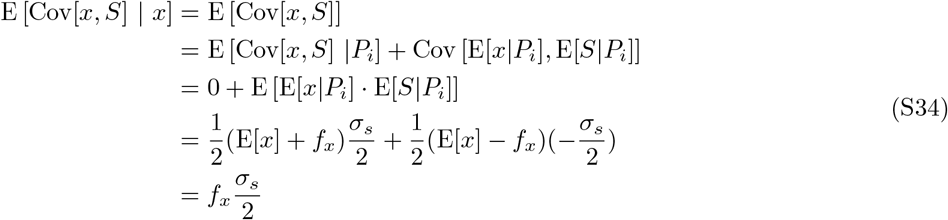

then

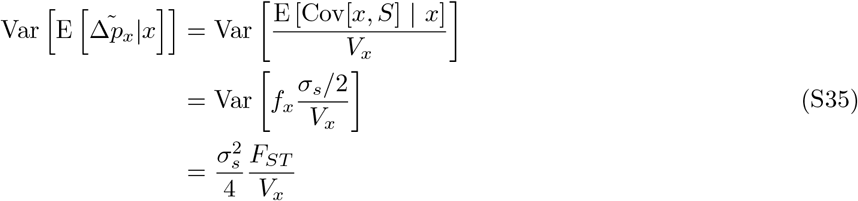

therefore,

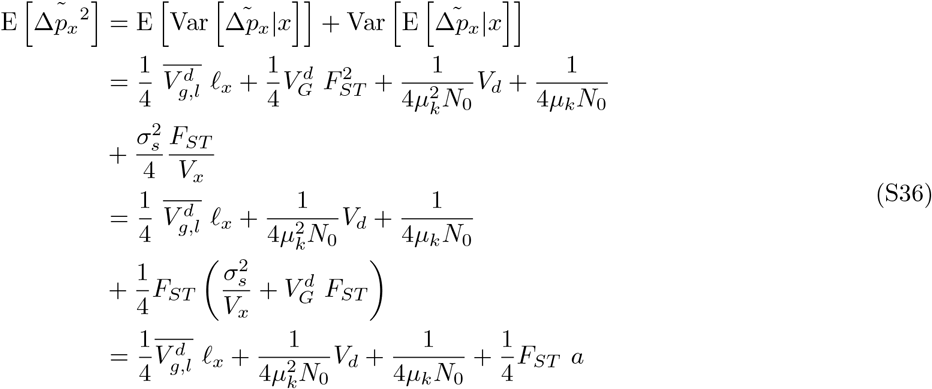

Note that 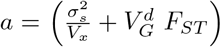 is related to the expected squared mean difference in number of offspring between population 1 and 2, with environmental stratification 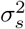 and genetic differentiation 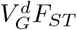. In fact,

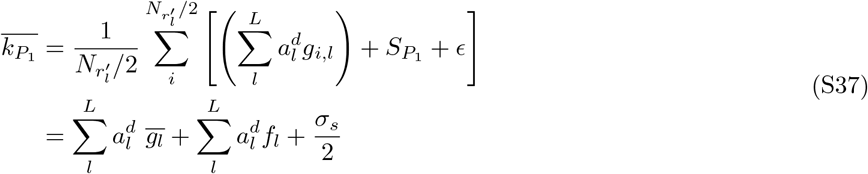

where 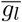 is the expected genotype at locus *l* in the ancestral population. Similarly, 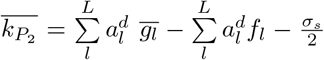 and

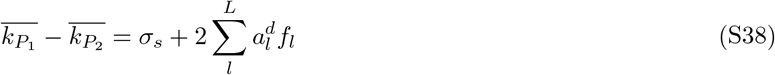

then we have

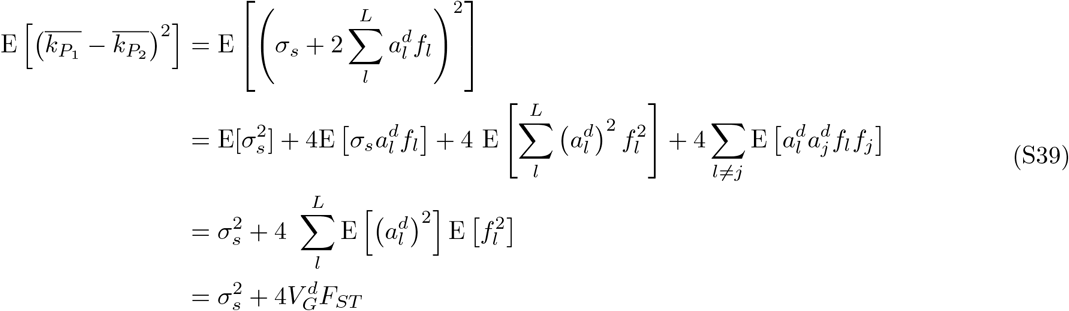

where in the third line we note that 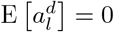, the effect size 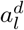 is independent from the drift term *f*_*l*_ and that the 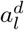 are independent and identically distributed. As noted by Berg *et al*. (2019), population structure may not affect the genome uniformly, in the sense that factors such as variation in recombination rate and background selection can drive correlations between LD score *ℓ*_*x*_ and *F*_*ST,x*_ along the genome. Following Berg *et al*. (2019), we show that these correlations are eventually absorbed by the slope of the regression (Δ*p*_*x*_^2^*/V*_*x*_) ∼ *ℓ*_*x*_, biasing the estimate of additive genic variance 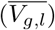. Under these conditions, the slope is given by:

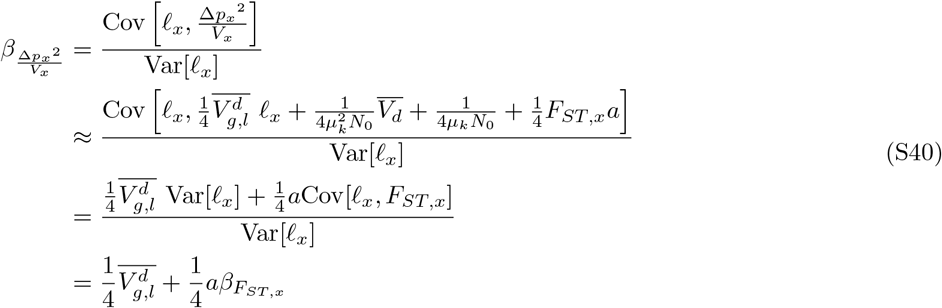

where 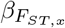 is the slope of *F*_*ST,x*_ regressed on LD score *ℓ*_*x*_. The intercept can be derived as:

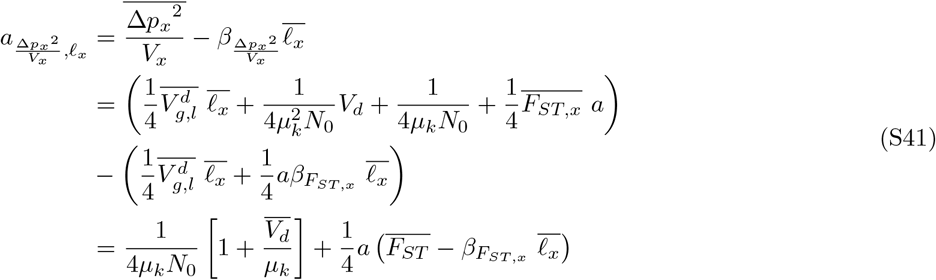

This is useful, for instance, when a linear spatial gradient in genotype (e.g., from a recent expansion) covaries with a linear spatial gradient in the environmental effect on offspring number. The steepness of this environmental gradient, the parameter *a*, which sets how strongly the phenotype is differentiated along it, determines how much the LD score regression slope is confounded.

### S2.3 Bulmer-effect and Assortative Mating

In deriving the expressions for the LD score regression we assumed that there was no systematic long-distance LD between the focal variant and causal variants elsewhere in the genome. However, this assumption will be violated by the Bulmer effect and assortative mating, altering the interpretation of the slope of LD score regression (Border *et al*., 022a; Veller *et al*., 2024). Here we explore these effects and how they play into our allele frequency change framework. The assumption about systematic long-distance LD comes in the expression for the contribution of genetic selection to allele frequency change (Eq. S18), at the second, third, fourth, and fifth terms of that equation. This systematic signed LD arises from a variety sources, two common ones being the Bulmer effect and assortative mating.

#### Bulmer-effect

As our effect sizes reflect fitness itself, we expect that if they have been subject to similar selection for multiple generations, there should be weak negative LD between pairs of selected loci (the Bulmer effect Bulmer, 1971, 1974). This acts to reduce the effect of selection at the focal site (e.g. Negm and Veller, 2026). This is quantifiable in the second, fourth, and fifth terms of Eq. S18, which involves the product of the pairwise LDs between the focal site *x* and the selected sites *l* and *j*, weighted by the square of their additive genic variance. However, note that if the sign of LD was random with respect to the sign of the effect sizes of the selected sites *l* and *j*, these terms would be zero in expectation.

We quantify the effects of this long distance LD mathematically using the “omitted-variable bias” approach. More in general, consider the case in which a response variable *Y* is generated by a process that can be modeled as

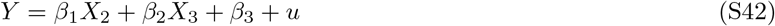

however, imagine that we do have measures only for the predictor *X*_2_ and thus decide to model *Y* as

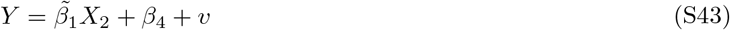

the estimated slope 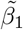 would then be

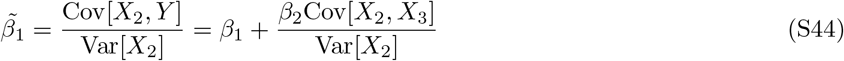

Concerning the squared allele frequency change in Eq. S18 (assuming that *F*_*x*_ ≈ 0) and focusing on the second term, which is expected to have relatively higher effect, this would lead to

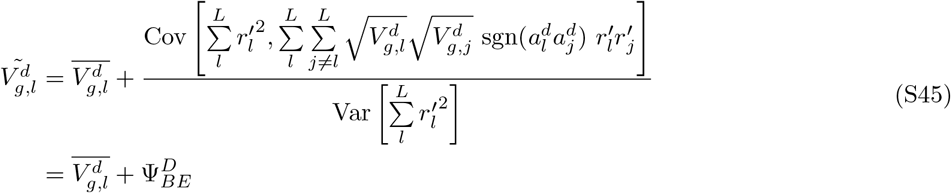

In Eq. S45, we define the bias in the estimated average additive genic variance 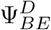, which is expected to be negative since Bulmer effect induces negative pairwise LDs between selected sites. This allows us to model the reduction on the impact of linked selection due to the Bulmer effect as:

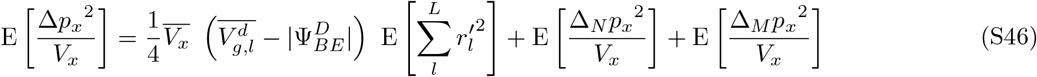

thus the Bulmer effect decreases the slope of the LD score regression. This leads to a bias in the slope estimated, as compared to that expected under the additive genic variance with no systematic LD. However, from our perspective the slope accurately captures the reduced allele frequency change due to selection due to the Bulmer effect.

#### Assortative Mating

We can think of assortative mating as acting on the third term of Eq. S18, where the squared of the trans-LD between the focal site *x* and the selected site *l* is considered. Then, similarly to Eq. S45, we can quantify the upward bias on 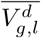 induced by assortative mating as

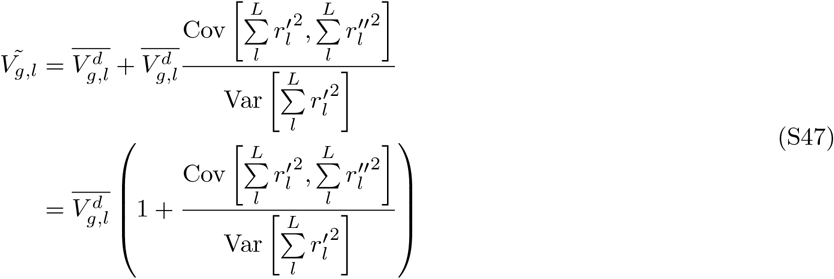

Then, we can define 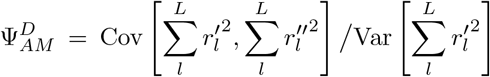 as the upward bias in the estimated average additive genic variance due to assortative mating. This allows us to model the increase in the impact of linked selection to the squared allele frequency change due to the assortative mating as:

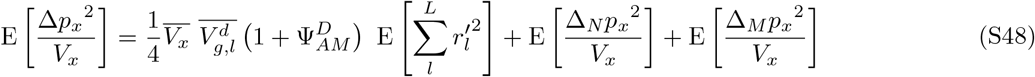

Thus assortative mating upwardly biases the LD score regression slope as compared to the genic additive variance expected with no systematic LD. However, again from our perspective, this inflation of the slope is part of the effect of assortative mating on allele frequency change.

### S2.4 Indirect Parental Genetic Effects

In the sections above, we have considered the model *k*_*i*_ = *µ*_*k*_*f*_*i*_ + *d*_*i*_, where the observed number of children of an individual *i* is composed by their expected contribution due to their genotype *f*_*i*_ and a deviation due to sampling noise or environmental factors *d*_*i*_. However, we can construct a more comprehensive model considering that the expected genetic contribution *f*_*i*_ should also take into account the indirect parental effects 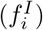, dependent on the genotypes of their parents. Accordingly, Eq. S5 can be re-written as:

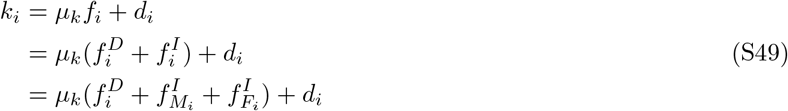

where 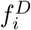 is the direct genetic effect and 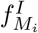 and 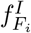 are the maternal and paternal indirect genetic effects of the focal individual. In this general additive model, the number of children of an individual *i* can be expressed as:

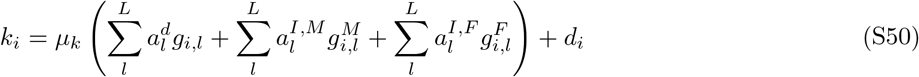

where 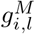 and 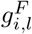 are the genotypes of the mother and father, respectively, of the individual *i* at the locus *l*. In Eq. S11, we have derived the genetic contribution to the allele frequency change by considering only direct genetic effects. We now also consider the indirect parental genetic effects.

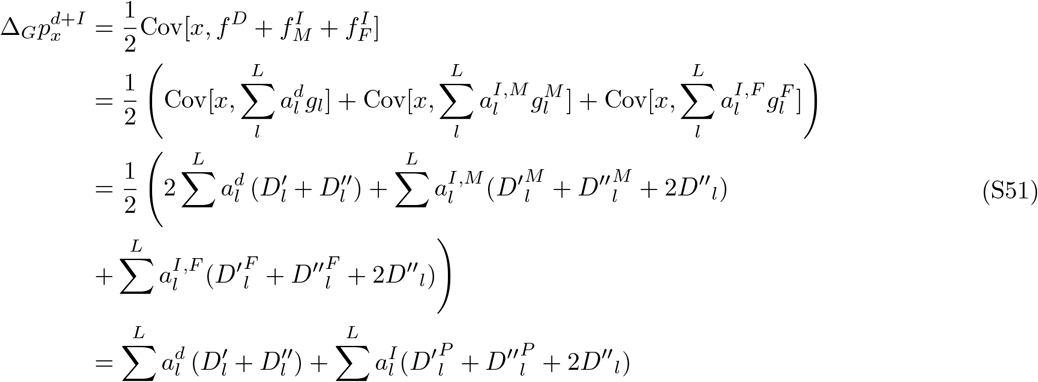

where the last line assumes equality between indirect paternal and maternal effects, 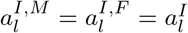, and equality in paternal and maternal cis-LD, 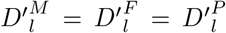 where *P* stands for parental population. The genotypic covariance between the focal locus *x* and the indirect genetic effect in Eq. S51 is derived by considering that each individual has received one of the two alleles at a locus from each of the two parents. Therefore,

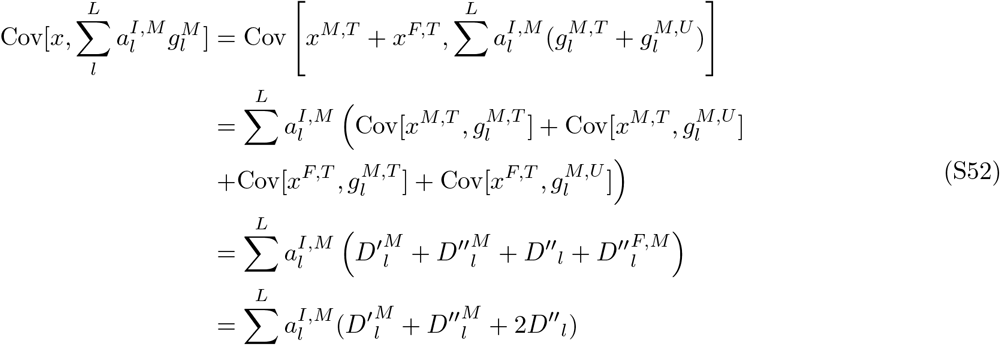

where *T* and *U* indicate alleles that have been transmitted or untransmitted from the parents, 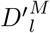 and 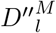 are the cis- and trans-LD in the mother, *D*″_*l*_ is the trans-LD in the focal population and 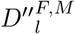 is the trans-LD between the parents of the focal population. In the last line, we have considered that 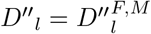, meaning that the trans-LD between parents is equal to the trans-LD in the focal population. We then write the squared of the allele frequency change as:

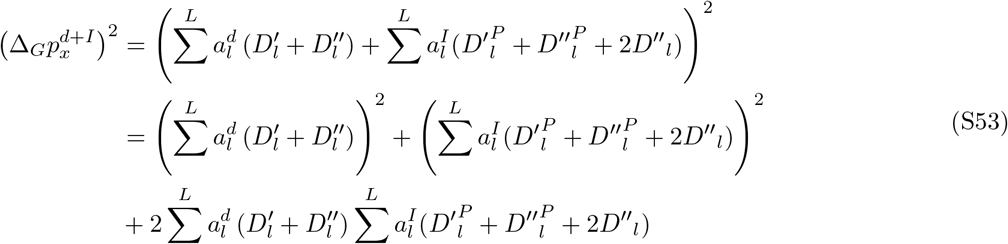

the first term has been derived in Eq. S12-S18, while the second term is given by:

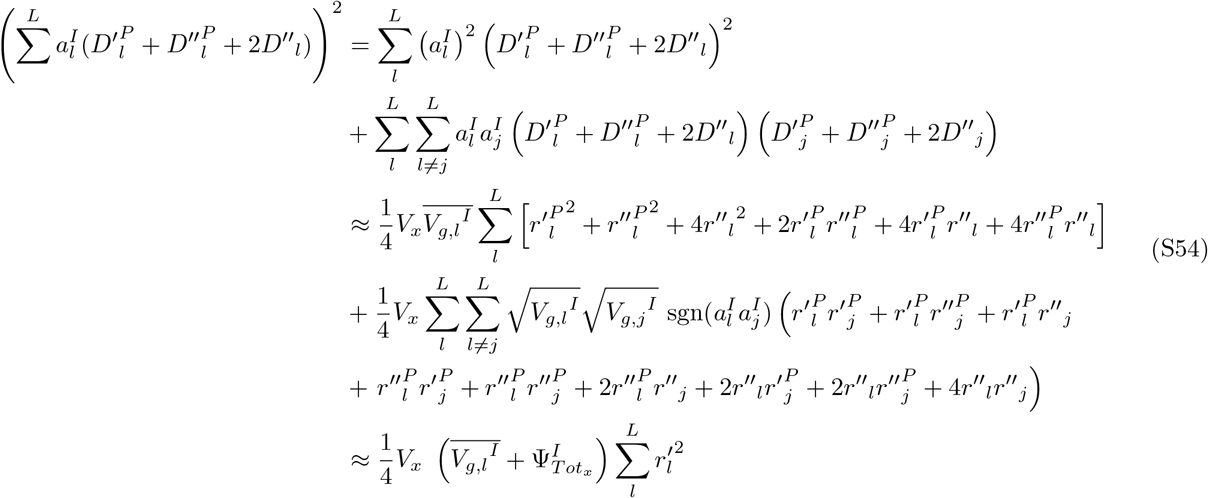

where 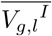 is the average additive genic variance of the indirect genetic effects and 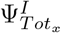 is the bias due to Bulmer-effect and assortative mating similar to Eq. S45. Note that the first approximation assumes 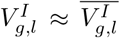, while the second relies on the fact that, since 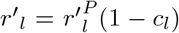, where *c*_*l*_ is the recombination rate, and *c*_*l*_ ≈ 0 as LD is typically estimated among nearby loci, then it follows 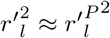. The third term of Eq. S53 is given by:

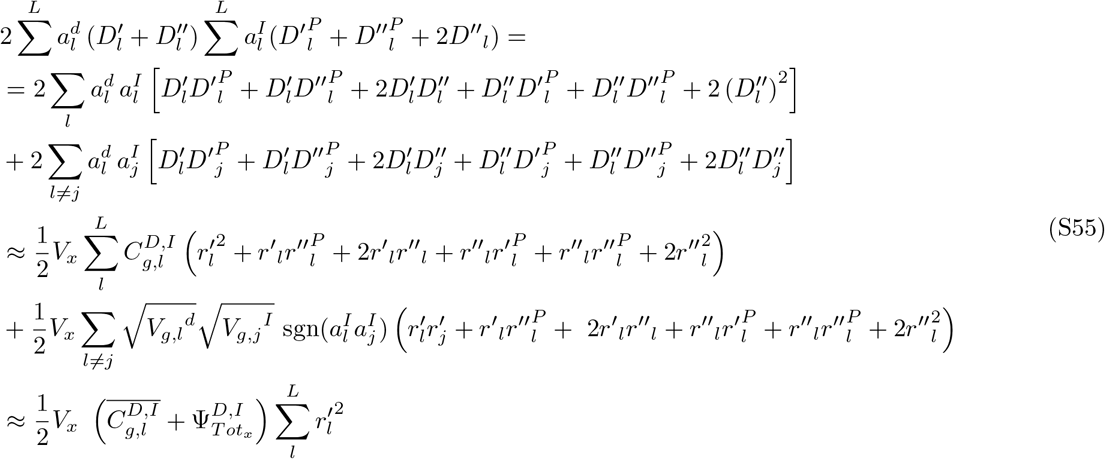

where 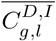 is the average additive genic covariance between direct and indirect genetic effects and 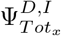 is the bias due to Bulmer-effect and assortative mating. Then we can re-write Eq. S53 as:

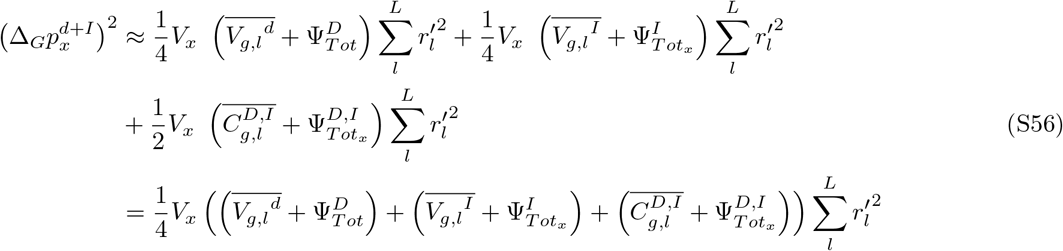

This shows that LD score regression on squared allele frequency change, after dividing by genotypic variance, yields estimates of average additive genic variance that are biased by indirect genetic effects, covariance between direct and indirect genetic effects, Bulmer-effect and assortative mating.

## S3 Estimating Allele Frequency Change from GWAS effect size for fitness

It is natural to normalize the squared allele frequency change (Eq. S4) by the observed variance in genotype across individuals *V*_*x*_(1 + *F*_*x*_), which scales both the effects of drift and selection at the locus. This scaled change can be written as

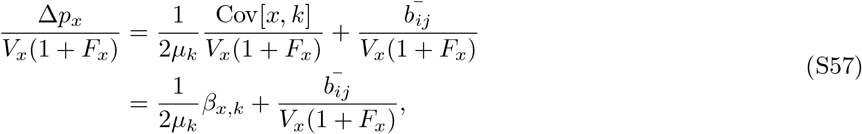

where *β*_*x,k*_ corresponds to the effect size of the focal site *x* in a GWAS for the number of offspring *k*. Noting that *k*_*i*_ = *µ*_*k*_*f*_*i*_ + *d*_*i*_, the above can be expanded as

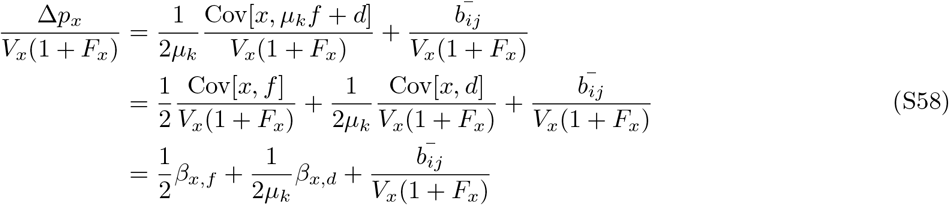

where *β*_*x,f*_ is the slope of the regression of the genetic (breeding) value *f* on individual genotype, and *β*_*x,d*_ is the slope of the regression of the environmental contribution to offspring number *d* on individual genotype. This means that we can estimate the squared allele frequency change over a single generation, by using GWAS effect sizes on the number of children such that

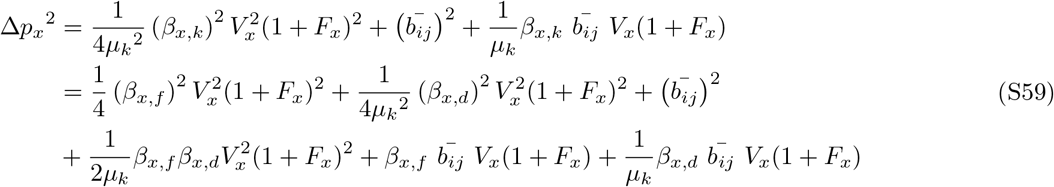

Assuming no covariance between the effect sizes for fitness and the effect sizes for the environmental contribution (*H*_0_), and taking the expectation of the normalized squared allele frequency change leads to:

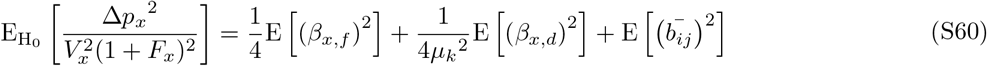

Note that in Eq. S59 we considered a phenotypic model that includes only direct genetic effects and the environment *k*_*i*_ = *µ*_*k*_*f*_*i*_ + *d*_*i*_. In the following sections, we extend this to a general model as in Eq. S49, by also incorporating the direct and indirect genetic effects of the partner(s) of the focal individual. This extension is necessary as unlike other individual-level phenotypes (say height) the phenotype ‘number of children’ manifestly represents the combined contribution of both members of a couple, which will have consequences for the interpretation of population and family-based GWAS. Under this model, we obtain:

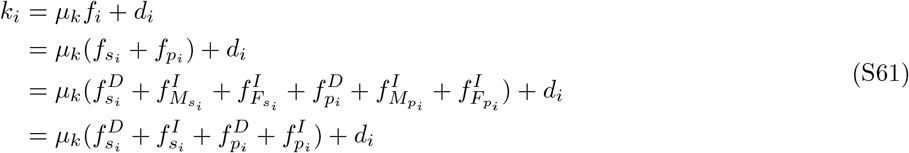

where 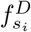 and 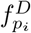 are the direct genetic effects of the focal individual and the partner, respectively; 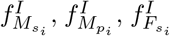 and 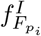 are the maternal and paternal indirect genetic effects of the focal individual and the partner, respectively; and *d*_*i*_ is the family sampling noise contribution. Note that in the case of an individual having multiple partners, the partner effects could be considered as the average across those partners. In the following subsections, we show how direct and indirect genetic effects compose the effect sizes estimated in a population and sib-GWAS.

### S3.1 Population GWAS effect sizes

In a population GWAS, we fit the model:

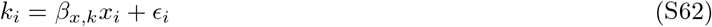

where *i* indicates the *i*th individual. Following Eq. S61, we can then write the estimated effect size of the focal locus *x* as:

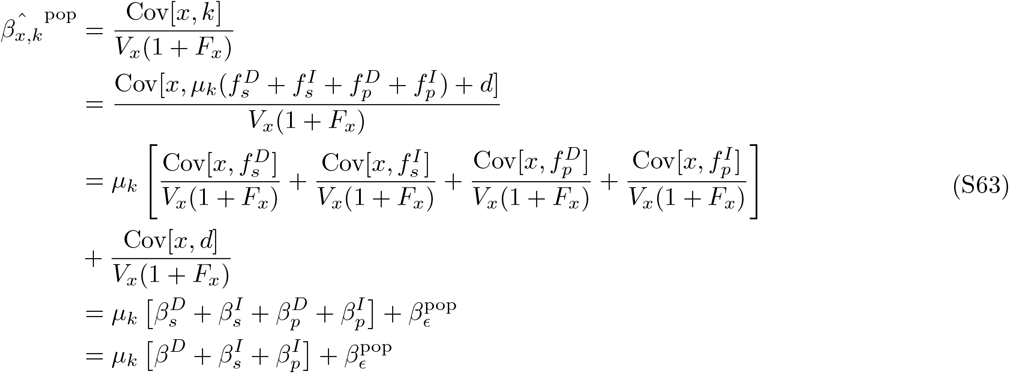

where the last line considers that the effect size of the direct genetic effect is the combined contribution of the focal individual and their partner(s) 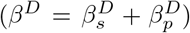. By taking the expectation of the squared effect size, we then obtain:

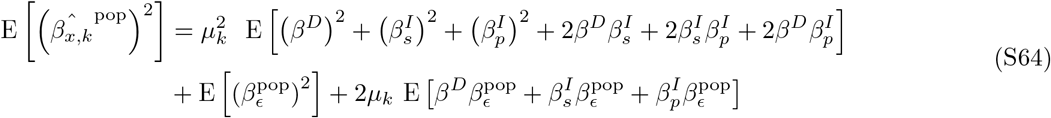

In the case where we could assume that there is no covariance between each of the effect sizes (*H*_0_), this expression simplifies to:

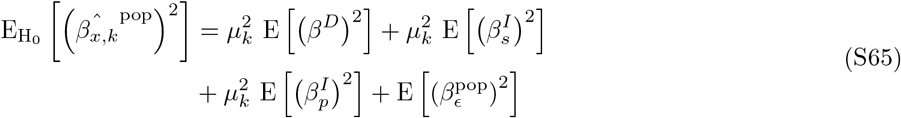

We can use the previous expressions to uncover the relationship between effect-size and LD score. Then, from Eq. S64, we can write

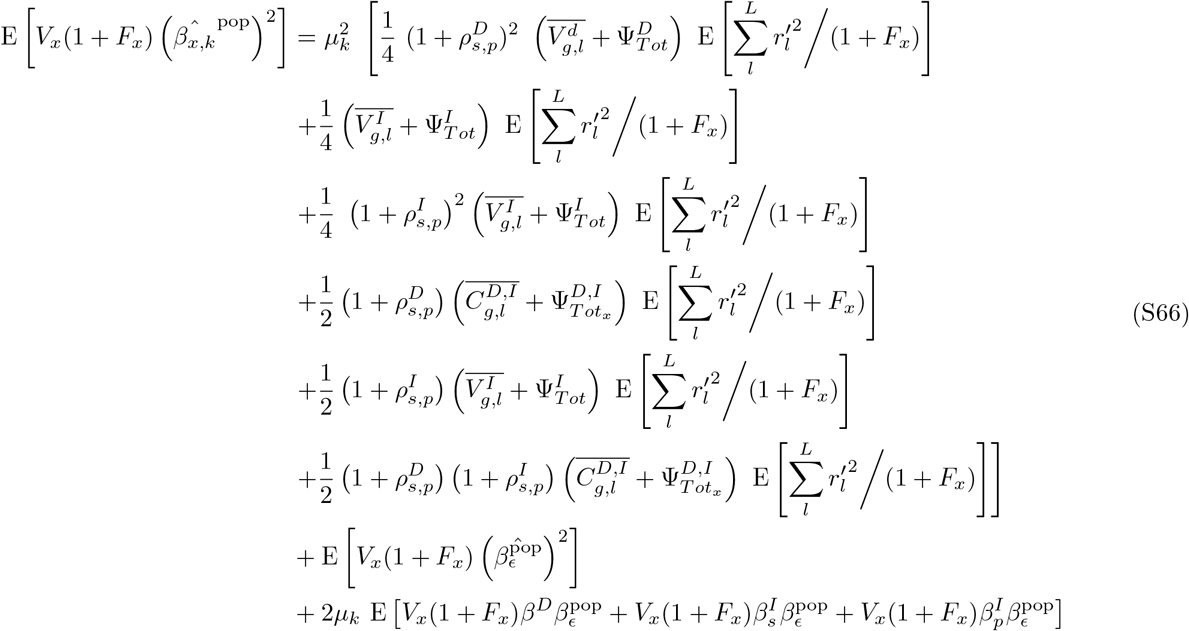

we can simplify this expression by defining 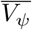 as the confounded additive genic variance, including Bulmer-effect, assortative mating and indirect genetic effects of the focal individuals and their partners:

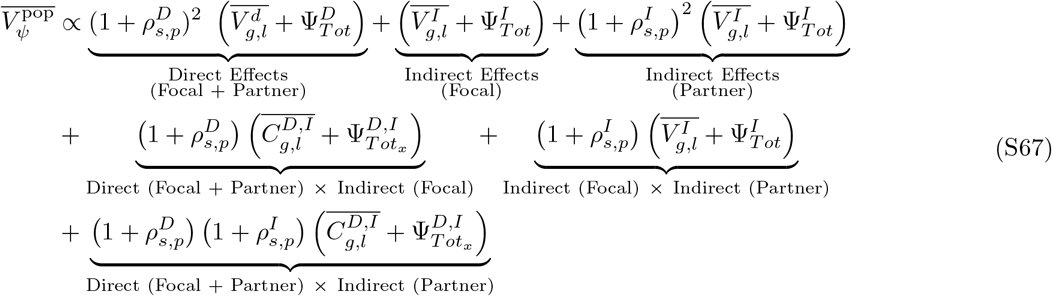

such that

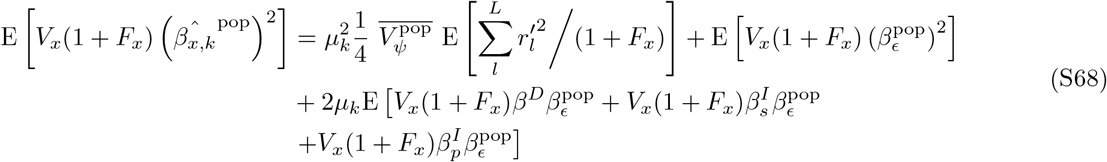

which shows that the LD score regression performed on the population GWAS effect-sizes for the number of children returns biased estimates of average additive genic variance per variant. Note that the estimated slope would be further biased upward if there exists positive covariance between direct or indirect genetic effects 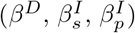 and environmental effects (*β*_*ϵ*_).

### S3.2 Family GWAS effect sizes

Sibling-GWAS, and family studies more broadly, are often used to mitigate various forms of environmental and genetic confounding (Young *et al*., 2019; Howe *et al*., 2022; Veller and Coop, 2024). In this section, we build on previous work but incorporate the effects of the partner(s). We assume that the siblings are sampled in the same ways as the individuals used in the population GWAS. We further assume that the genotypes at our focal locus of interest are randomly distributed over any interacting environments or genotypes elsewhere in the genome. These two assumptions ensure that the direct effect sizes have the same interpretation in both the sib- and population-based GWAS. For simplicity we consider a sibling-GWAS conducted by regressing the sibling difference in phenotype on their difference in genotype, i.e. fitting the model:

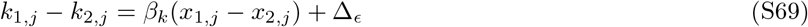

where *j* indicates the *j*^*th*^ family, 1 and 2 refer to the two siblings and Δ_*ϵ*_ is the difference in sampling noise between the two siblings. Then

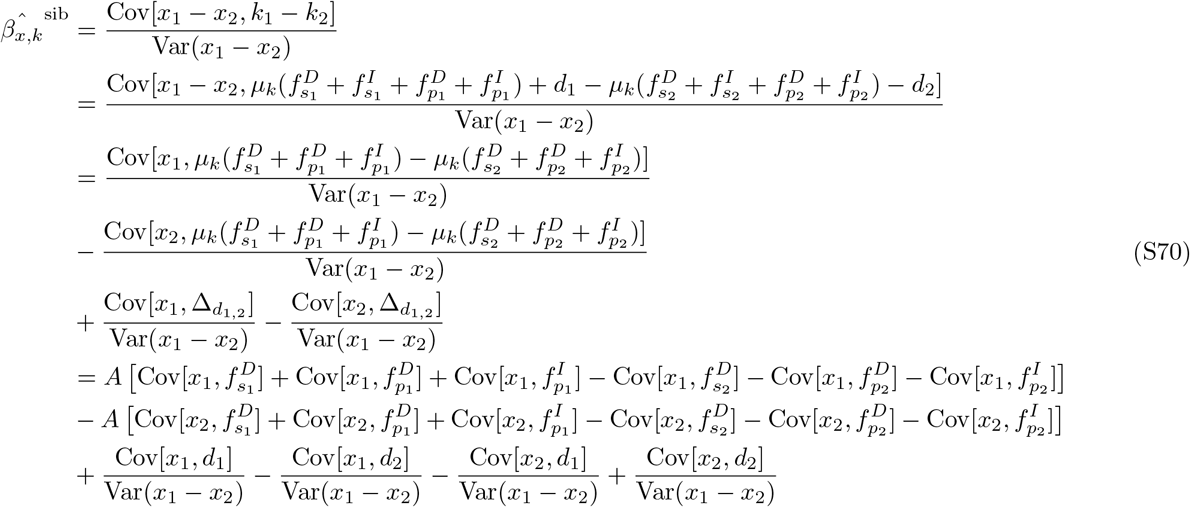

where 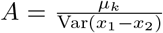. In the second line, the indirect genetic effects of the two siblings cancel out, since they are exposed to the same family indirect effects. Then Eq. S70 can be written in terms of effect-sizes as:

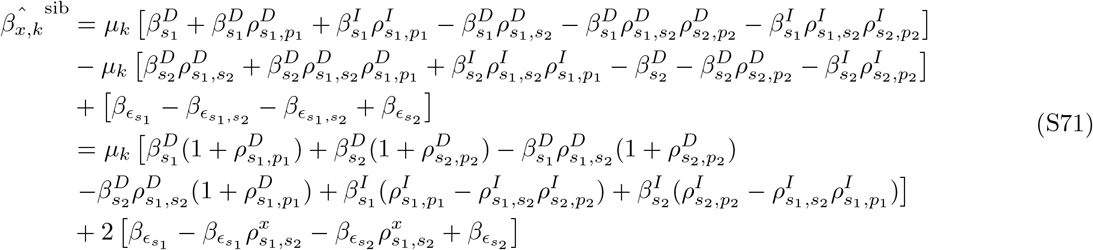

Note that 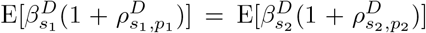 and corresponds to the measured direct genetic effect for the number of children in a GWAS (*β*^*D*^), which includes the direct genetic effects from both the focal individual and their partner(s). Moreover, several of the terms in Eq. S71 are derived by considering the general relationship *β*_*B*_ = *β*_*A*_*ρ*_*A,B*_, where *ρ*_*A,B*_ represents the correlation between *A* and *B*. Then

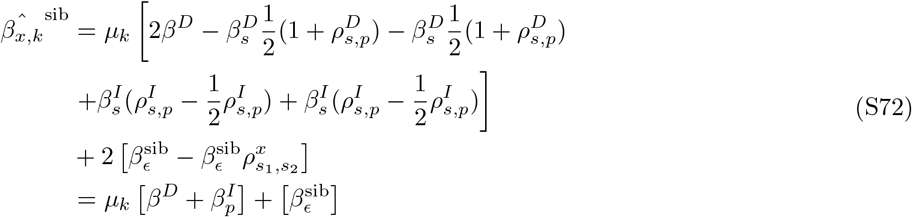

where 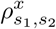 is the correlation in genotype between siblings and is equal to 0.5 in expectation, and 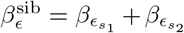. The expected squared effect size would then be

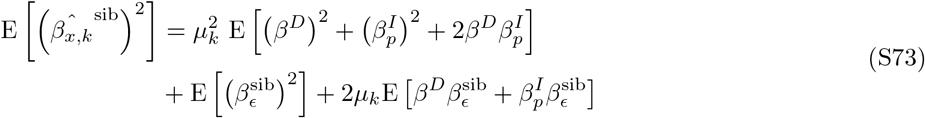

for a panmictic population (*H*_0_), this expression simplifies considerably:

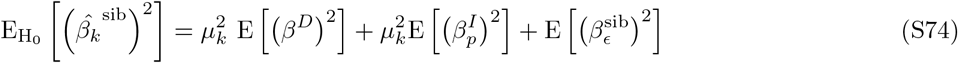

As in section S3.1, we uncover the relationship between effect-size and LD score and write

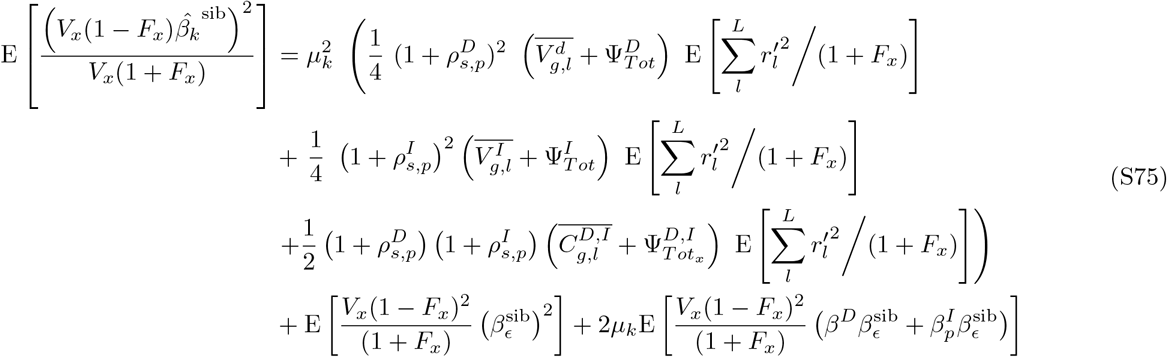

we can simplify this expression by defining 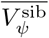 as the confounded additive genic variance, including Bulmer-effect, assortative mating and indirect genetic effects of the partners:

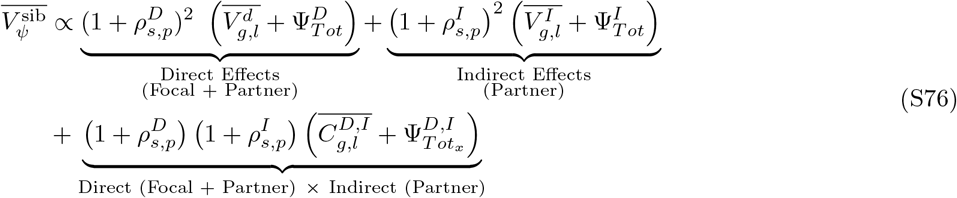

such that

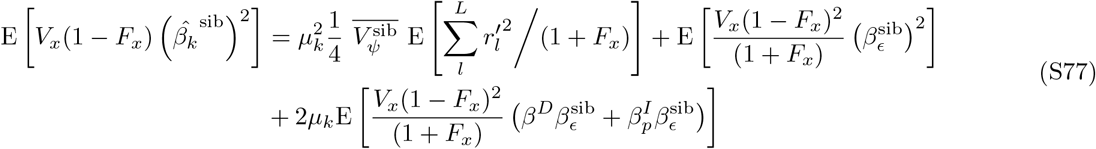

Thus, the intercept of LD score regression on sibs effect-sizes reflects sampling noise (second term in Eq. S77) while the slope absorbs the direct genetic effects of the focal individuals and their partner(s) (i.e., additive genic variance), the inderect effects of the partner(s) and the covariance in direct and indirect effects. As for Eq. S68 note that the estimated slope would be further biased upward if there exists positive covariance between direct or indirect genetic effects 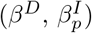 and environmental effects 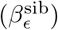.

### S3.3 Cross-Product Population and Family GWAS Effect-Sizes

In the main text we make use of the product of the sibling and population GWAS effect sizes. Here we derive the expectation of this quantity and its relationship with LD score regression. Noting that the denominator of a sibling GWAS effect size is given by 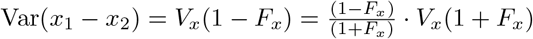, we can rewrite Eq. S72 in terms of population GWAS effect sizes, where the denominator is the observed genotypic variance instead of the frequency of heterozygotes.

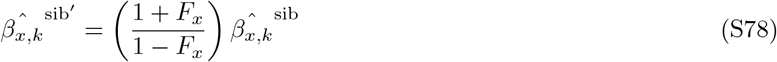

then we can

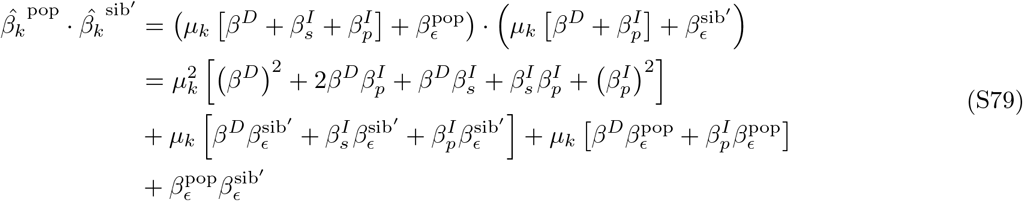

In expectation, this becomes:

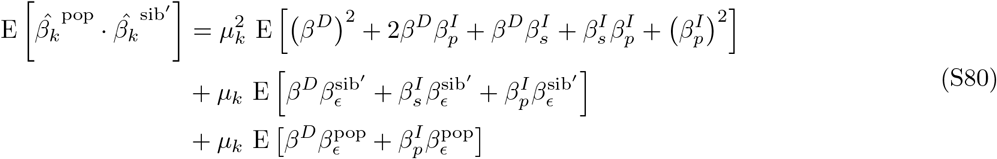

where the term 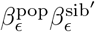 from Eq. S79 disappears since environmental effects of the population and sibling GWAS are uncorrelated, assuming that there are not overlapping individuals between these two cohorts. In the case in which there is no correlation between indirect, direct effects with environmental effects (*H*_0_), this expression simplifies to:

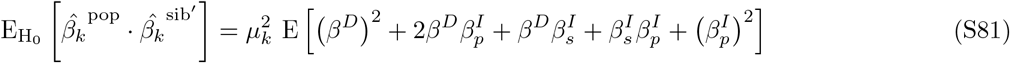

We can use Eq. S80 to uncover the relationship between effect-size and LD score:

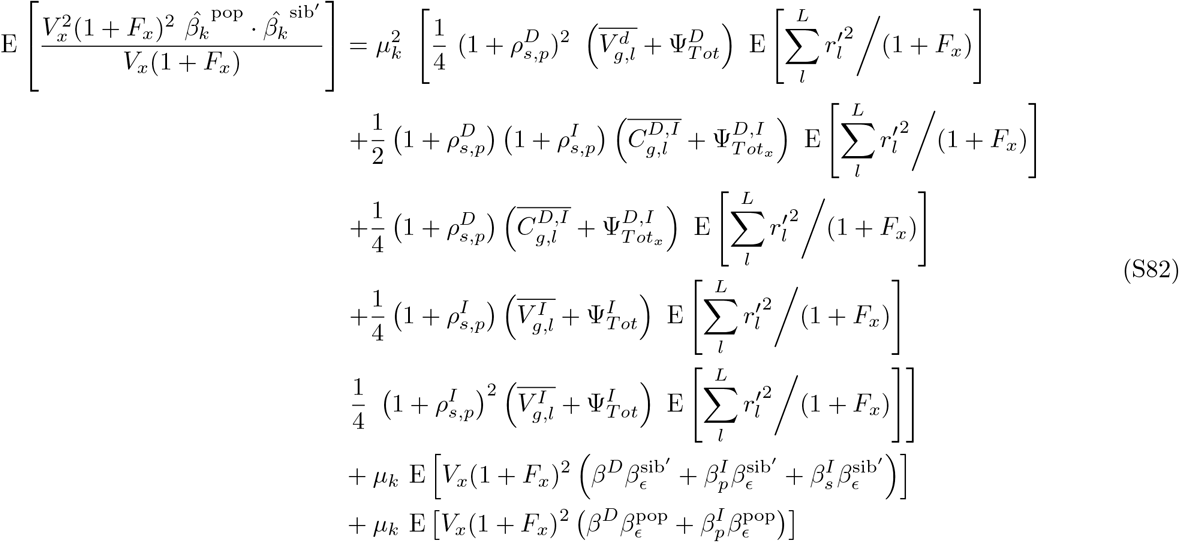

we can simplify this expression by defining 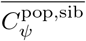 as the confounded additive genic covariance, including Bulmer-effect, assortative mating and indirect genetic effects of the focal individuals and their partners:

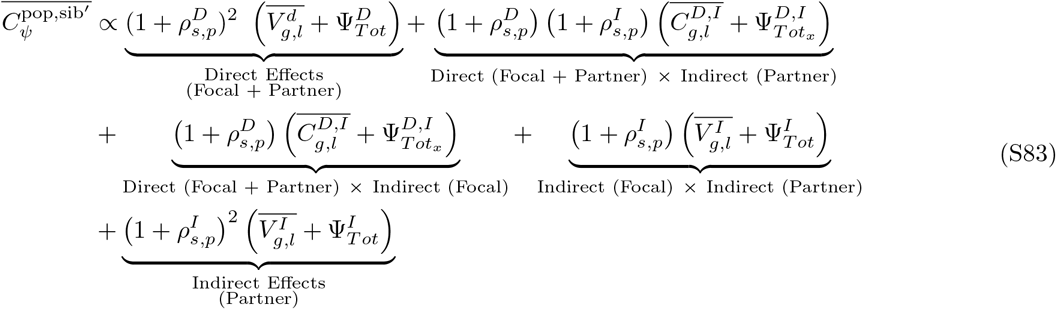

such that

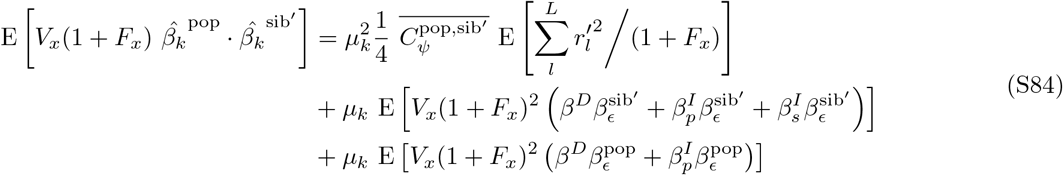

through all these equations the direct effects appear alongside the correlation with the indirect effects of the partner. The direct effects include the contribution of the focal individual and the correlation with their partner, through assortative mating.

## S4 *χ*^2^ and Squared Allele Frequency Change

In a typical GWAS analysis, LD score regression is performed against the *χ*^2^ statistics for each SNP. In this section, we revisit the link between *χ*^2^ and GWAS effect sizes *β*_*x,k*_ and, building on it, their connection to squared allele frequency change 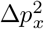. Under the hypothesis that *β*_*x,k*_ ≠ 0, the squared of the Wald’s test statistics 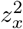 (used for Hypothesis testing) is asymptotically distributed as a non-central *χ*^2^ distribution with 1 degree of freedom and non-central parameter (*β*_*x,k*_*/SE*)^2^, such that:

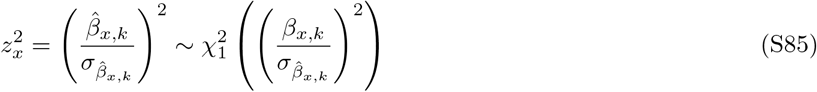

where 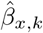 and 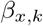 are the estimated and true effect sizes of site *x*, respectively, while 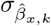 is the standard error of the estimator 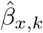:

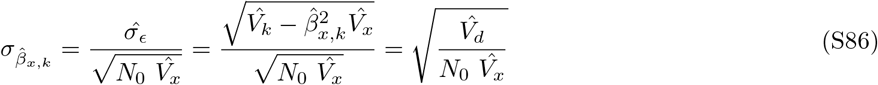

where 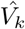 is the observed variance in reproductive success, 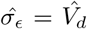 is the environmental variance in reproductive success, 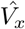 is the observed genotypic variance and *N*_0_ is the sample size. We can then define the non-central 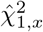 statistic as:

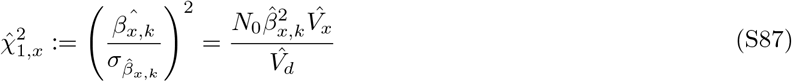

Following Eq. S59 and ignoring the Mendelian component to squared allele frequency change since it is not absorbed by the GWAS effect sizes, we can write 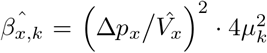 and then express 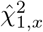 in terms of squared allele frequency change:

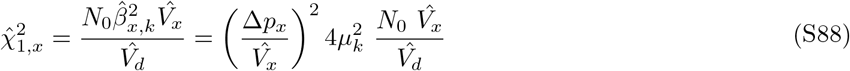

In Section S2, we showed that the intercept of the LD score regression on normalized squared allele frequency change, corresponding to the average environmental contribution to 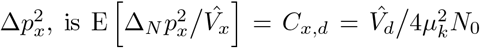 replacing this term in Eq. S88 and by taking the expectation, we obtain:

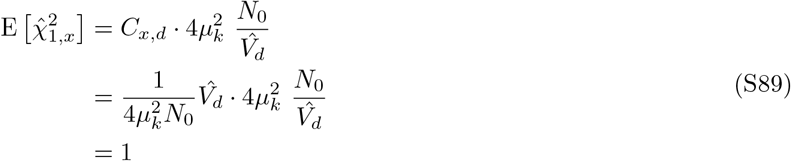

which corresponds to the expected intercept in a typical LD score regression on *χ*^2^ GWAS summary statistics in absence of population structure Bulik-Sullivan *et al*. (2015). With real data, 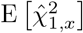 is likely to be *>* 1 both because of environmental stratification, genetic differentiation and because LD score on common alleles does not capture the contribution of rarer alleles, which will therefore be absorbed by the intercept, now defined as *C*_*x,R*+*d*_.

## S5 Deriving GWAS effect sizes on reproductive output from selection coefficients

Here, we summarize why selection coefficients are equivalent to the effect sizes obtained from a population GWAS of reproductive output over the complete life cycle. In particular, consider a model of allele frequency change under directional selection (Walsh and Lynch, 2018, Eq. 5.1c)

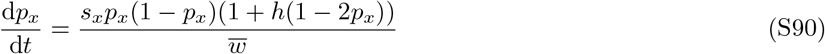

where *p*_*x*_ is the frequency of the randomly chosen allele at the focal site *x, s*_*x*_ is the selection coefficient, *h* is the dominance coefficient (*h* = 0 under an additive model) and 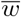 is the mean relative fitness, which is equal to 1. Then

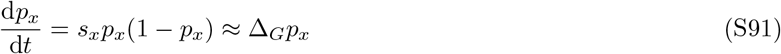

As shown in Eq. S4, 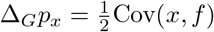, then 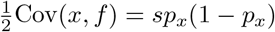 and

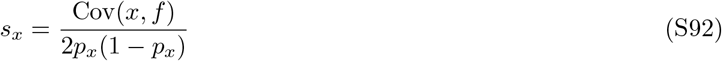

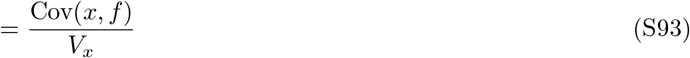

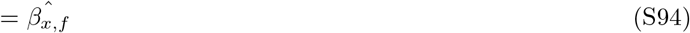

meaning that the selection coefficient *s*_*x*_ is equal to the effect size at site *x* in a population GWAS for the number of offspring.

## S6 The connection of genetic covariance of population- and sibling-based GWAS effects to the fundamental and secondary theorems of natural selection

The fundamental and secondary theorems of natural selection define the partial change due to selection in the population mean fitness and the change in a character correlated with fitness (Fisher, 1930; Price, 1970; Robertson, 1966). Both of these theorems define the change in terms of the additive genetic variance specified in the product of the ‘average excess’ and the ‘average effect’ of an allele (Fisher, 1930, 1941; Crow and Nagylaki, 1976). The average excess is interpreted as the regression coefficient of fitness on the additive genotype at a locus, i.e. the target of our population based GWAS. The average effect of an allele is interpreted as the causal effect of the allele on fitness, corresponding to a hypothetical manipulation experiment where one allele is swapped for the other at conception (Lee and Chow, 2013). The sibling GWAS can be interpreted as estimating the causal effect of the allele in the cohort if can assume that the siblings are representative of the WB samples as a whole, that there are no indirect sib effects, and that genotypes are randomly distributed across interacting backgrounds (Veller *et al*., 2024). The LD score regression of the product of the population- and sibling-based GWAS is estimating the genetic covariance of these two quantities, and thus the average of the ‘average excess’ and the ‘average effect’ of an allele across loci.

## 3 Supplementary Figures

**Figure S1:**
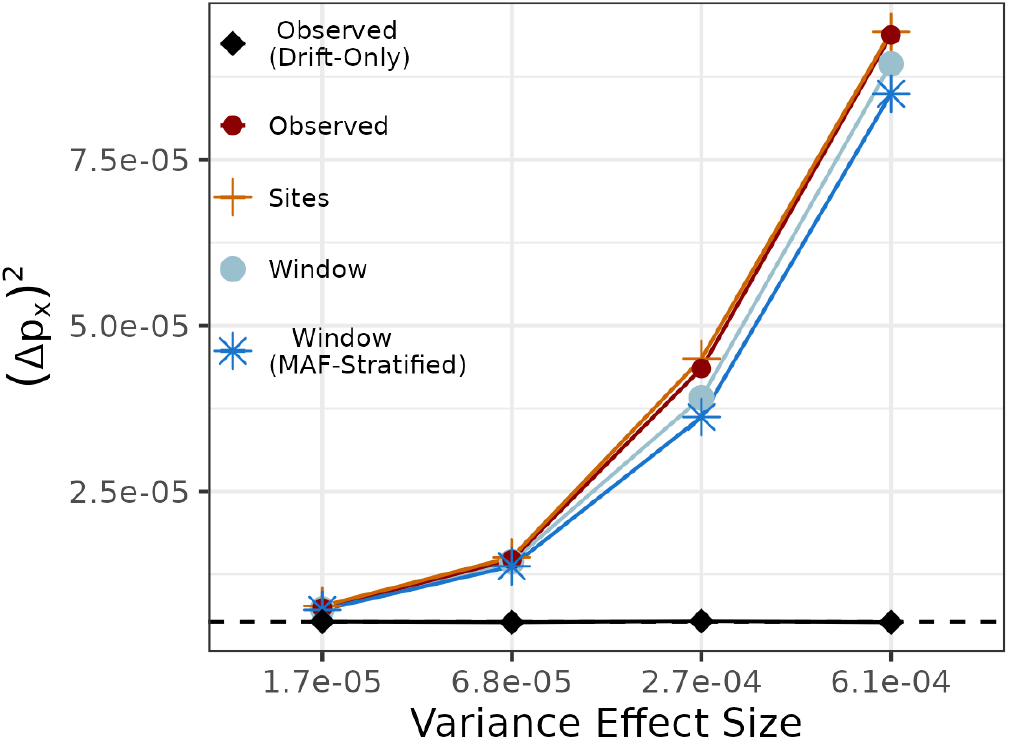
Predicting variance in allele frequency change over a single generation using the “Window”, “Window (MAF-Stratified)” LD Score (neutral + selected sites) and “Sites” LD Score (selected sites) based on Eq. 6 and Eq. S28. The dashed horizontal line denotes the mathematical expectation for a scenario with only drift. The x-axis distinguishes simulations by the variance of the distribution from which allele effect sizes were sampled. The y-axis shows the average squared allele frequency change. Simulations were carried out with *N* = 10, 000 diploid individuals, 5 Mbp chromosomes, a rescaled recombination map of human chromosome 22, and 25% of variant sites under directional selection.

**Figure S2:**
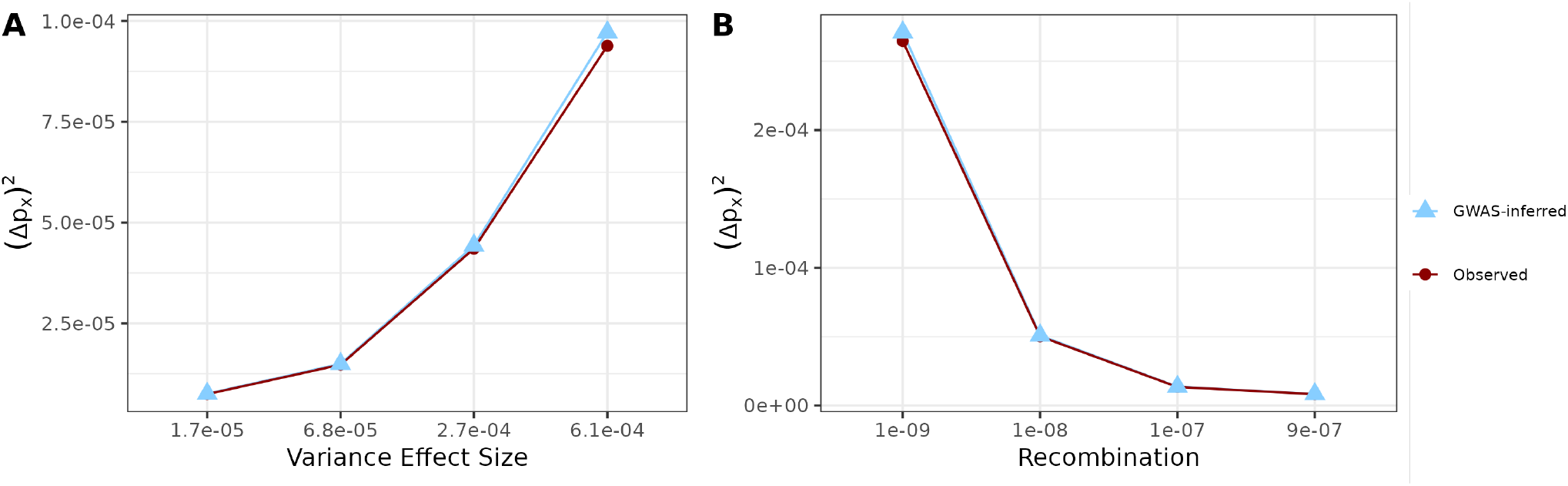
Population GWAS effect sizes on reproductive success predicts average squared allele frequency change over a single generation. **A)**. Simulations as described in Fig. S1. **B)** Simulations varying in constant recombination rate (x-axis) but with the same variance in effect size (*V*_*g*_ = 2.7 *×* 10^−4^). The prediction is based on Eq. S60.

### S1 Number of live births - Female UK Biobank phenotype

#### S1.1 Non-partitioned LD-score

**Figure S3:**
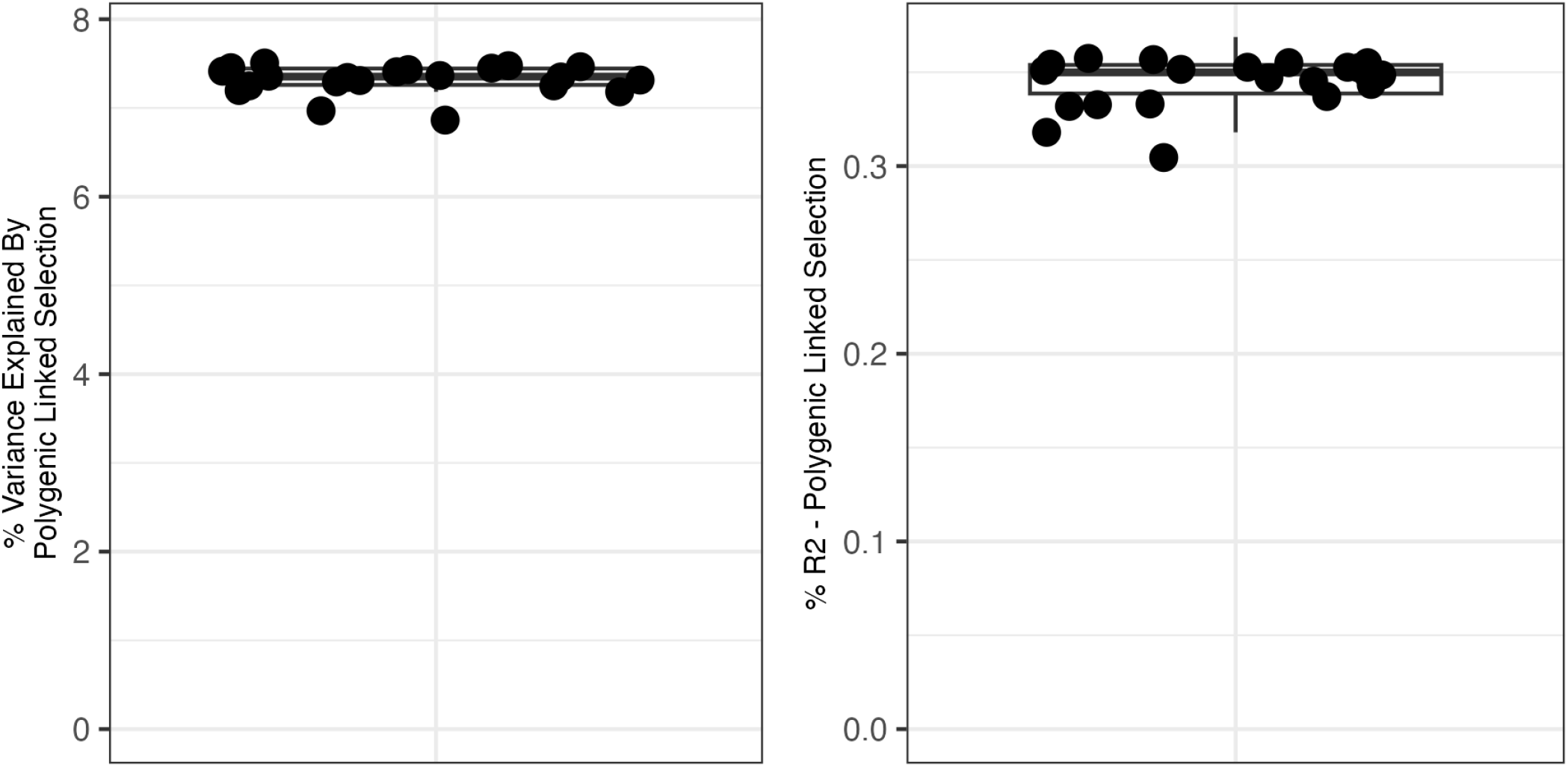
The contribution of polygenic linked selection on genome-wide squared allele frequency change estimated from population GWAS effect sizes in females on the phenotype ‘number of live births’ without PCs covariates. Left and right panels show the percentage of squared allele frequency change and R-squared explained by polygenic linked selection. Each dot represents a leave-one-chromosome-out replicate of the estimate. Polygenic linked selection is estimated by considering both LD-score and B-value and following Eq. 10,13-16.

**Figure S4:**
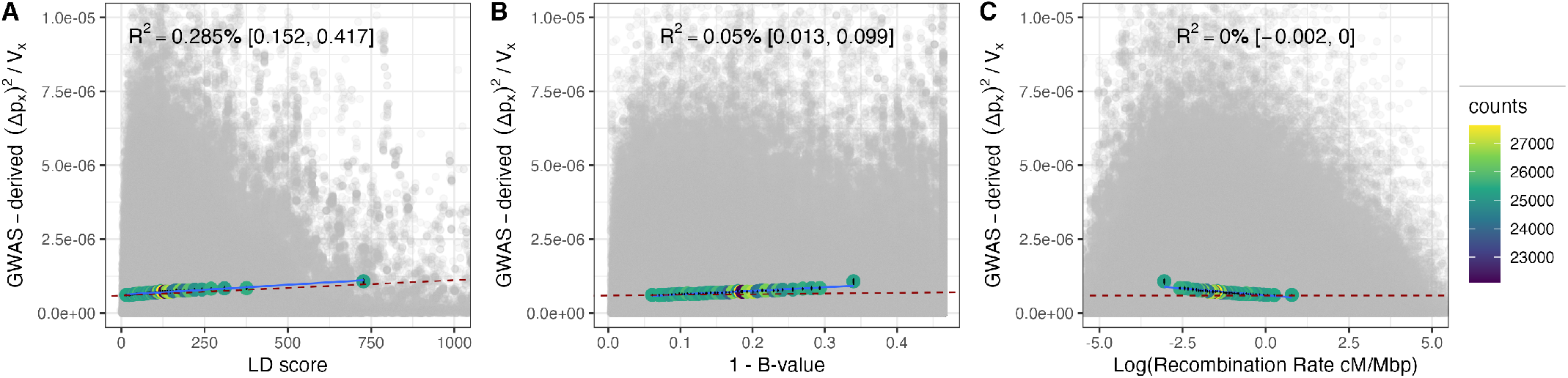
Decomposing the squared allele frequency change inferred from population GWAS effect sizes for common SNPs. Here the GWAS is in females on the phenotype ‘number of live births’ with PCA covariates. Each grey dot is a single SNP’s estimated squared allele frequency change, normalized by *V*_*x*_(1 + *F*_*x*_). In each panel the same set of SNPs are plotted by the LD score, B-value, log_10_ recombination rate. The colored points show the average calculated in 2.5% quantiles of the LD score predictor. The dashed red line shows the slope obtained from the multivariate regression, the *R*^2^ and its 95% confidence interval based on the leave-one-chromosome-out procedure is reported as a percentage at the top of each panel. The blue line shows the linear regression on the average colored points.

**Figure S5:**
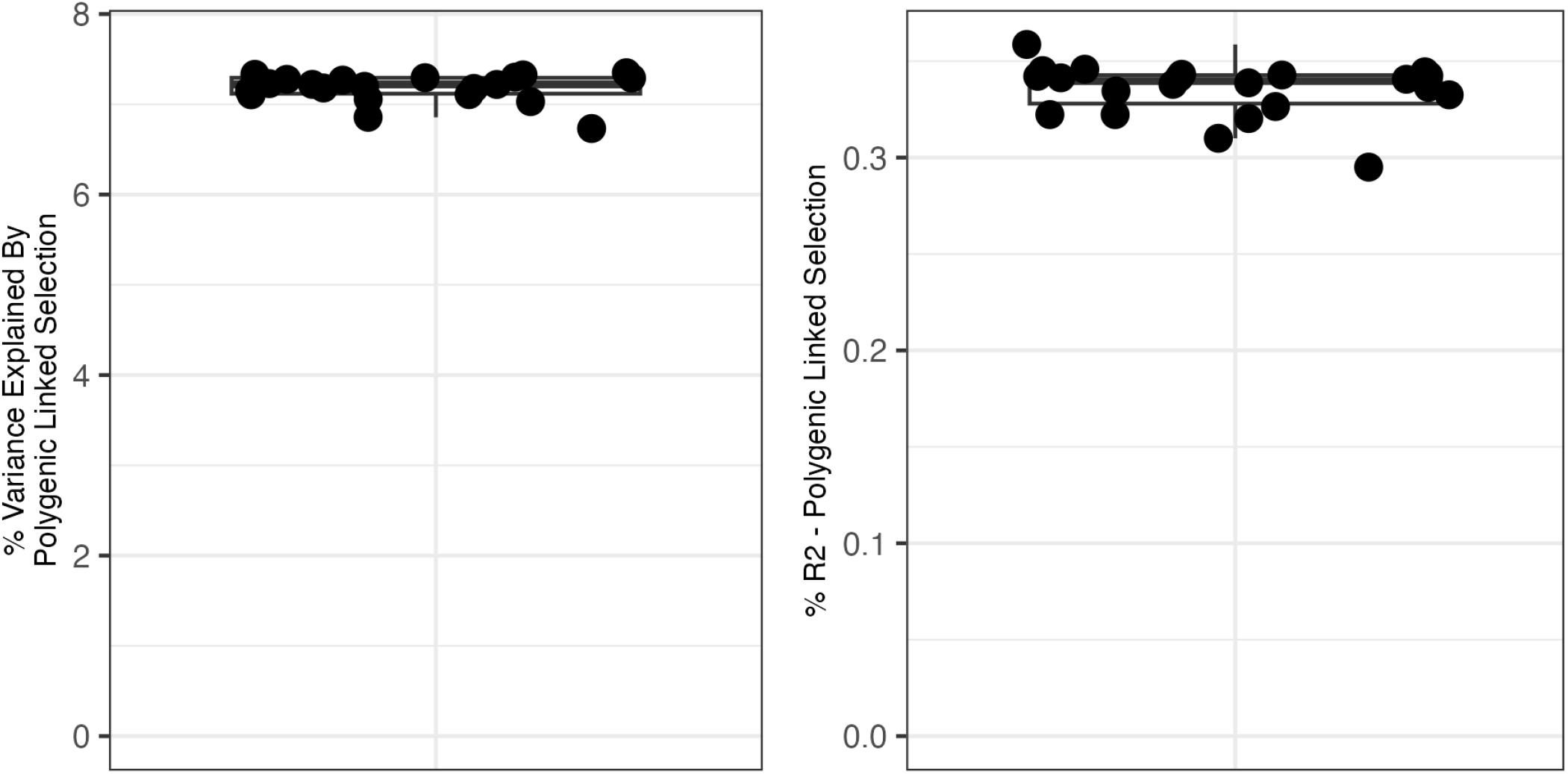
The contribution of polygenic linked selection on genome-wide squared allele frequency change estimated from population GWAS effect sizes in females on the phenotype ‘number of live births’ with PCs covariates. See caption in Fig. S3 for more details.

**Figure S6:**
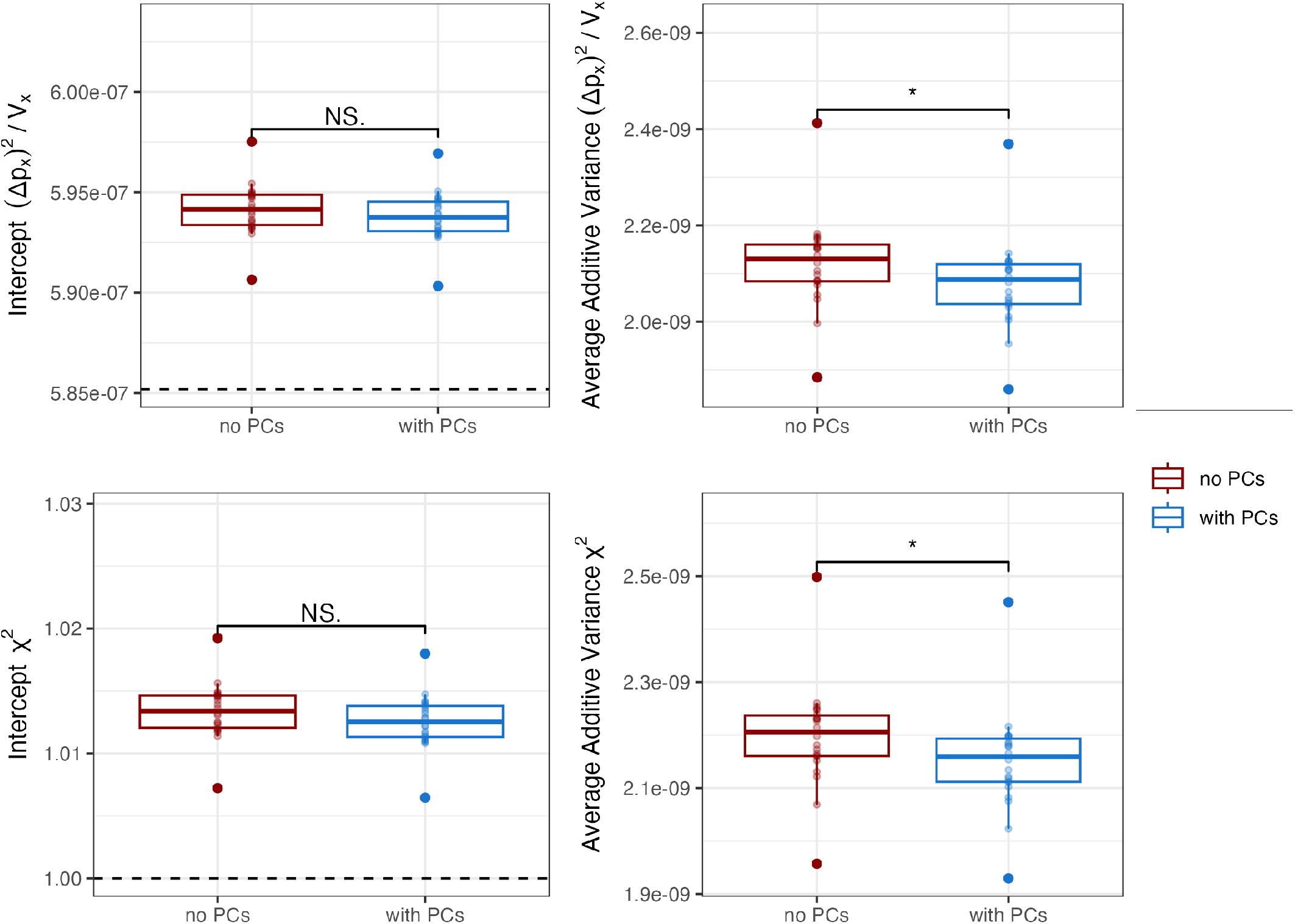
Multiregression results on genome-wide squared allele frequency change estimated from population GWAS effect size in females on the phenotype ‘number of live births’ with and without PCs as covariates. First and second row shows the intercept and slope from the multiregression on the squared allele frequency change and population GWAS *χ*^2^ statistics, respectively. The multiregression was carried out including as predictors LD score, B-value and log_*e*_ recombination rate. Each dot represents a leave-one-chromosome-out replicate of the estimate.

**Figure S7:**
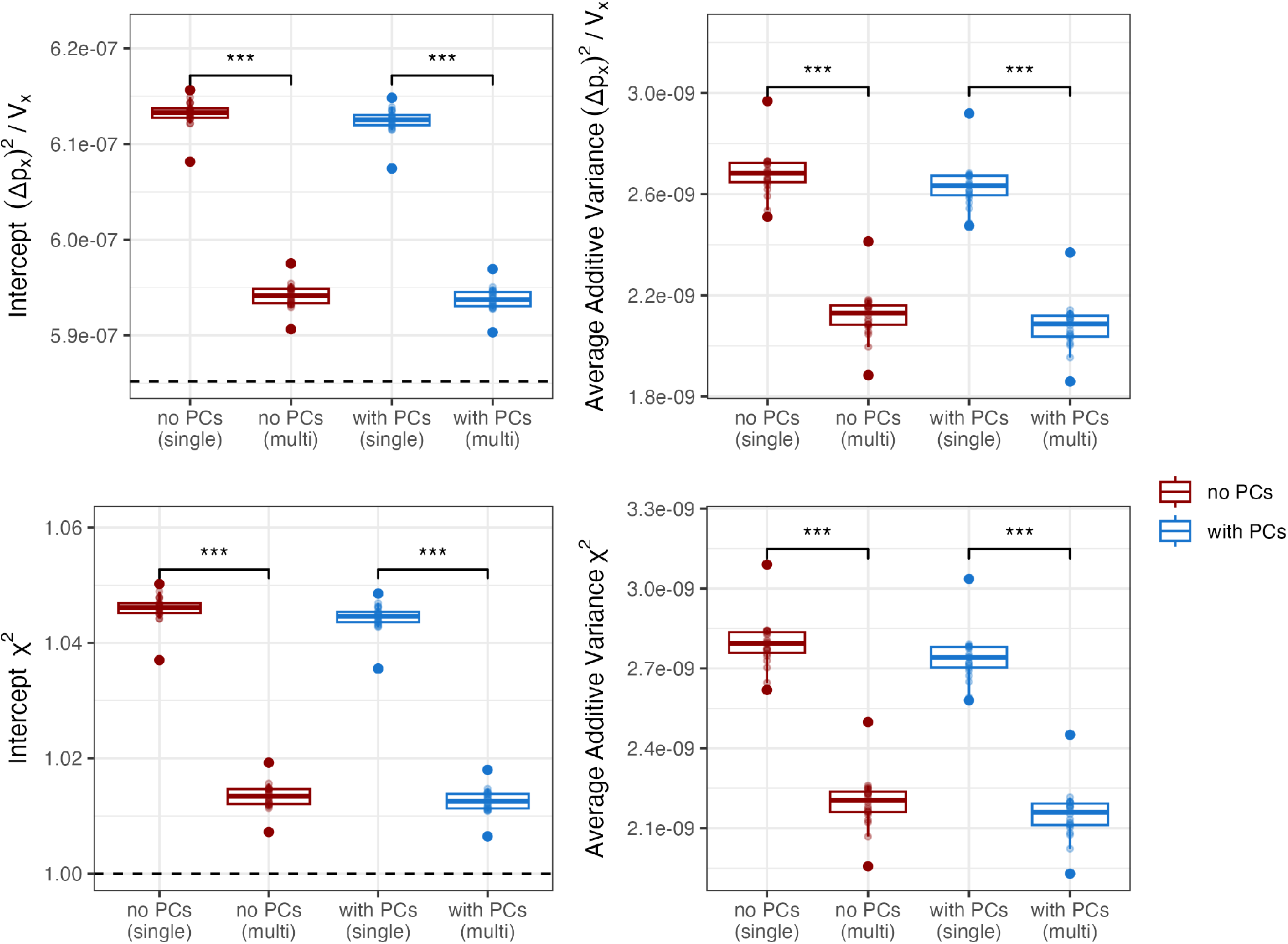
Multivariate and univariate regression results on genome-wide squared allele frequency change estimated from population GWAS effect size in females on the phenotype ‘number of live births’ with and without PCs as covariates. Univariate regression was performed by using LD score as the only predictor. See caption in Fig. S6 for more details.

#### S1.2 Partitioned LD-score

**Figure S8:**
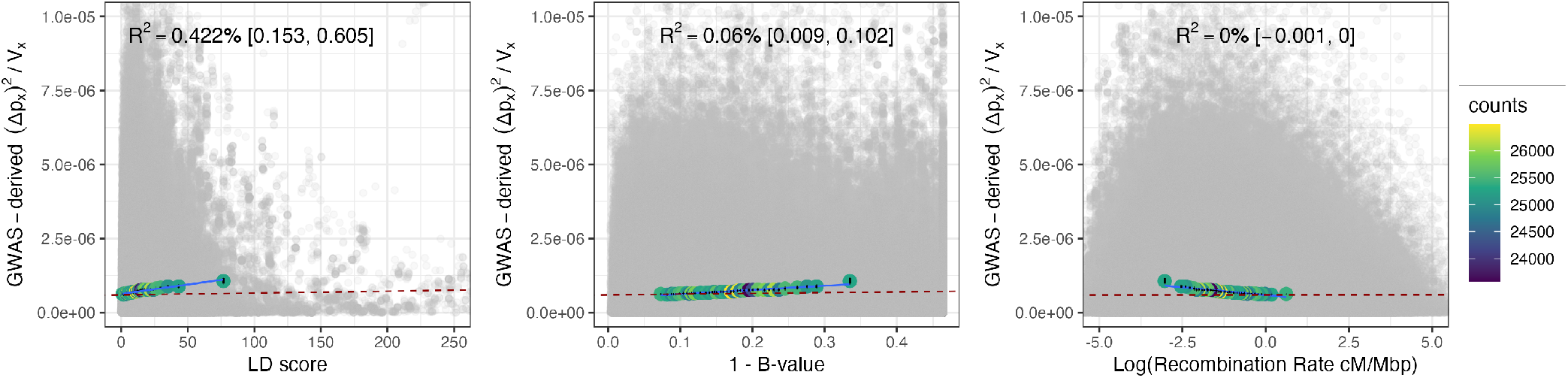
Decomposing the squared allele frequency change inferred from population GWAS effect sizes, estimated in females for the phenotype ‘number of live births’ without PC covariates, using MAF-stratified LD score. See caption in Fig. S4 for more details.

**Figure S9:**
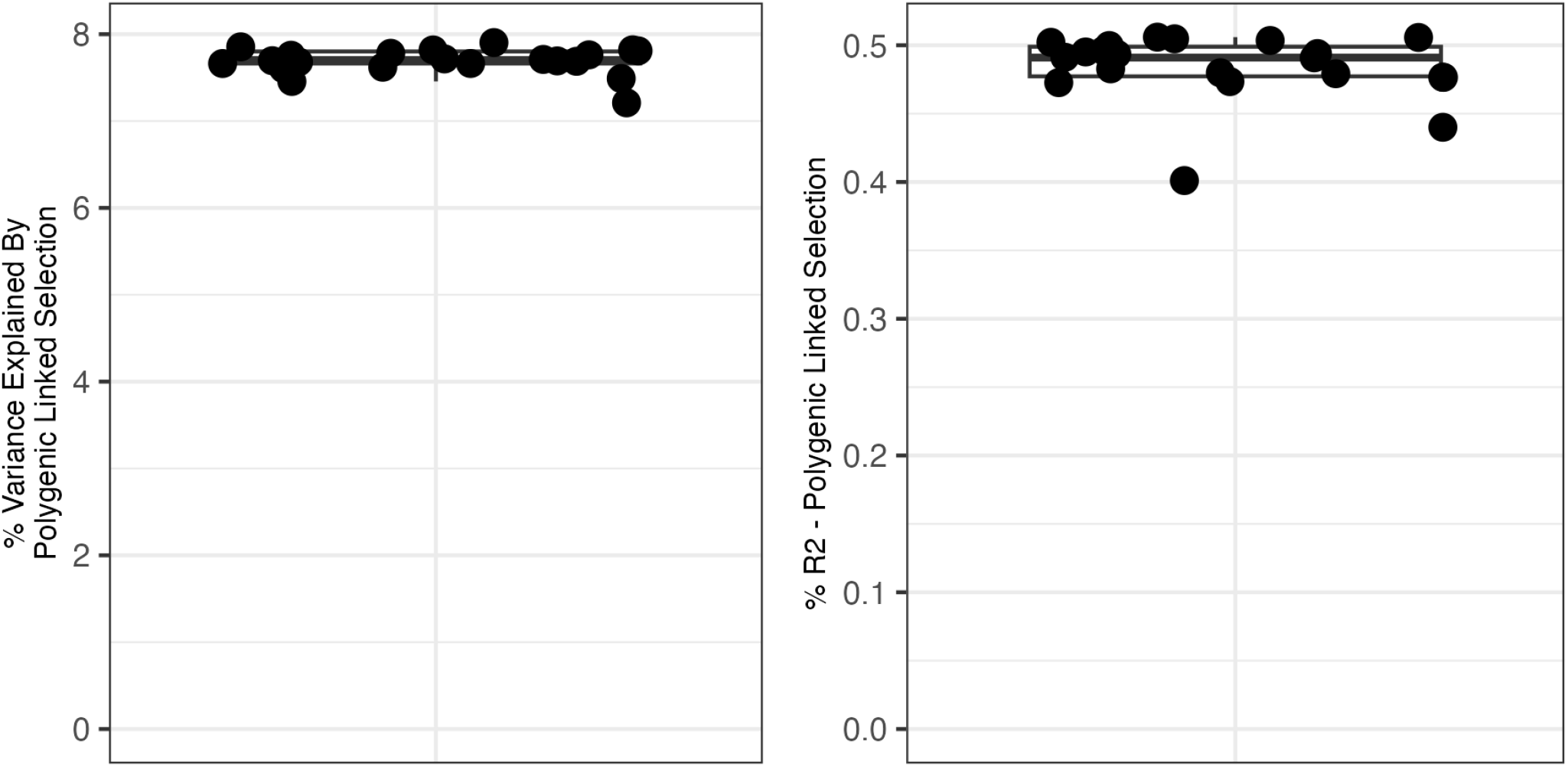
The contribution of polygenic linked selection on genome-wide squared allele frequency change inferred from population GWAS effect sizes, estimated in females for the phenotype ‘number of live births’ without PCs covariates. Here, estimates are based on a MAF-stratified LD score regression, together with B-value and log_*e*_ recombination rate. See caption in Fig. S3 for more details.

**Figure S10:**
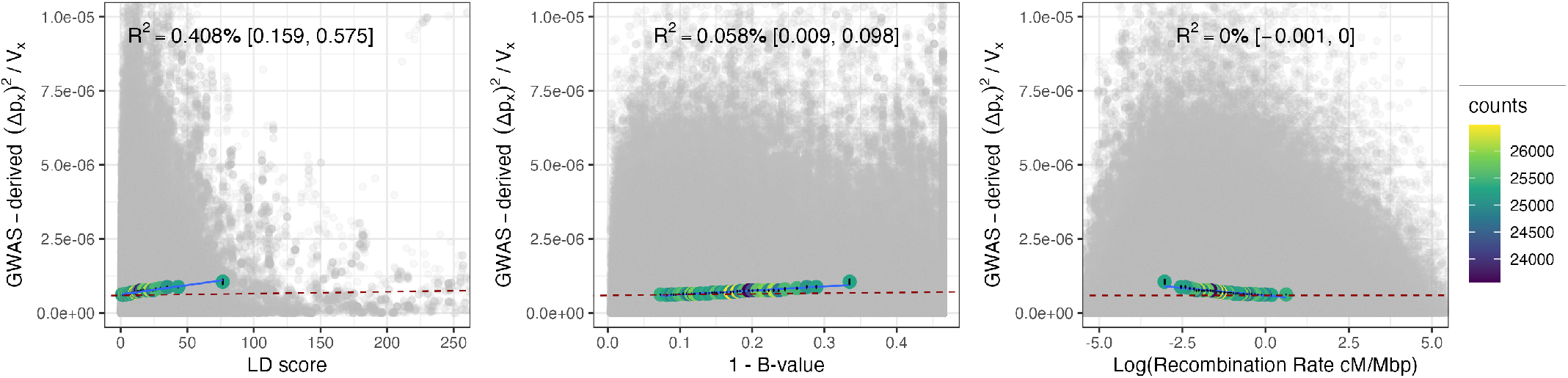
Decomposing the squared allele frequency change inferred from population GWAS effect sizes, estimated in females for the phenotype ‘number of live births’ with PC covariates, using MAF-stratified LD score. See caption in Fig. S4 for more details.

**Figure S11:**
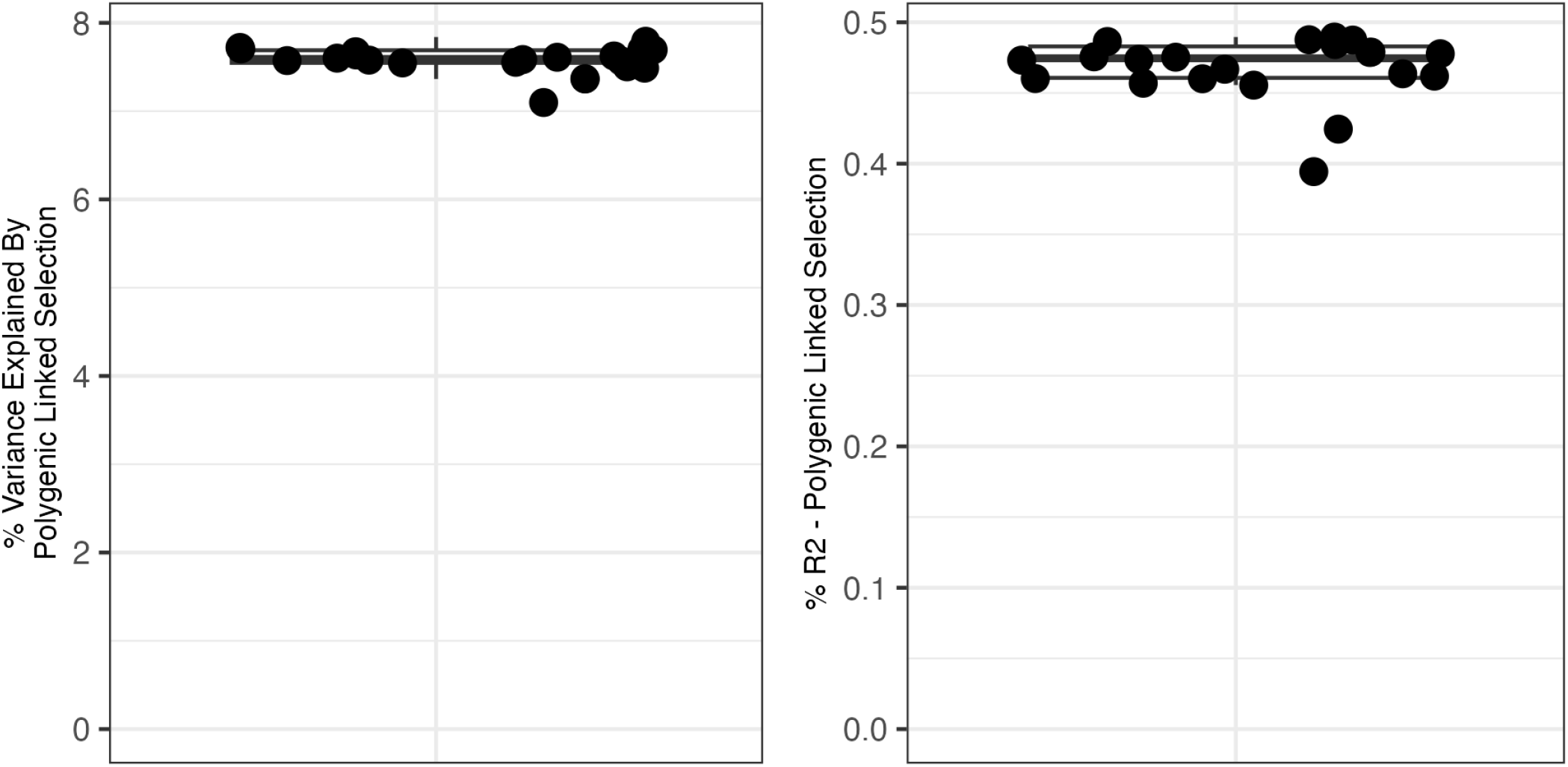
The contribution of polygenic linked selection on genome-wide squared allele frequency change inferred from population GWAS effect sizes, estimated in females for the phenotype ‘number of live births’ with PCs covariates. See caption in Fig. S9 for more details.

### S2 Number of children fathered - Male UK Biobank phenotype

#### S2.1 Non-partitioned LD-score

**Figure S12:**
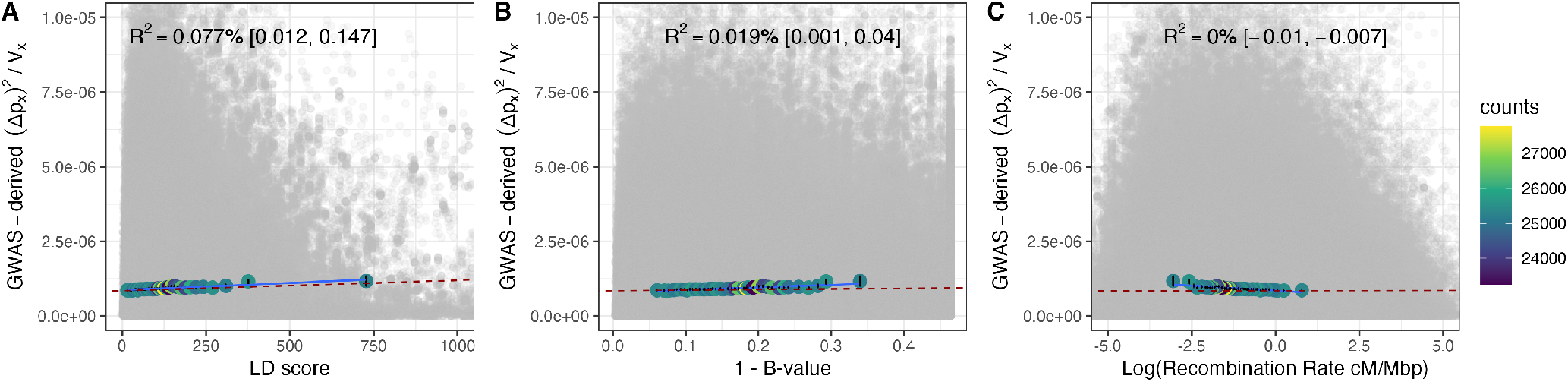
Decomposing the squared allele frequency change inferred from population GWAS effect sizes for common SNPs. Here the GWAS is in males on the phenotype ‘number of children fathered’ without PCs covariates. See caption in Fig. S4 for more details.

**Figure S13:**
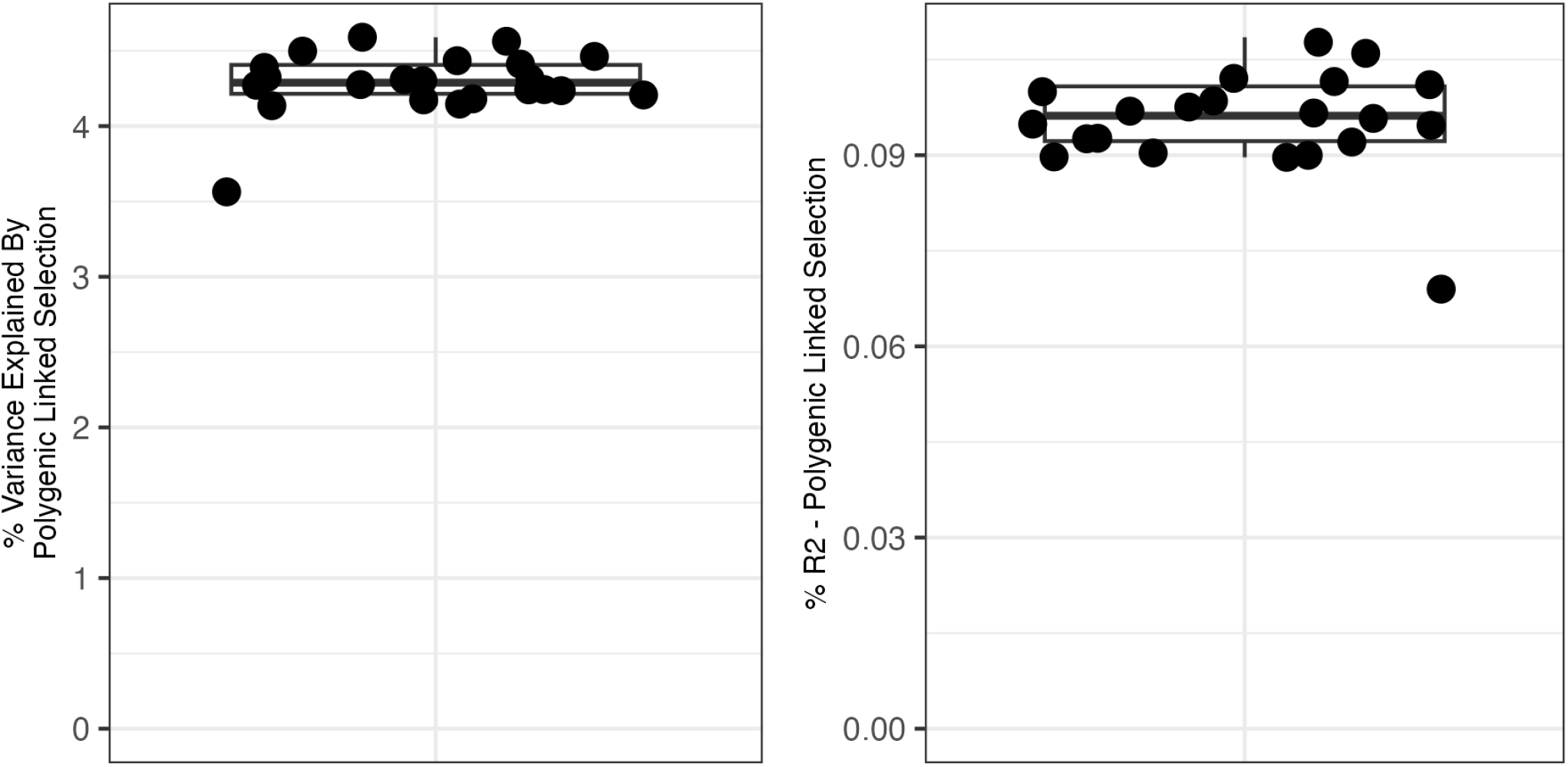
The contribution of polygenic linked selection on genome-wide squared allele frequency change estimated from population GWAS effect sizes in males on the phenotype ‘number of children fathered’ without PCs covariates. See caption in Fig. S3 for more details.

**Figure S14:**
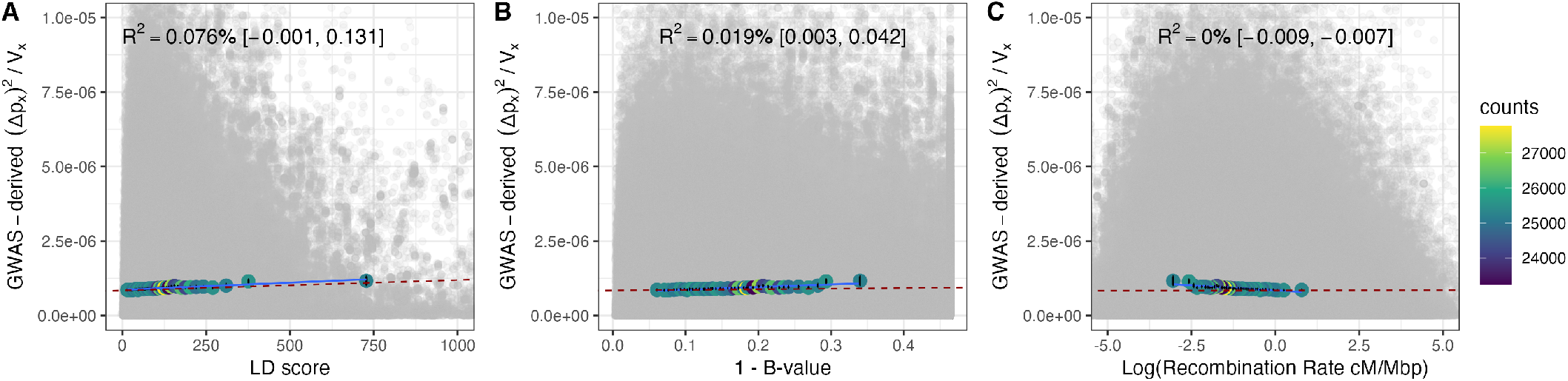
Decomposing the squared allele frequency change inferred from population GWAS effect sizes for common SNPs. Here the GWAS is in males on the phenotype ‘number of children fathered’ with PCs covariates. See caption in Fig. S4 for more details.

**Figure S15:**
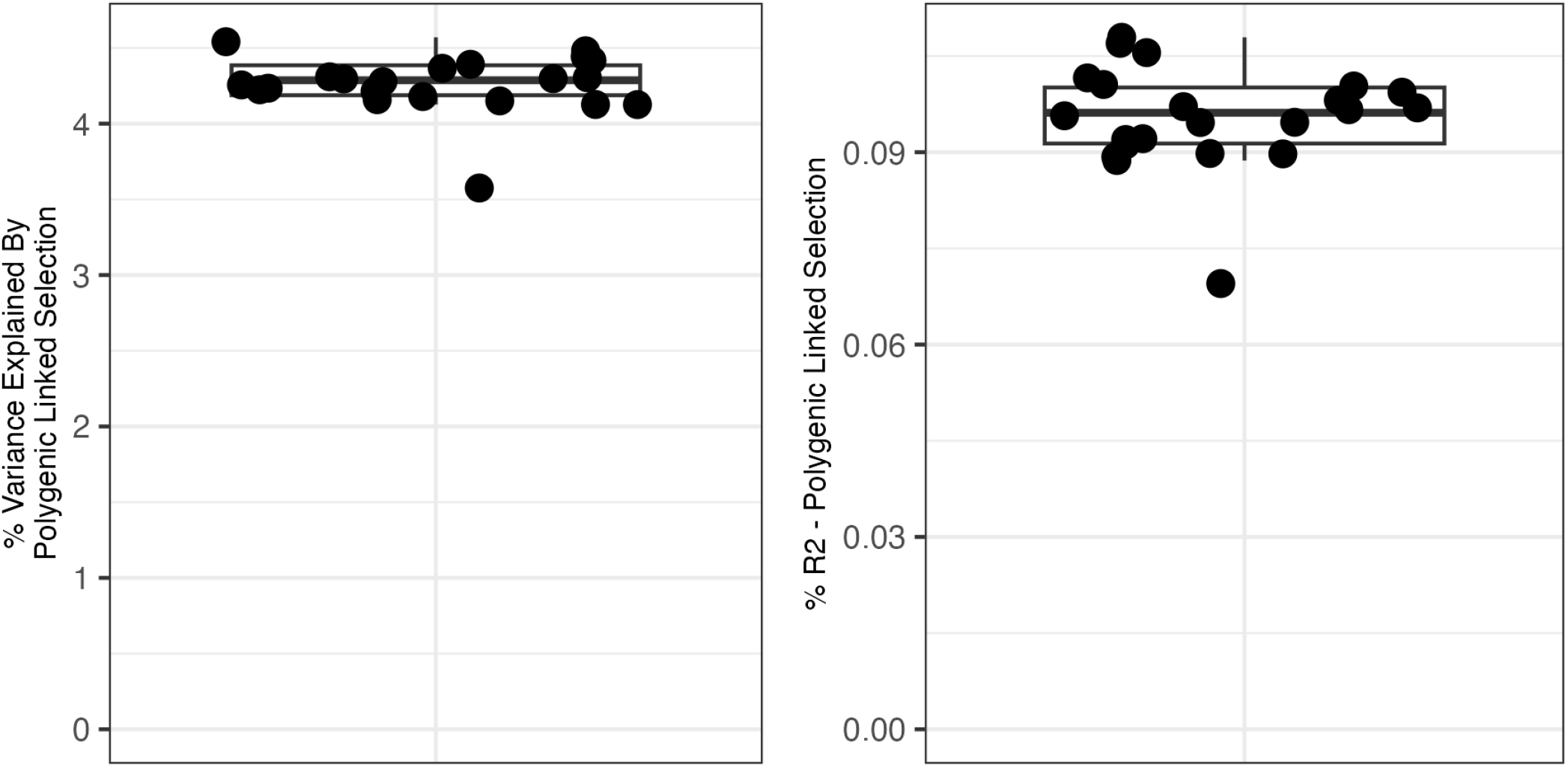
The contribution of polygenic linked selection on genome-wide squared allele frequency change estimated from population GWAS effect sizes in males on the phenotype ‘number of children fathered’ with PCs covariates. See caption in Fig. S3 for more details.

**Figure S16:**
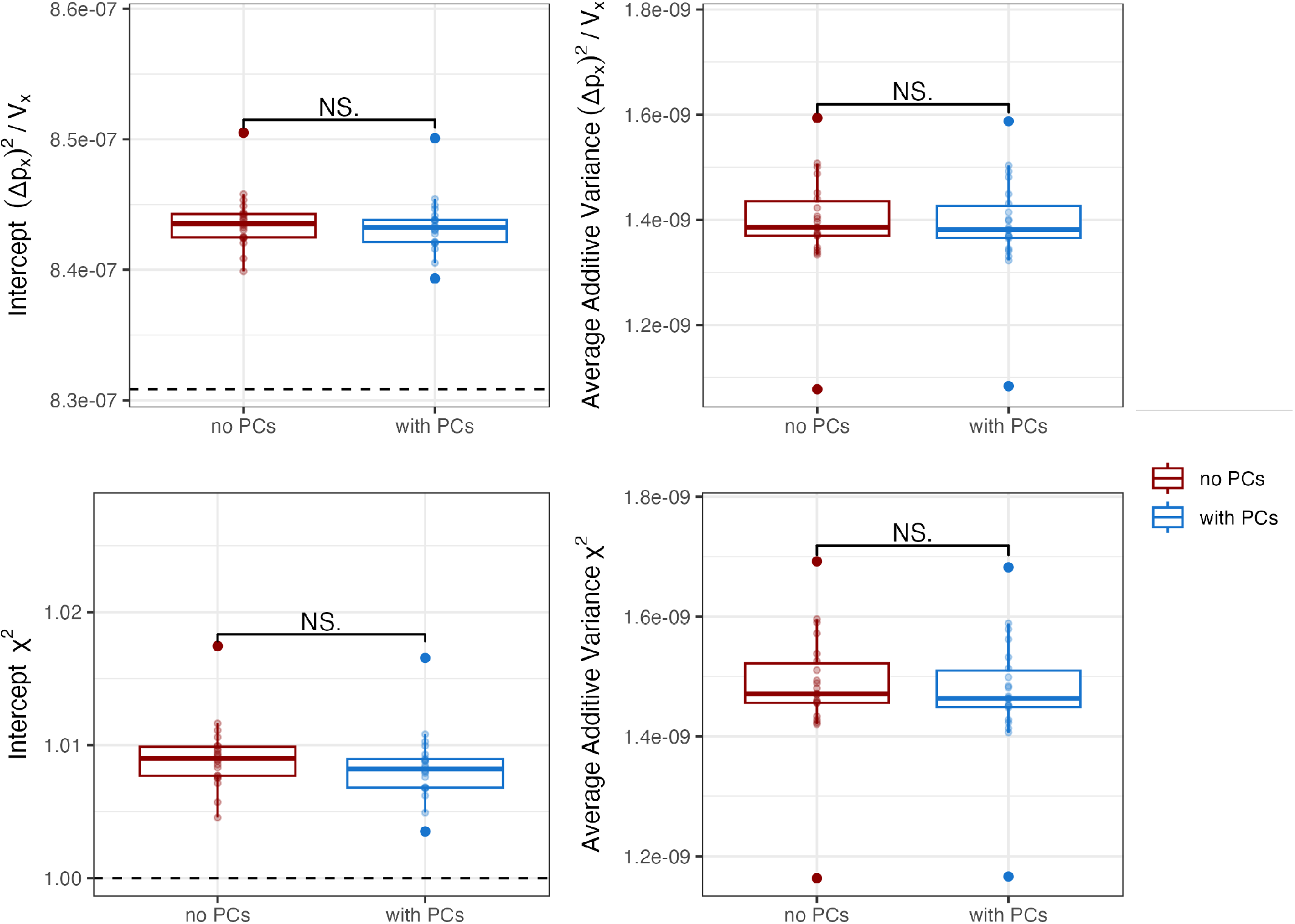
Multiregression results on genome-wide squared allele frequency change estimated from population GWAS effect size in males on the phenotype ‘number of children fathered’ with and without PCs as covariates. See caption in Fig. S6 for more details.

**Figure S17:**
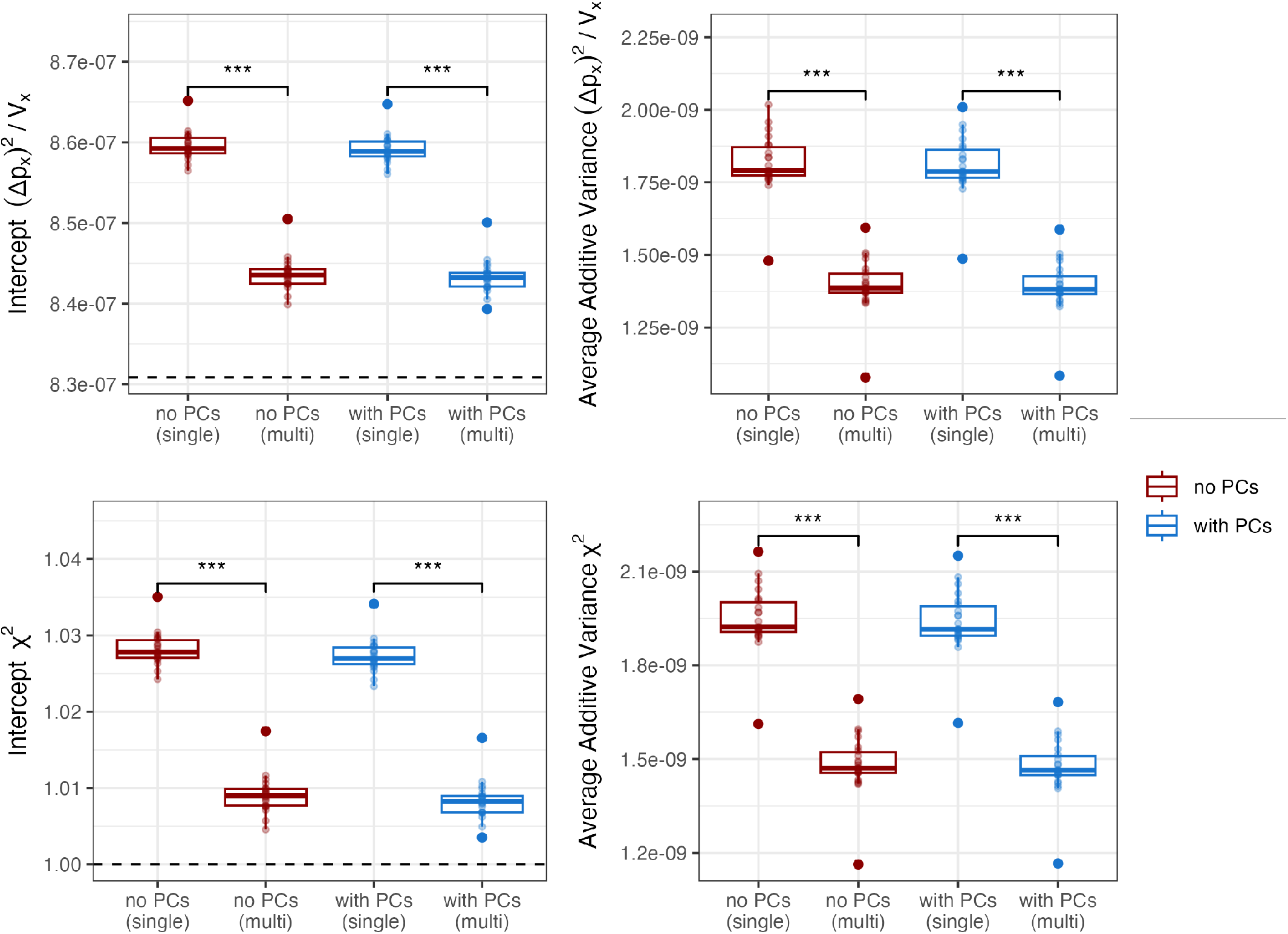
Multivariate and univariate regression results on genome-wide squared allele frequency change estimated from population GWAS effect size in males on the phenotype ‘number of children fathered’ with and without PCs as covariates. Univariate regression was performed by using LD score as the only predictor. See caption in Fig. S7 for more details.

#### S2.2 Partitioned LD-score

**Figure S18:**
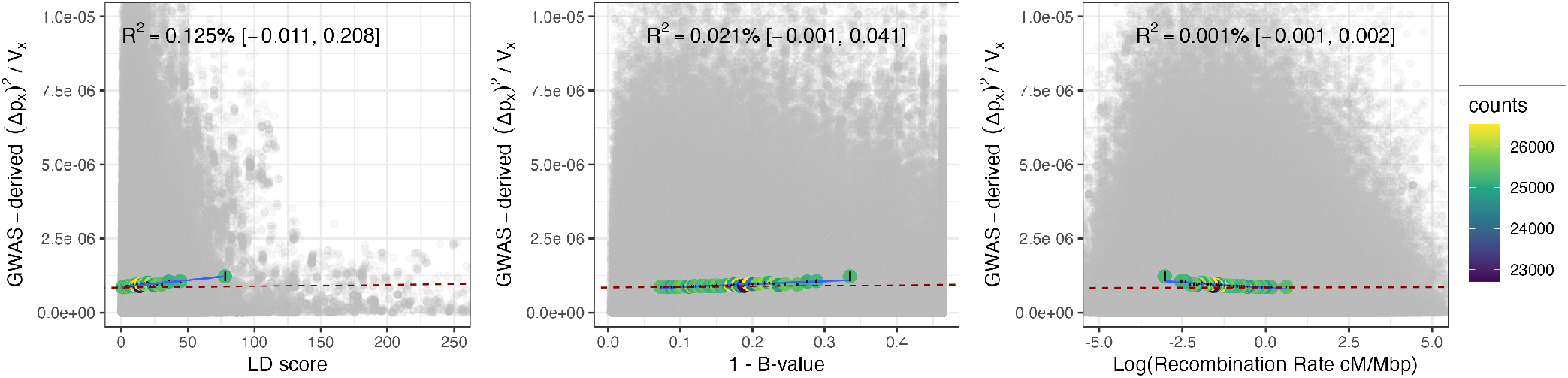
Decomposing the squared allele frequency change inferred from population GWAS effect sizes, estimated in males for the phenotype ‘number of children fathered’ without PC covariates, using MAF-stratified LD score. See caption in Fig. S12.

**Figure S19:**
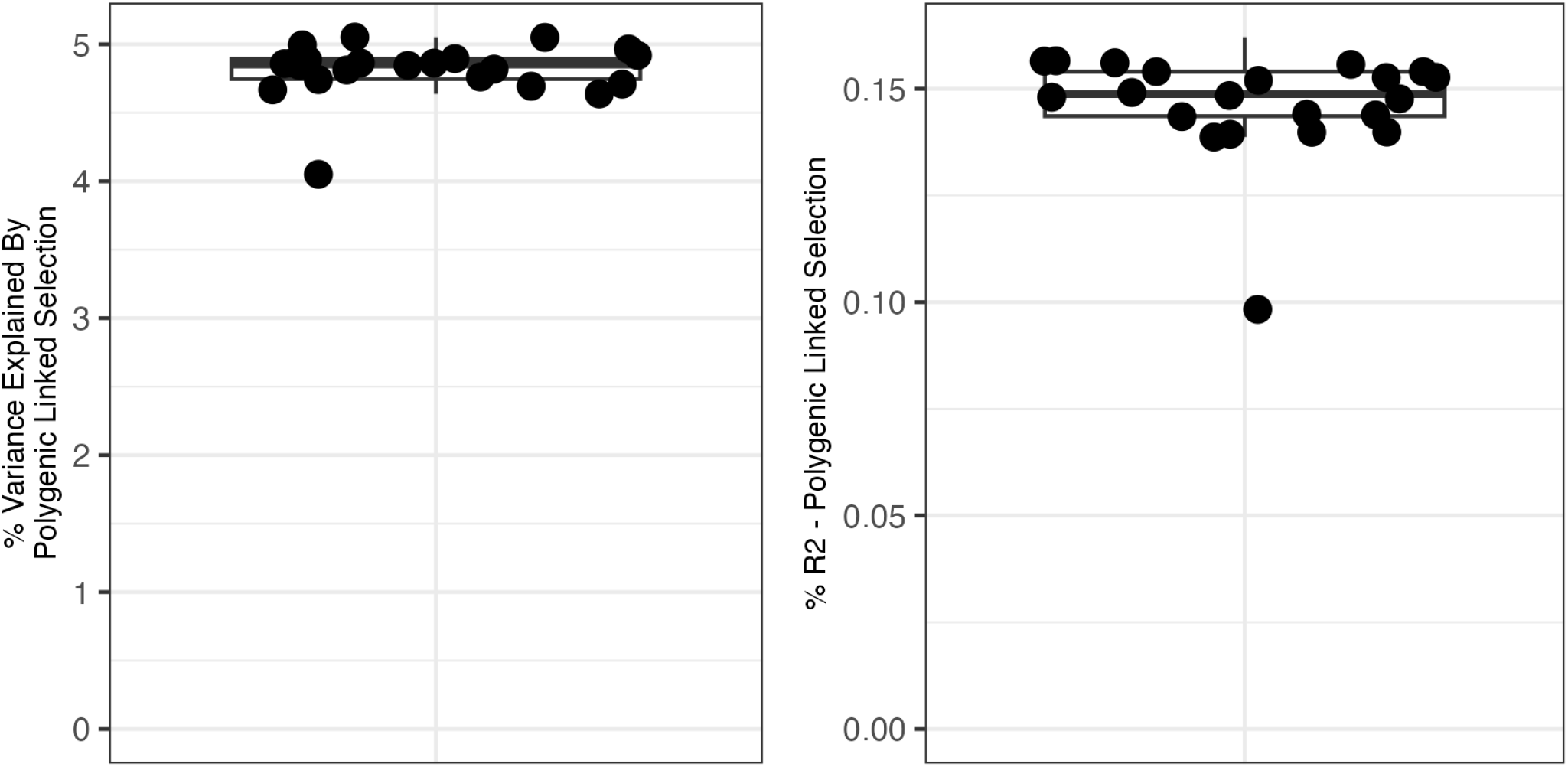
The contribution of polygenic linked selection on genome-wide squared allele frequency change inferred from population GWAS effect sizes, estimated in males for the phenotype ‘number of children fathered’ without PCs covariates. Here, estimates are based on a MAF-stratified LD score regression, together with B-value and log_*e*_ recombination rate. See caption in Fig. S3 for more details.

**Figure S20:**
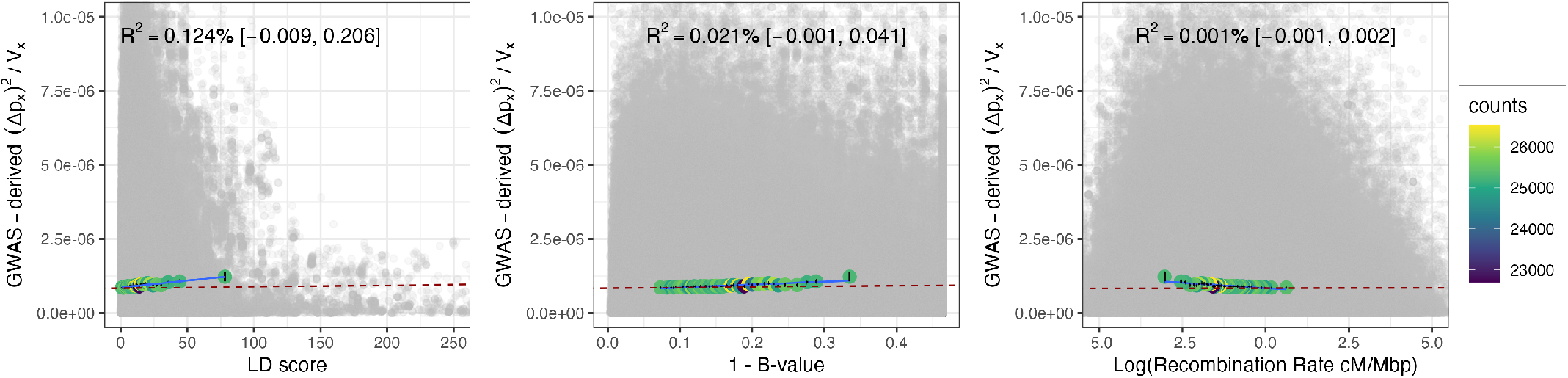
Decomposing the squared allele frequency change inferred from population GWAS effect sizes, estimated in males for the phenotype ‘number of children fathered’ with PC covariates, using MAF-stratified LD score. See caption in Fig. S4 for more details.

**Figure S21:**
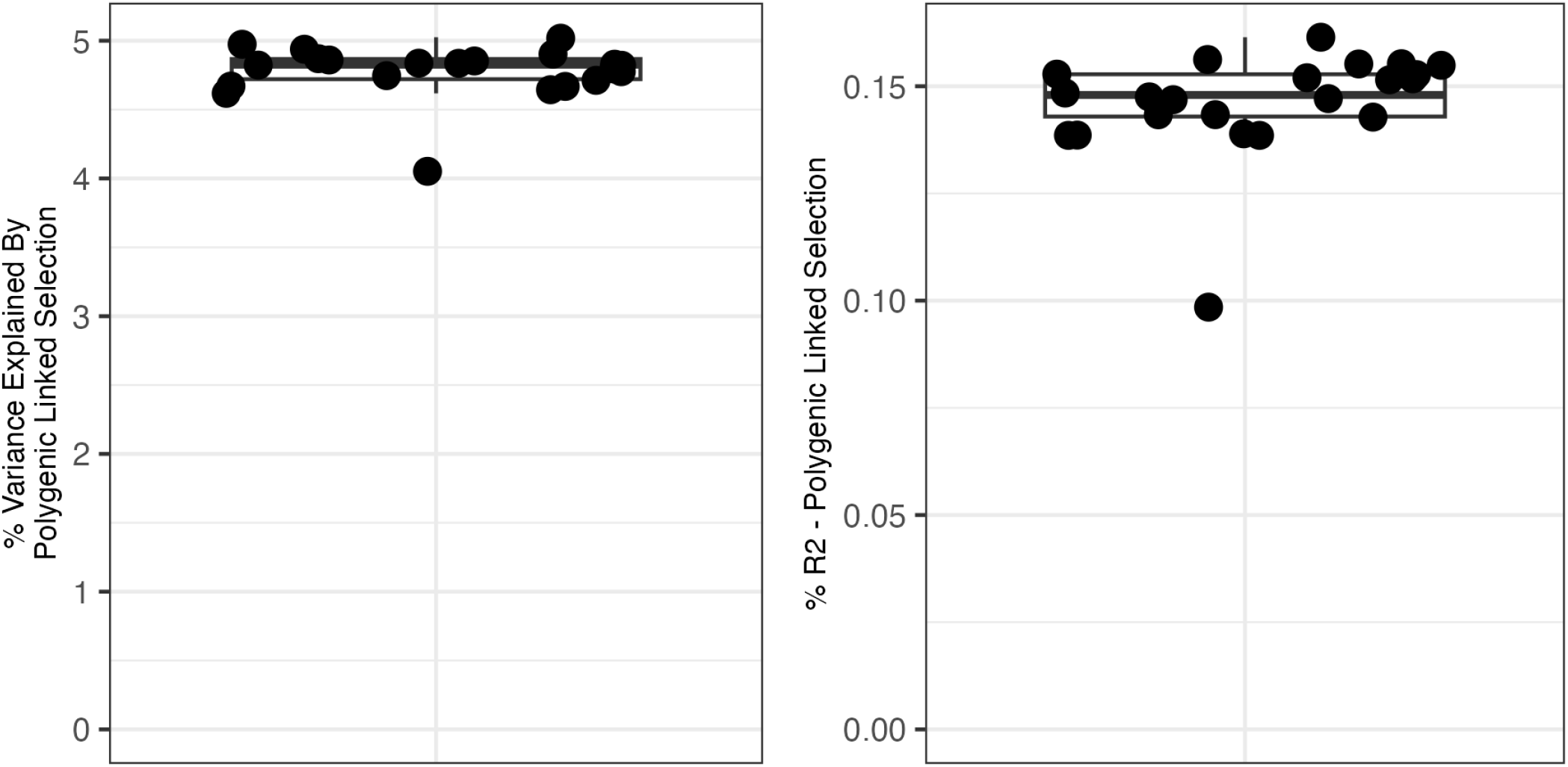
The contribution of polygenic linked selection on genome-wide squared allele frequency change inferred from population GWAS effect sizes, estimated in males for the phenotype ‘number of children fathered’ with PCs covariates. Here, estimates are based on a MAF-stratified LD score regression, together with B-value and log_*e*_ recombination rate. See caption in Fig. S3 for more details.

### S3 Combined Male and Female UK Biobank cohorts

#### S3.1 Weighted-Average Approach

**Figure S22:**
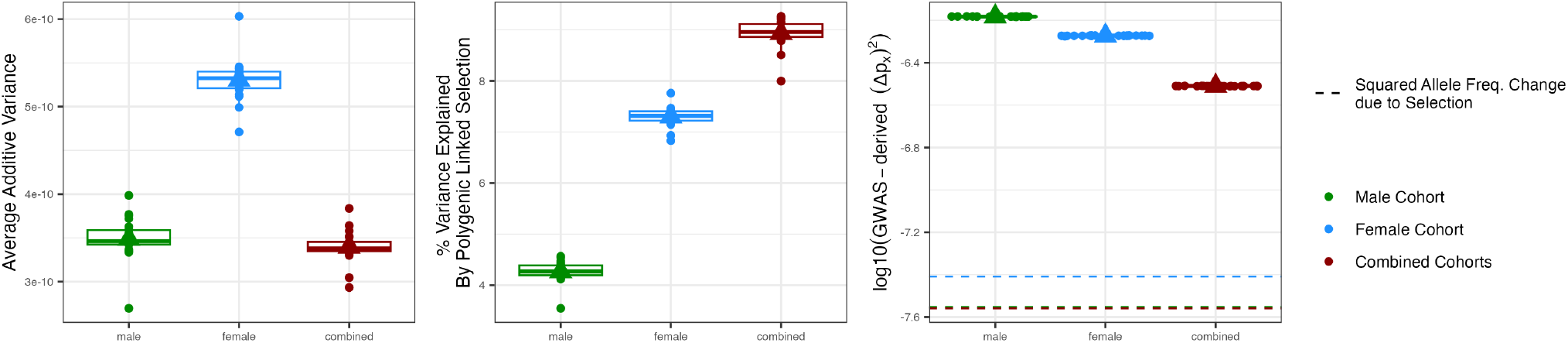
The contribution of polygenic linked selection on genome-wide squared allele frequency change inferred from population GWAS effect sizes estimated in males for the phenotype ‘number of children fathered’, in females for the phenotype ‘number of live births’, and from the combined effect size obtained by weighted averaging. Population GWAS were performed without PCs covariates. The left panel indicates the additive genic variance estimated in each cohort. The right panel shows the estimates of average total genome-wide squared allele frequency change (selection + drift). Here, estimates are based on multivariate regression including LD score, B-value and log_*e*_ recombination rate as predictors. See caption in Fig. S3 for more details.

**Figure S23:**
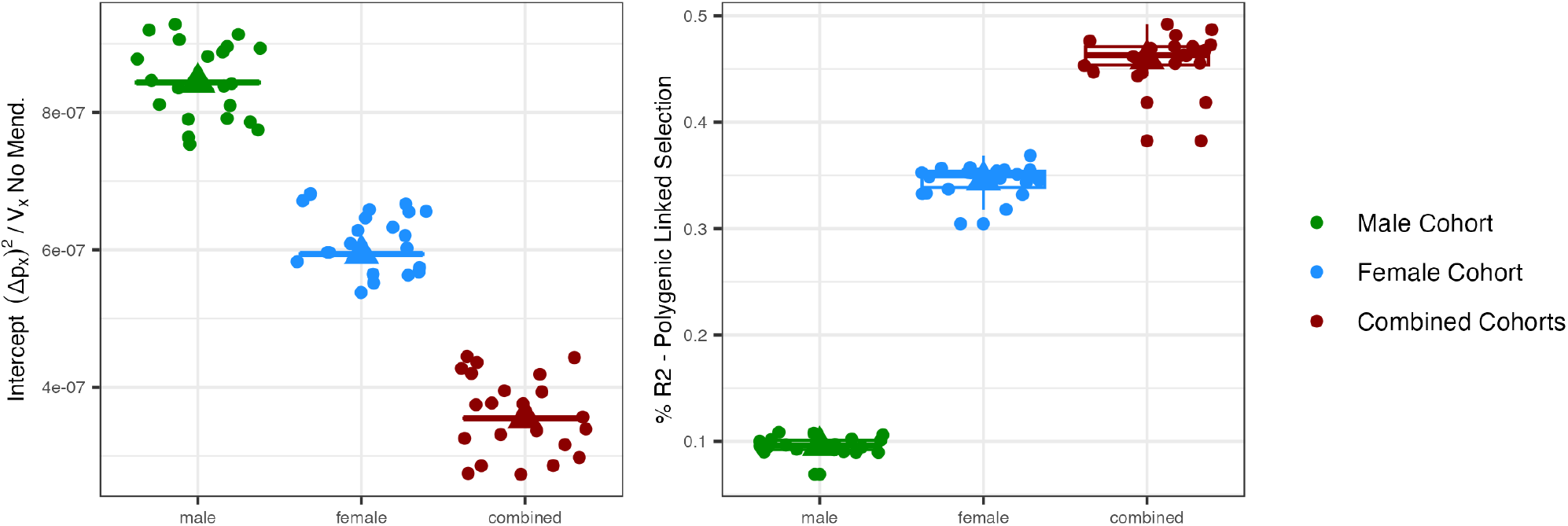
Intercept of the multivariate regressions in Fig. S22 and the corresponding R-squared explained by polygenic linked selection. The intercept absorbs the environmental contribution of the average squared allele frequency change (excluding Mendelian noise).

#### S3.2 Fixed-Effect Approach

**Figure S24:**
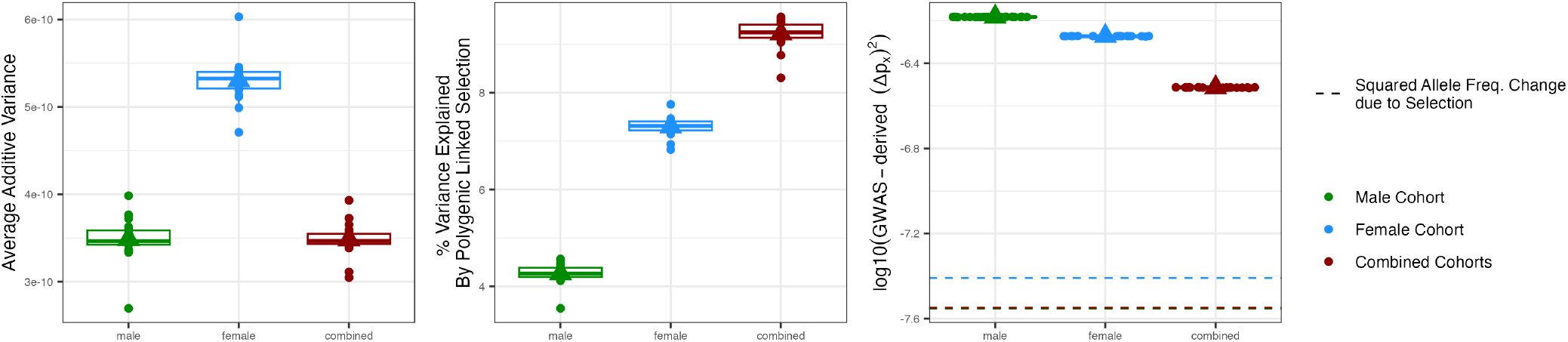
The contribution of polygenic linked selection on genome-wide squared allele frequency change inferred from population GWAS effect sizes estimated in males for the phenotype ‘number of children fathered’, in females for the phenotype ‘number of live births’, and from the combined effect size obtained by the fixed-effect approach. Population GWAS were performed without PCs covariates. The left panel indicates the additive genic variance estimated in each cohort. The right panel shows the estimates of average total genome-wide squared allele frequency change (selection + drift). Here, estimates are based on multivariate regression including LD score, B-value and log_*e*_ recombination rate as predictors. See caption in Fig. S3 for more details.

**Figure S25:**
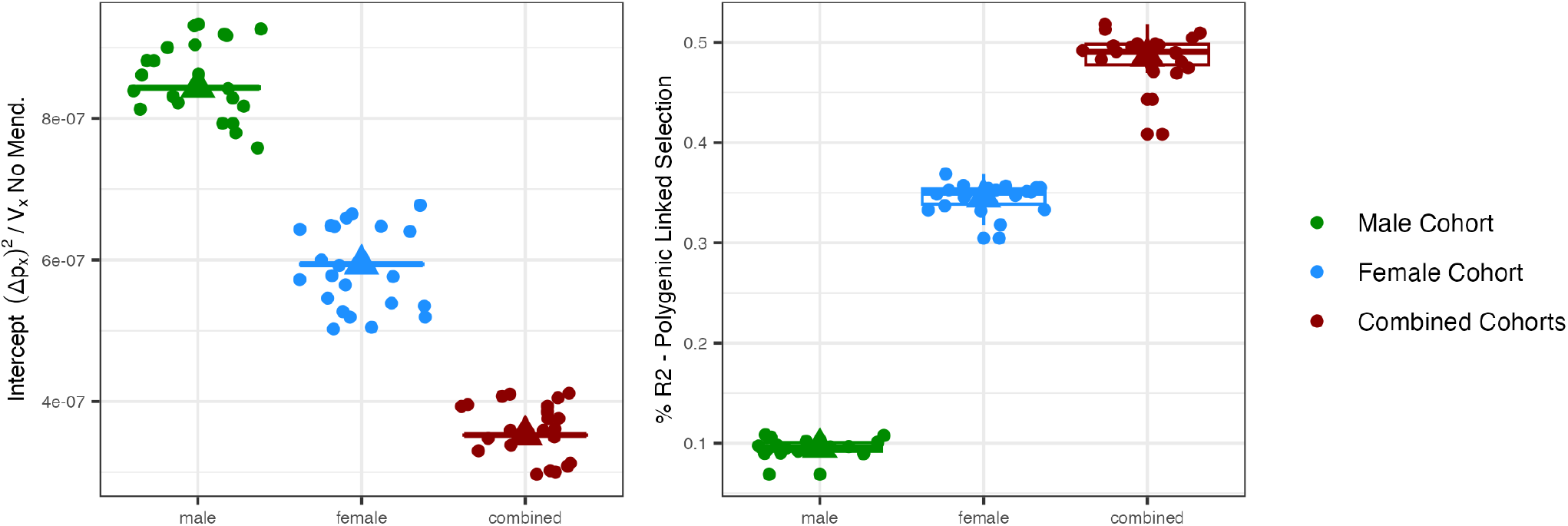
Intercept of the multivariate regressions in Fig. S24 and the corresponding R-squared explained by polygenic linked selection. The intercept absorbs the environmental contribution of the average squared allele frequency change (excluding Mendelian noise).

#### S3.3 Random-Effect Approach

**Figure S26:**
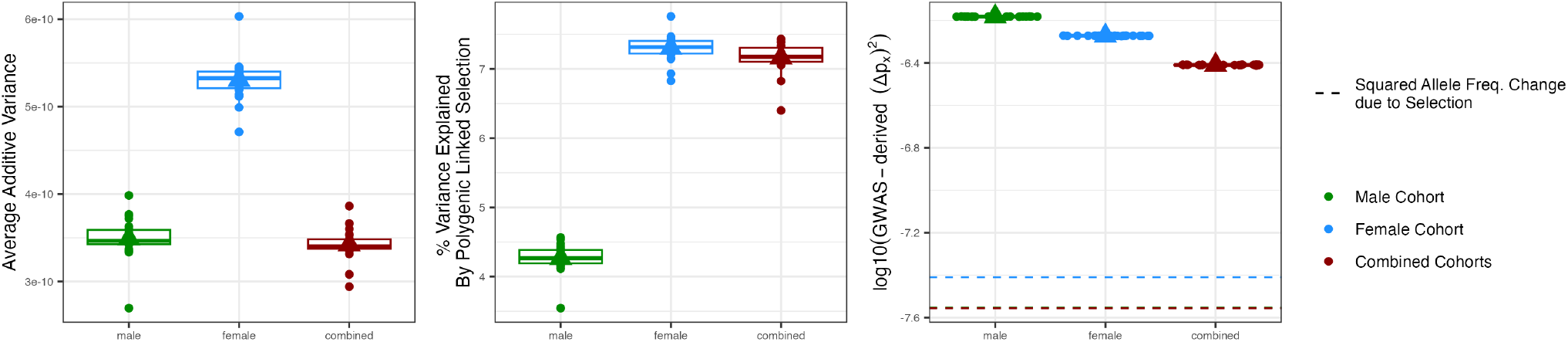
The contribution of polygenic linked selection on genome-wide squared allele frequency change inferred from population GWAS effect sizes estimated in males for the phenotype ‘number of children fathered’, in females for the phenotype ‘number of live births’, and from the combined effect size obtained by the random-effect approach. Population GWAS were performed without PCs covariates. The left panel indicates the additive genic variance estimated in each cohort. The right panel shows the estimates of average total genome-wide squared allele frequency change (selection + drift). Here, estimates are based on multivariate regression including LD score, B-value and log_*e*_ recombination rate as predictors. See caption in Fig. S3 for more details.

**Figure S27:**
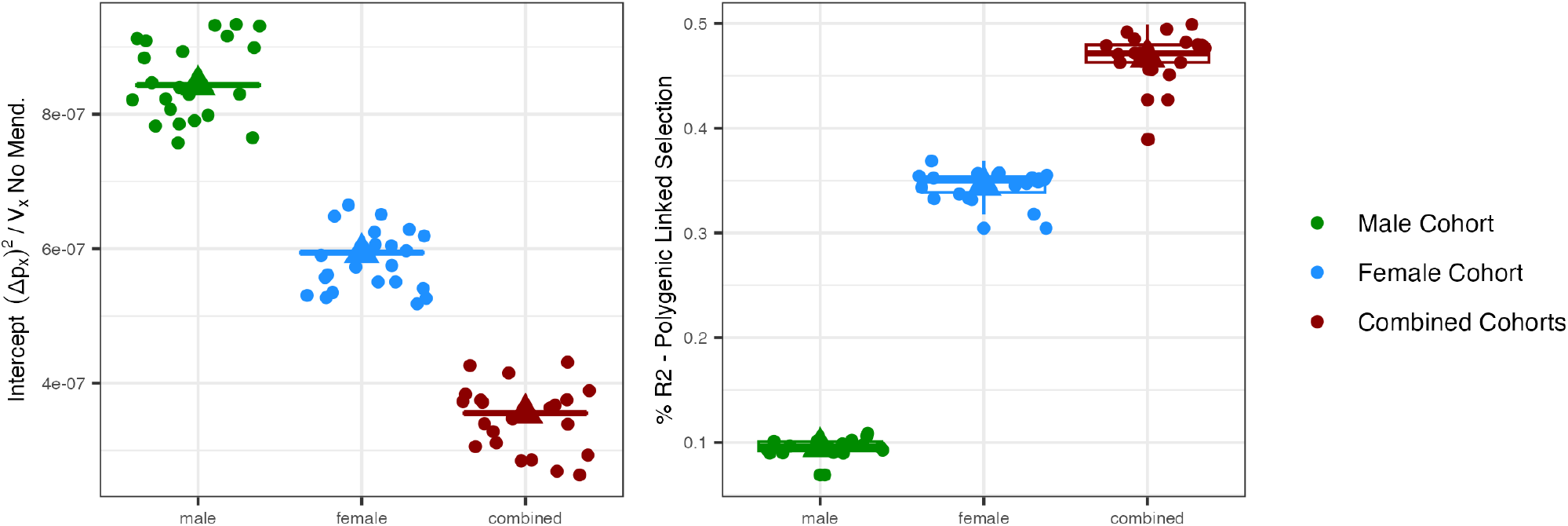
Intercept and the R-squared explained by polygenic linked selection from the multivariate regressions in Fig. S26. The intercept absorbs the environmental contribution of the average squared allele frequency change (excluding Mendelian noise).

### S4 LD score regression on sibling-GWAS effect sizes

**Figure S28:**
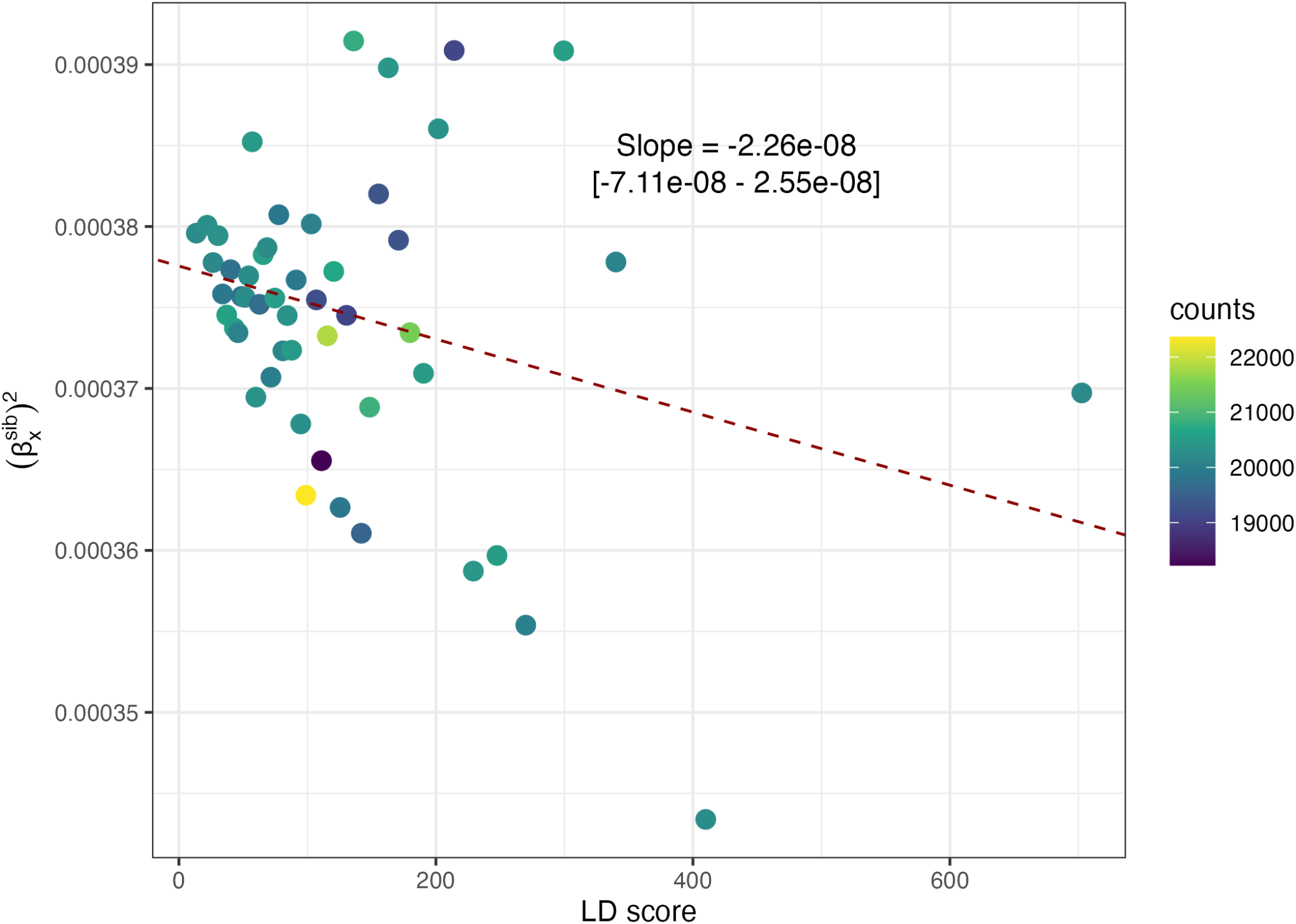
LD score regression using sibling-based GWAS for the female phenotype ‘number of live births’. Points show average values in bins of 2% quantiles of LD score. The linear regression through the underlying data is shown, and summary statistics are given as text. The SNP effect sizes from sib-GWAS has slight adjustment for inbreeding (*F*_*x*_) to place estimated effects on the same scale as for the population-based GWAS (see Eqs. S68, S77 and S84).

### S5 Ancient and Contemporary Selection

**Figure S29:**
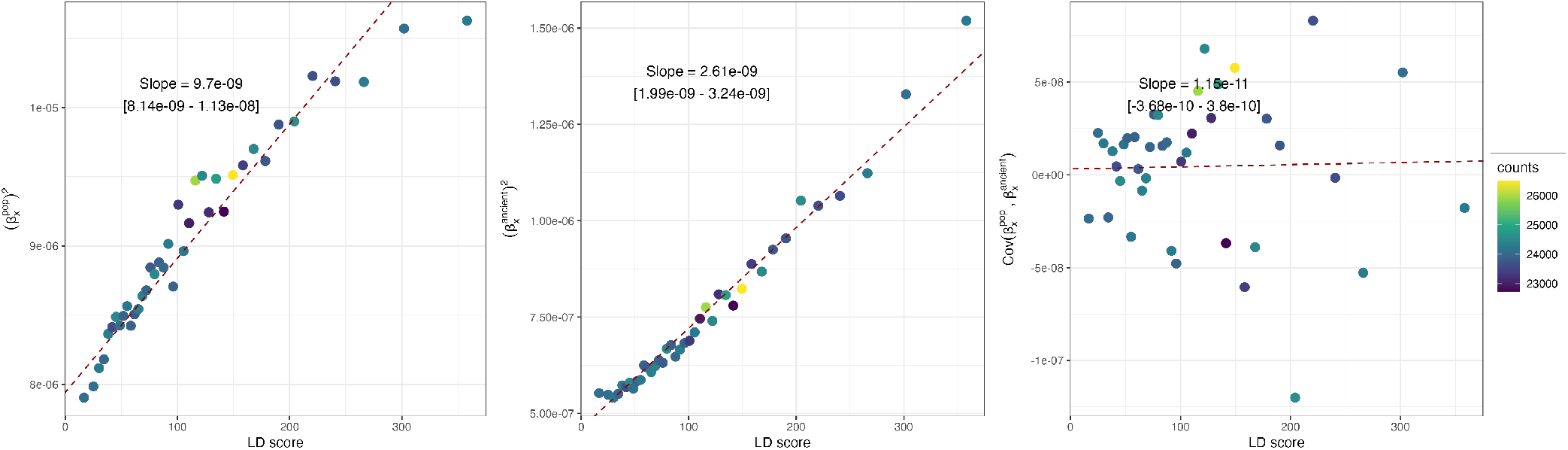
Univariate LD score regression on the effect sizes from the female UK Biobank cohort and on those inferred from temporal allele frequency gradients spanning the last 14, 000 years. **Left panel)**. LD score regression on the squared effect sizes from the population GWAS in the female cohort of the UK Biobank on the phenotype ‘number of live births’. **Central panel)**. LD score regression on the squared effect sizes inferred from genome-wide selection coefficients estimated from ancient allele frequency changes (Akbari *et al*., 2026). **Right panel)**. LD score regression on the product of UK Biobank-based effect sizes and ancient DNA-inferred effect sizes (the slope is not significantly different from zero). The dashed red line shows the slope obtained from the LD score regression through the underlying data, the slope is reported at the top of each panel with the corresponding 95% confidence interval obtained with the leave-one-chromosome-out approach. Across all panels, LD score values above 400 were removed since they represented clear outliers in our analysis.

### S6 LD score regression on published population GWAS effect sizes

**Figure S30:**
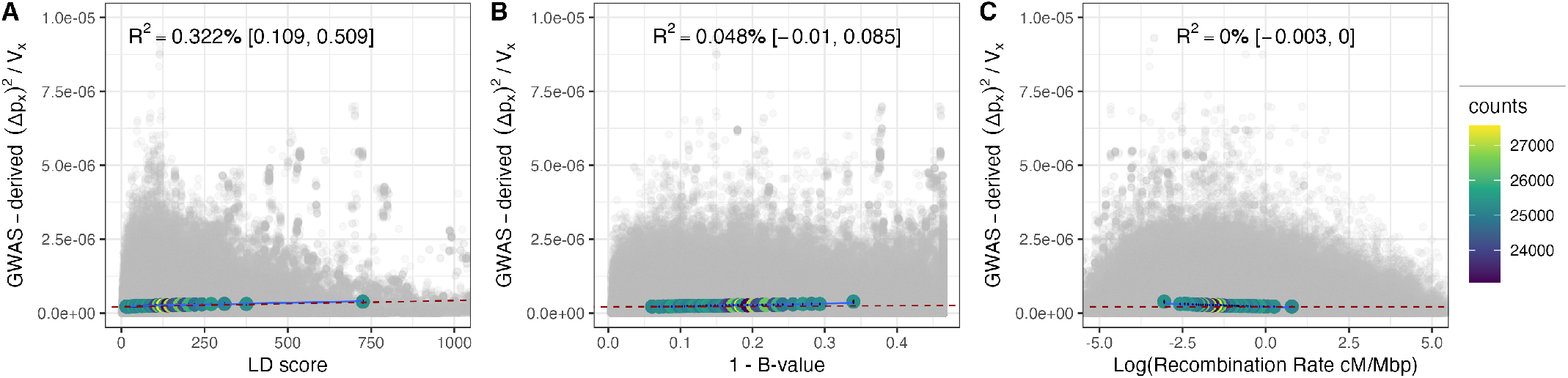
Decomposing the squared allele frequency change inferred from population GWAS effect sizes for common SNPs. Here the GWAS is in females on the phenotype ‘number of live births’ with PCA covariates from Howrigan *et al*. (2023). For details, see caption in Fig. S4. The summary statistics for this multiregression 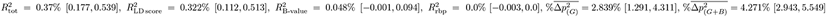.

**Figure S31:**
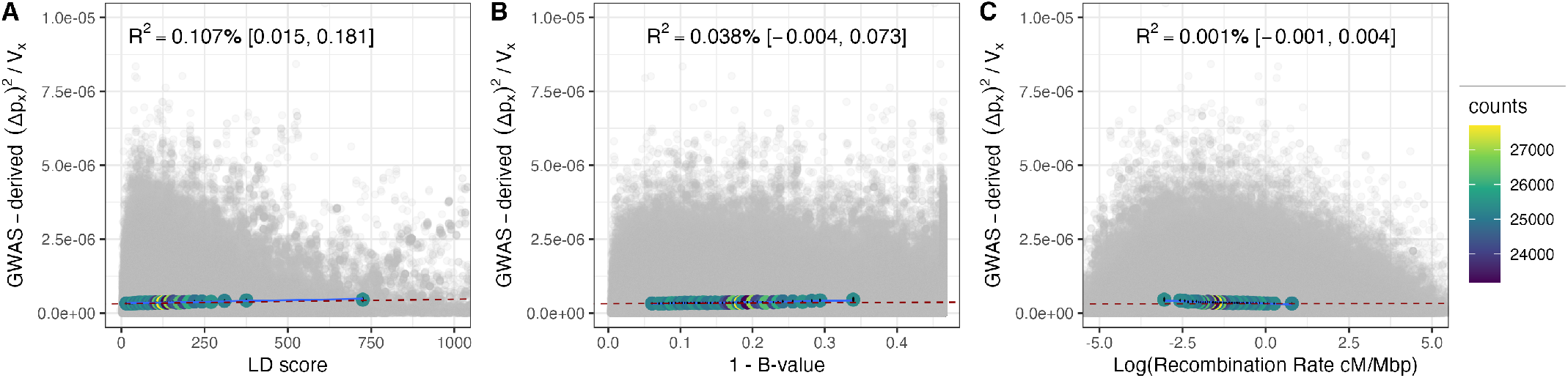
Decomposing the squared allele frequency change inferred from population GWAS effect sizes for common SNPs. Here the GWAS is in males on the phenotype ‘number of children fathered’ with PCs covariates from Howrigan *et al*. (2023). For details, see caption in Fig. S4. The summary statistics for this multiregression are 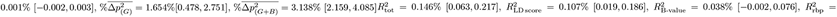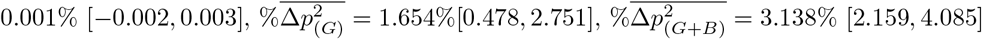.

## 4 Supplementary Tables

### S1 Number of live births - Female UK Biobank phenotype

#### S1.1 Non-partitioned LD-score

**Table S1:** R-squared and % of average squared allele frequency change based on the multivariate LD score regression on the population GWAS effect sizes in females on the phenotype ‘number of live births’ without PCs covariates. Values within brackets indicate the 95% confidence interval based on the leave-one-chromosome-out approach.

| | $R^2$ | $\% \Delta p_x^2$ |
| --- | --- | --- |
| $\ell_x$ | 0.294% [0.152, 0.426] | 4.72% [2.789, 6.533] |
| $B_x$ | 0.051% [0.005, 0.095] | 2.617% [1.489, 3.779] |
| $\log_e(r_{bp_x})$ | 0% [-0.002, 0] | -0.021% [-0.315, 0.292] |
| $\ell_x + B_x$ | 0.345% [0.205, 0.472] | 7.336% [5.537, 9.053] |
| $\ell_x + B_x + \log_e(r_{bp_x})$ | 0.345% [0.204, 0.471] | 7.316% [5.682, 8.886] |

**Table S2:** R-squared and % of average squared allele frequency change based on the multivariate LD score regression on the population GWAS effect sizes in females on the phenotype ‘number of live births’ with PCs covariates. Values within brackets indicate the 95% confidence interval based on the leave-one-chromosome-out approach.

| | $R^2$ | $\% \Delta p_x^2$ |
| --- | --- | --- |
| $\ell_x$ | 0.285% [0.147, 0.413] | 4.63% [2.743, 6.402] |
| $B_x$ | 0.05% [0.005, 0.092] | 2.566% [1.482, 3.685] |
| $\log_e(r_{bp_x})$ | 0% [-0.002, 0] | -0.007% [-0.294, 0.298] |
| $\ell_x + B_x$ | 0.334% [0.198, 0.458] | 7.196% [5.435, 8.877] |
| $\ell_x + B_x + \log_e(r_{bp_x})$ | 0.334% [0.198, 0.457] | 7.189% [5.581, 8.735] |

**Table S3:** Decomposition of the average squared allele frequency change into polygenic linked selection, environment and Mendelian noise components, based on population GWAS effect sizes estimated in the female UKB cohort for the phenotype ‘number of live births’, with and without PCs covariates. Values within brackets indicate the 95% confidence interval based on the leave-one-chromosome-out approach.

|  | Without PCs | With PCs |
| --- | --- | --- |
| Selection ($\ell_x + B_x$) | 3.90 [2.90, 4.85] $\times 10^{-8}$ | 3.82 [2.85, 4.74] $\times 10^{-8}$ |
| Environment | 2.16 [2.10, 2.23] $\times 10^{-7}$ | 2.16 [2.10, 2.22] $\times 10^{-7}$ |
| Mendelian | 2.77 [2.76, 2.78] $\times 10^{-7}$ | 2.77 [2.76, 2.78] $\times 10^{-7}$ |
| Total | 5.31 [5.25, 5.38] $\times 10^{-7}$ | 5.31 [5.24, 5.38] $\times 10^{-7}$ |

#### S1.2 Partitioned LD-score

**Table S4:** R-squared and % of average squared allele frequency change based on the MAF-stratified multivariate LD score regression on population GWAS effect sizes estimated in the female UKB cohort for the phenotype ‘number of live births’, without PCs covariates. Values within brackets indicate the 95% confidence interval based on the leave-one-chromosome-out approach.

| | $R^2$ | $\% \Delta p_x^2$ |
| --- | --- | --- |
| MAF-bin1 | 0% [-0.002, 0.001] | 0% [NA, NA] |
| MAF-bin2 | 0.003% [-0.005, 0.009] | 0% [NA, NA] |
| MAF-bin3 | 0.028% [0, 0.053] | 0.288% [-0.198, 0.769] |
| MAF-bin4 | 0.021% [-0.017, 0.052] | 0.066% [-0.593, 0.302] |
| MAF-bin5 | 0.05% [0.009, 0.088] | 0.203% [-0.559, 0.897] |
| MAF-bin6 | 0.189% [-0.092, 0.438] | 1.776% [-0.106, 4.211] |
| MAF-bin7 | 0.035% [-0.017, 0.073] | 0.392% [-0.316, 1.017] |
| MAF-bin8 | 0.047% [-0.018, 0.098] | 0.889% [-0.045, 1.774] |
| MAF-bin9 | 0.015% [-0.006, 0.031] | 0.162% [-0.446, 0.726] |
| MAF-bin10 | 0.036% [-0.009, 0.074] | 1.126% [0.318, 1.849] |
| $B_x$ | 0.06% [0.009, 0.102] | 2.788% [1.6, 3.864] |
| $\log_e(r_{bp_x})$ | 0% [-0.001, 0] | -0.04% [-0.304, 0.194] |
| all MAF-bins | 0.422% [0.153, 0.605] | 4.903% [3.617, 5.981] |
| all MAF-bins + $B_x$ | 0.482% [0.21, 0.658] | 7.69% [6.02, 9.043] |
| all MAF-bins + $B_x$ + $\log_e(r_{bp_x})$ | 0.482% [0.21, 0.657] | 7.65% [6.042, 8.911] |

**Table S5:** R-squared and % of average squared allele frequency change based on the MAF-stratified multivariate LD score regression on population GWAS effect sizes estimated in the female UKB cohort for the phenotype ‘number of live births’, with PCs covariates. Values within brackets indicate the 95% confidence interval based on the leave-one-chromosome-out approach.

| | $R^2$ | $\% \Delta p_x^2$ |
| --- | --- | --- |
| MAF-bin1 | 0% [-0.002, 0] | 0% [NA, NA] |
| MAF-bin2 | 0.003% [-0.005, 0.009] | 0% [NA, NA] |
| MAF-bin3 | 0.028% [-0.001, 0.054] | 0.299% [-0.191, 0.786] |
| MAF-bin4 | 0.021% [-0.017, 0.052] | 0.074% [-0.572, 0.35] |
| MAF-bin5 | 0.048% [0.009, 0.085] | 0.209% [-0.528, 0.883] |
| MAF-bin6 | 0.175% [-0.08, 0.404] | 1.692% [-0.082, 3.987] |
| MAF-bin7 | 0.035% [-0.016, 0.074] | 0.401% [-0.301, 1.026] |
| MAF-bin8 | 0.049% [-0.017, 0.1] | 0.925% [-0.001, 1.809] |
| MAF-bin9 | 0.013% [-0.007, 0.029] | 0.128% [-0.489, 0.702] |
| MAF-bin10 | 0.036% [-0.008, 0.072] | 1.117% [0.325, 1.825] |
| $B_x$ | 0.058% [0.009, 0.098] | 2.731% [1.584, 3.77] |
| $\log_e(r_{bp_x})$ | 0% [-0.001, 0] | -0.028% [-0.285, 0.2] |
| all MAF-bins | 0.408% [0.159, 0.575] | 4.845% [3.621, 5.908] |
| all MAF-bins + $B_x$ | 0.466% [0.214, 0.628] | 7.576% [5.959, 8.923] |
| all MAF-bins + $B_x$ + $\log_e(r_{bp_x})$ | 0.466% [0.213, 0.628] | 7.548% [5.983, 8.815] |

**Table S6:**
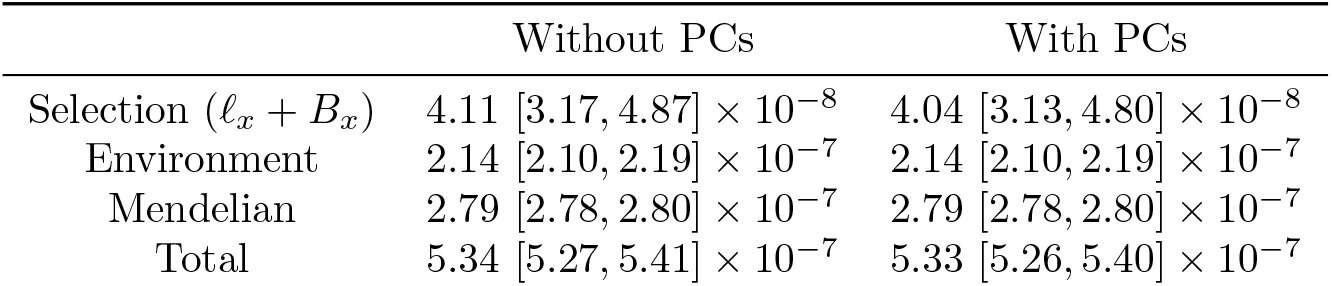
Decomposition of the average squared allele frequency change into the MAF-stratified polygenic linked selection, environment and Mendelian noise components, based on population GWAS effect sizes estimated in the female UKB cohort for the phenotype ‘number of live births’, with and without PCs covariates. Values within brackets indicate the 95% confidence interval based on the leave-one-chromosome-out approach.

### S2 Number of children fathered - Male UK Biobank phenotype

#### S2.1 Non-partitioned LD-score

**Table S7:** R-squared and % of average squared allele frequency change based on the multivariate LD score regression on the population GWAS effect sizes in males on the phenotype ‘number of children fathered’ without PCs covariates. Values within brackets indicate the 95% confidence interval based on the leave-one-chromosome-out approach.

| | $R^2$ | $\% \Delta p_x^2$ |
| --- | --- | --- |
| $\ell_x$ | 0.077% $[0.004, 0.14]$ | 2.519% $[0.918, 4.033]$ |
| $B_x$ | 0.019% $[-0.001, 0.037]$ | 1.764% $[0.835, 2.697]$ |
| $\log_e(r_{bp_x})$ | 0% $[-0.001, 0.001]$ | -0.101% $[-0.271, 0.079]$ |
| $\ell_x + B_x$ | 0.096% $[0.013, 0.167]$ | 4.283% $[2.317, 6.166]$ |
| $\ell_x + B_x + \log_e(r_{bp_x})$ | 0.096% $[0.013, 0.167]$ | 4.182% $[2.298, 5.993]$ |

**Table S8:** R-squared and % of average squared allele frequency change based on the multivariate LD score regression on the population GWAS effect sizes in males on the phenotype ‘number of children fathered’ with PCs covariates. Values within brackets indicate the 95% confidence interval based on the leave-one-chromosome-out approach.

| | $R^2$ | $\% \Delta p_x^2$ |
| --- | --- | --- |
| $\ell_x$ | 0.076% $[0.006, 0.138]$ | 2.511% $[0.944, 3.99]$ |
| $B_x$ | 0.019% $[-0.002, 0.037]$ | 1.757% $[0.822, 2.695]$ |
| $\log_e(r_{bp_x})$ | 0% $[-0.001, 0.001]$ | -0.098% $[-0.266, 0.08]$ |
| $\ell_x + B_x$ | 0.095% $[0.015, 0.165]$ | 4.268% $[2.347, 6.104]$ |
| $\ell_x + B_x + \log_e(r_{bp_x})$ | 0.095% $[0.014, 0.165]$ | 4.17% $[2.328, 5.936]$ |

**Table S9:**
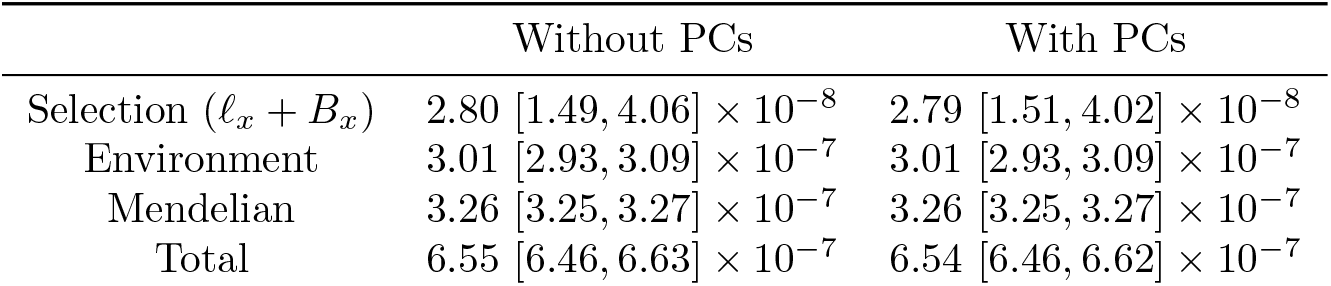
Decomposition of the average squared allele frequency change into polygenic linked selection, environment and Mendelian noise components, based on population GWAS effect sizes estimated in the male UKB cohort for the phenotype ‘number of live births’, with and without PCs covariates. Values within brackets indicate the 95% confidence interval based on the leave-one-chromosome-out approach.

#### S2.2 Partitioned LD-score

**Table S10:** R-squared and % of average squared allele frequency change based on the MAF-stratified multivariate LD score regression on population GWAS effect sizes estimated in the male UKB cohort for the phenotype ‘number of children fathered’, without PCs covariates. Values within brackets indicate the 95% confidence interval based on the leave-one-chromosome-out approach.

| | $R^2$ | $\% \Delta p_x^2$ |
| --- | --- | --- |
| MAF-bin1 | 0% $[-0.002, 0.002]$ | 0.005% $[-0.154, 0.013]$ |
| MAF-bin2 | 0% $[-0.002, 0]$ | 0% $[NA, NA]$ |
| MAF-bin3 | 0.007% $[-0.004, 0.016]$ | 0.175% $[-0.13, 0.488]$ |
| MAF-bin4 | 0.001% $[-0.003, 0.005]$ | 0% $[NA, NA]$ |
| MAF-bin5 | 0.018% $[-0.005, 0.037]$ | 0.32% $[-0.342, 0.965]$ |
| MAF-bin6 | 0.015% $[-0.011, 0.036]$ | 0.258% $[-0.204, 0.75]$ |
| MAF-bin7 | 0.035% $[0.004, 0.063]$ | 0.635% $[0.065, 1.159]$ |
| MAF-bin8 | 0.045% $[-0.046, 0.112]$ | 1.295% $[-0.553, 3.148]$ |
| MAF-bin9 | 0.002% $[-0.012, 0.011]$ | 0.233% $[-0.574, 1.003]$ |
| MAF-bin10 | 0% $[-0.005, 0.002]$ | 0.048% $[-0.588, 0.367]$ |
| $B_x$ | 0.021% $[-0.001, 0.041]$ | 1.829% $[0.881, 2.788]$ |
| $\log_e(r_{bp_x})$ | 0.001% $[-0.001, 0.002]$ | -0.117% $[-0.29, 0.069]$ |
| all MAF-bins | 0.125% $[-0.011, 0.208]$ | 2.968% $[1.313, 4.101]$ |
| all MAF-bins + $B_x$ | 0.146% $[0, 0.237]$ | 4.797% $[2.645, 6.437]$ |
| all MAF-bins + $B_x$ + $\log_e(r_{bp_x})$ | 0.146% $[0, 0.238]$ | 4.68% $[2.619, 6.243]$ |

**Table S11:** R-squared and % of average squared allele frequency change based on the MAF-stratified multivariate LD score regression on population GWAS effect sizes estimated in the male UKB cohort for the phenotype ‘number of children fathered’, with PCs covariates. Values within brackets indicate the 95% confidence interval based on the leave-one-chromosome-out approach.

| | $R^2$ | $\% \Delta p_x^2$ |
| --- | --- | --- |
| MAF-bin1 | 0% [-0.003, 0.002] | 0.01% [-0.139, 0.05] |
| MAF-bin2 | 0% [-0.002, 0] | 0% [NA, NA] |
| MAF-bin3 | 0.007% [-0.004, 0.016] | 0.176% [-0.125, 0.485] |
| MAF-bin4 | 0.002% [-0.003, 0.005] | 0% [NA, NA] |
| MAF-bin5 | 0.017% [-0.005, 0.036] | 0.304% [-0.347, 0.938] |
| MAF-bin6 | 0.015% [-0.01, 0.035] | 0.253% [-0.187, 0.723] |
| MAF-bin7 | 0.034% [0.004, 0.062] | 0.621% [0.057, 1.142] |
| MAF-bin8 | 0.046% [-0.046, 0.113] | 1.301% [-0.53, 3.137] |
| MAF-bin9 | 0.003% [-0.012, 0.011] | 0.239% [-0.557, 0.997] |
| MAF-bin10 | 0% [-0.005, 0.002] | 0.047% [-0.588, 0.374] |
| $B_x$ | 0.021% [-0.001, 0.041] | 1.822% [0.872, 2.782] |
| $\log_e(r_{bp_x})$ | 0.001% [-0.001, 0.002] | -0.114% [-0.284, 0.07] |
| all MAF-bins | 0.124% [-0.009, 0.206] | 2.952% [1.358, 4.071] |
| all MAF-bins + $B_x$ | 0.145% [0.002, 0.235] | 4.774% [2.688, 6.395] |
| all MAF-bins + $B_x$ + $\log_e(r_{bp_x})$ | 0.145% [0.001, 0.236] | 4.66% [2.663, 6.206] |

**Table S12:** Decomposition of the average squared allele frequency change into the MAF-stratified polygenic linked selection, environment and Mendelian noise components, based on population GWAS effect sizes estimated in the male UKB cohort for the phenotype ‘number of children fathered’, with and without PCs covariates. Values within brackets indicate the 95% confidence interval based on the leave-one-chromosome-out approach.

|  | Without PCs | With PCs |
| --- | --- | --- |
| Selection ($\ell_x + B_x$) | 3.16 [1.71, 4.26] $\times 10^{-8}$ | 3.14 [1.74, 4.23] $\times 10^{-8}$ |
| Environment | 2.97 [2.90, 3.07] $\times 10^{-7}$ | 2.97 [2.90, 3.07] $\times 10^{-7}$ |
| Mendelian | 3.29 [3.28, 3.30] $\times 10^{-7}$ | 3.29 [3.28, 3.30] $\times 10^{-7}$ |
| Total | 6.58 [6.50, 6.66] $\times 10^{-7}$ | 6.58 [6.50, 6.66] $\times 10^{-7}$ |

### S3 Combined Male and Female UK Biobank cohorts

#### S3.1 Weighted-Average Approach

**Table S13:**
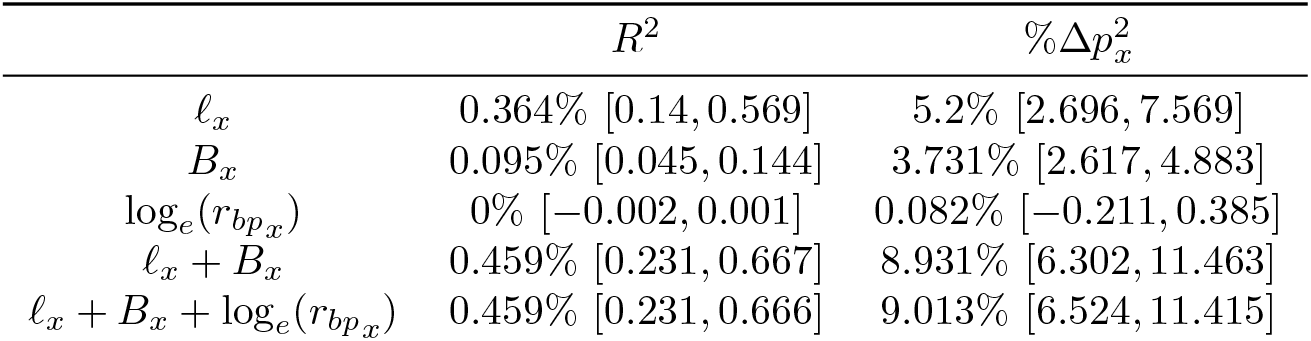
R-squared and % of average squared allele frequency change based on the multivariate LD score regression on the combined female and male UKB population GWAS effect sizes, without PCs covariates. Female and male cohorts were combined using the “weighted-average” approach. Values within brackets indicate the 95% confidence interval based on the leave-one-chromosome-out approach.

**Table S14:**
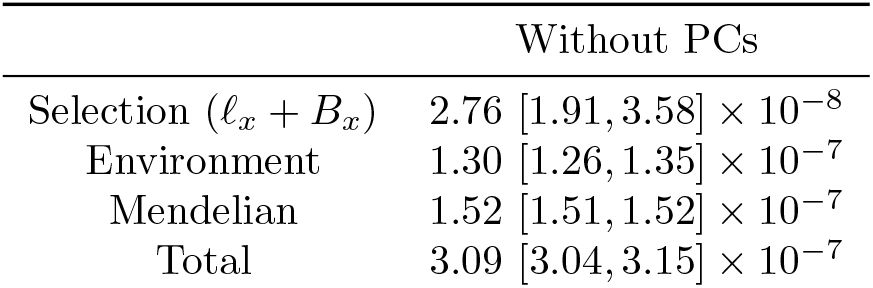
Decomposition of the average squared allele frequency change into polygenic linked selection, environment and Mendelian noise components, based on the “weighted-average” combined female and male UKB population GWAS effect sizes, without PCs covariates. Values within brackets indicate the 95% confidence interval based on the leave-one-chromosome-out approach.

|  | Without PCs |
| --- | --- |
| Selection ($\ell_x + B_x$) | $2.76 [1.91, 3.58] \times 10^{-8}$ |
| Environment | $1.30 [1.26, 1.35] \times 10^{-7}$ |
| Mendelian | $1.52 [1.51, 1.52] \times 10^{-7}$ |
| Total | $3.09 [3.04, 3.15] \times 10^{-7}$ |

#### S3.2 Fixed-Effect Approach

**Table S15:** R-squared and % of average squared allele frequency change based on the multivariate LD score regression on the combined female and male UKB population GWAS effect sizes, without PCs covariates. Female and male cohorts were combined using the “fixed-effect” approach. Values within brackets indicate the 95% confidence interval based on the leave-one-chromosome-out approach.

| | $R^2$ | $\% \Delta p_x^2$ |
| --- | --- | --- |
| $\ell_x$ | 0.386% [0.157, 0.595] | 5.384% [2.876, 7.755] |
| $B_x$ | 0.1% [0.047, 0.152] | 3.832% [2.705, 4.998] |
| $\log_e(r_{bp_x})$ | 0.001% [-0.003, 0.002] | 0.117% [-0.167, 0.412] |
| $\ell_x + B_x$ | 0.485% [0.253, 0.698] | 9.215% [6.604, 11.73] |
| $\ell_x + B_x + \log_e(r_{bp_x})$ | 0.486% [0.254, 0.697] | 9.332% [6.852, 11.727] |

**Table S16:**
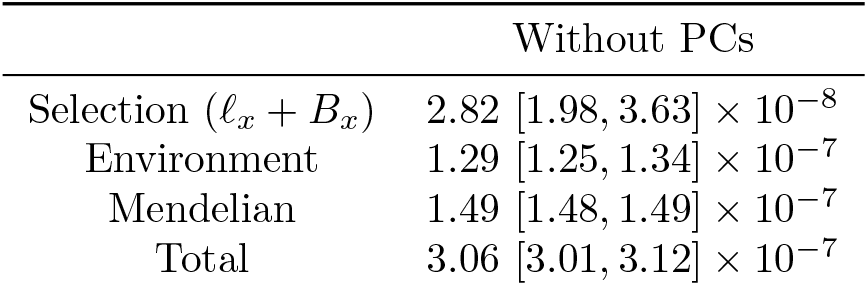
Decomposition of the average squared allele frequency change into polygenic linked selection, environment and Mendelian noise components, based on the “fixed-effect” combined female and male UKB population GWAS effect sizes, without PCs covariates. Values within brackets indicate the 95% confidence interval based on the leave-one-chromosome-out approach.

#### S3.3 Random-Effect Approach

**Table S17:** R-squared and % of average squared allele frequency change based on the multivariate LD score regression on the combined female and male UKB population GWAS effect sizes, without PCs covariates. Female and male cohorts were combined using the “random-effect” approach. Values within brackets indicate the 95% confidence interval based on the leave-one-chromosome-out approach.

| | $R^2$ | $\% \Delta p_x^2$ |
| --- | --- | --- |
| $\ell_x$ | 0.37% $[0.144, 0.577]$ | 4.161% $[2.163, 6.054]$ |
| $B_x$ | 0.097% $[0.048, 0.147]$ | 2.999% $[2.118, 3.91]$ |
| $\log_e(r_{bp_x})$ | 0% $[-0.003, 0.002]$ | 0.079% $[-0.154, 0.321]$ |
| $\ell_x + B_x$ | 0.467% $[0.237, 0.678]$ | 7.16% $[5.047, 9.198]$ |
| $\ell_x + B_x + \log_e(r_{bp_x})$ | 0.468% $[0.238, 0.677]$ | 7.239% $[5.243, 9.168]$ |

**Table S18:**
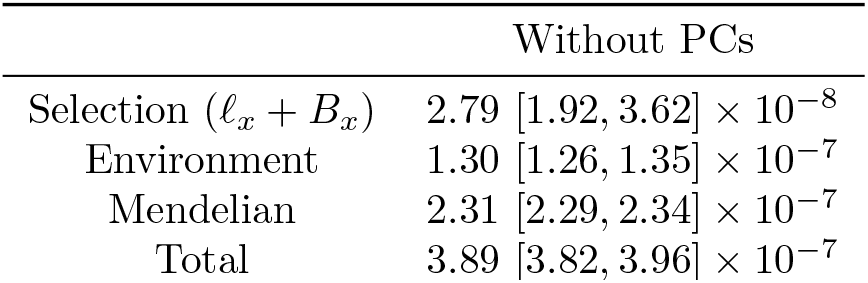
Decomposition of the average squared allele frequency change into polygenic linked selection, environment and Mendelian noise components, based on the “random-effect” combined female and male UKB population GWAS effect sizes, without PCs covariates. Values within brackets indicate the 95% confidence interval based on the leave-one-chromosome-out approach.

## Notes

### Competing Interest Statement

The authors have declared no competing interest.

